# Forecasting estuarine biodiversity and population abundance for conservation: spatial scale, forecast horizon, and choice of measure

**DOI:** 10.64898/2026.04.20.719659

**Authors:** Masami Fujiwara

## Abstract

Conservation decisions increasingly rely on forecasts, but how far ahead biodiversity and population abundance can be forecast, and at which spatial scale, is rarely known. I used monthly monitoring data on fishes and invertebrates in seven Texas estuaries to forecast biodiversity (Hill numbers, which weight rare and common species differently) and the abundance of eight common taxa, 1 to 24 months ahead, for each estuary and the coast as a whole. Each series was divided into a seasonal cycle and departures from it. Forecasts were evaluated by accuracy and by skill, the improvement in accuracy over two references: the seasonal cycle plus the average departure of all earlier months (cumulative average) or of the last 12 months (12-month average). The seasonal cycle, which repeats each year, made up much of the month-to-month variation, and approximately two-thirds of single-estuary forecasts were within a set tolerance one month ahead. Beyond the seasonal cycle, forecasts were better than the cumulative average mainly one or two months ahead, and water temperature and salinity did not improve them. Pooling the estuaries made forecasts more accurate at every lead time, but improved skill only a few months ahead. Biodiversity was forecast more accurately than abundance, but without improved skill. After shoreline biodiversity increased in 2020 and stayed high, the high skill against the cumulative average disappeared against the 12-month average. I suggest that conservation planning use the seasonal expectation adjusted to the recent level, with a buffer, and that forecast skill be reported against both references.

## Introduction

Ecological forecasting has been advocated as a way to improve planning and decision-making for changing ecosystems (Clark et al. 2001), including ecosystems affected by a changing climate. Ecological prediction has since expanded rapidly (Mouquet et al. 2015), and near-term forecasts are now used in some management settings (Dietze et al. 2018) although such applications remain few. However, predictability, the ability to forecast, differs systematically with how far ahead a forecast looks, with the spatial scale at which biodiversity is observed, and with what is measured (Beckage et al. 2011; Ward et al. 2014; Petchey et al. 2015). These differences matter for conservation because decisions are made for particular time frames and spatial scales, and the question is whether biodiversity or the abundance of a population can be forecast well enough for the decision at hand. Few studies have evaluated this across several time frames and spatial scales together.

Conservation decisions need forecasts at different lead times (the number of months between the forecast date, when a forecast is made, and the month it forecasts; Payne et al. 2017; Dietze et al. 2018), but predictability changes with lead time. Responding to a flood or an oil spill calls for forecasts weeks to months ahead, whereas designating a protected area requires forecasts years ahead. Forecast error is often expected to grow with lead time as unpredictable variation accumulates (Petchey et al. 2015; Dietze 2017): a forecast one month ahead can use the recent level of a series, whereas a forecast 12 months ahead must rely on slower change. In contrast, the seasonal cycle repeats each year and can be forecast at any lead time, and it affects biodiversity as well as abundance because some species are rare only in some seasons (Hewitt et al. 2016). Therefore, the seasonal cycle must be separated from the variation to be forecast. The forecast horizon summarizes predictability as the longest lead time up to which forecasts stay accurate enough or better than a reference forecast (Petchey et al. 2015). For example, forecasts accurate enough one to three months ahead but not four have a three-month horizon.

Conservation actions are often taken at single sites, whereas systematic conservation planning and ecosystem-based management coordinate actions across regions (Margules and Pressey 2000; Leslie and McLeod 2007). Monitoring data can be summarized at either scale. Pooling observations over a region averages out local variation and sampling error while keeping the variation shared among places (Levin 1992; Liebhold et al. 2004; Wang and Loreau 2014). However, when different processes drive local dynamics, pooling combines unrelated signals (Wang and Loreau 2014), and local forecasts may be as good as regional ones. Therefore, spatial scale and lead time should be evaluated together.

Predictability may also differ between population abundance and biodiversity indices that combine many species. The population is the unit for which extinction risk is usually assessed (Lande 1993; Fujiwara 2007). Aggregate properties of a community vary less than its populations because the fluctuations of different species partly cancel (Doak et al. 1998; Tilman 1999). Therefore, biodiversity indices may be forecast more accurately than abundance. However, lower variability does not by itself make changes easier to predict. The order q of Hill numbers sets the weight given to rare and common species (Hill 1973; Jost 2006; Chao et al. 2014). For example, q = 0 (species richness) counts all species equally, whereas q = 1 (the exponential of Shannon entropy) and q = 2 (the inverse Simpson index) give more weight to common species.

A forecast can be evaluated in terms of its accuracy and its skill. Accuracy is how close forecasts are to the values observed later, and it can be measured as the root mean squared error (RMSE) of forecasts made without knowledge of those values (Hyndman and Koehler 2006). A forecast is accurate enough when its error stays within a tolerance, the largest miss acceptable for a decision. Skill is how much better a forecast is than a reference forecast; it is 0 when the forecast is no better and positive when it is better (Murphy 1988; Hyndman and Athanasopoulos 2021). Therefore, skill increases when the forecast improves and decreases when the reference improves. A natural reference is the seasonal cycle plus the average departure from it over all earlier months (the cumulative average), which a manager could compute from past data alone. A second reference, the seasonal cycle plus the average departure over the last 12 months (the 12-month average), follows recent levels and separates the skill of a model from knowing the recent level (Wesselkamp et al. 2025).

Crossing accuracy and skill gives four management situations: forecast-led, forecast with buffer, mean-led, and precaution. Forecasts that are within the tolerance and better than the cumulative average can lead decisions (forecast-led). Forecasts that are better but outside the tolerance can be used with a buffer, an allowance for forecast error such as a prediction interval, and revised as observations accumulate (forecast with buffer; Williams 2011). Forecasts that are within the tolerance but no better than the cumulative average add nothing to it, so decisions can rely on the cumulative average (mean-led). Forecasts that are neither within the tolerance nor better than the cumulative average call for a precautionary approach (precaution; Cooney 2004). Because accuracy and skill vary continuously, these situations summarize a gradient, and the lines between them are set by management.

Here, I used monthly data from a standardized monitoring program in seven estuaries on the Texas coast from 1982 to 2024, which sampled shorelines with bag seines and open-bay water with bay trawls (Martinez-Andrade 2018; Fujiwara et al. 2019). Biodiversity (Hill numbers of order q = 0, 1, and 2) and the abundance of eight common taxa were forecast for each estuary and for the coast as a whole with time-series models, with and without water temperature and salinity. Each forecast used only data from earlier months and looked 1 to 24 months ahead. I asked four questions: (1) how much of the month-to-month variation can be forecast beyond the seasonal cycle; (2) how accuracy and skill differ between single estuaries and the coast, and with lead time; (3) whether predictability differs between biodiversity and abundance and between the two sampling methods; and (4) where the series fall among the four management situations.

## Methods

Online Resource 1 gives additional methodological detail.

### Study system and data

The data were collected in seven Texas estuaries, from Galveston Bay in the north to Lower Laguna Madre in the south, by the Texas Parks and Wildlife Department (TPWD) Coastal Fisheries Marine Resource Monitoring Program, a standardized fisheries-independent monitoring program (Martinez-Andrade 2018; Fujiwara et al. 2019). Each month from 1982 to 2024, multiple sites were drawn at random in each estuary and sampled with two sampling methods, following stratified cluster sampling within each estuary and month (all organisms caught in a haul/tow form one cluster). On average, 18 bag seine hauls were made in each estuary each month, and 21 bay trawl tows in five estuaries and 11 in the two Laguna Madre estuaries (Table S1, Online Resource 2). Bag seines (18.3-m-long nets pulled along the shoreline) sample shallow shoreline habitats, and bay trawls (6.1-m-wide otter trawls towed for 10 min) sample deeper open-bay water. Because the two methods sample different parts of the community, they were analyzed separately throughout. No samples were taken in April and May 2020 (COVID-19 restrictions), and bay trawl sampling started in May 1982 in four estuaries (Table S2).

Before analysis, I removed taxa that were not counted as individuals (colonial invertebrates and some gelatinous invertebrates), not recorded consistently over time (sea squirts, possibly not recorded in the early years), or not targeted by the survey (reptiles, amphibians, and birds). These taxa made up 10.0% of the bay trawl catch by number (9.3% sea squirts) and 0.2% of the bag seine catch. The remaining catch included 494 taxa in bag seine samples and 538 in bay trawl samples (Table S3). Fishes made up 65% and 67% of the catch by number, respectively, and the rest were invertebrates such as shrimps, crabs, squids, and jellyfish.

Water temperature and salinity were measured with each sample. In bag seine samples, the coastwide monthly mean temperature had a median of 24.9 °C (10th to 90th percentile, 16.1 to 30.9 °C), and the monthly mean salinity had a median of 25.6 (18.0 to 31.9); bay trawl samples gave similar values (Online Resource 1). Median salinity ranged from approximately 17 in Galveston Bay to approximately 39 in Upper Laguna Madre. Most of the largest departures from the usual values for the calendar month coincided with floods in 1991 to 1992, 1998, 2002, 2007, and 2015, a drought in 2011, and a deep freeze in December 1989 (sizes and sources in Online Resource 1).

### Forecast targets

The first class of target was community biodiversity, measured with Hill numbers of order q = 0, 1, and 2 (Hill 1973; Jost 2006), which express biodiversity as an effective number of species. They were estimated from the pooled catch of all hauls/tows in each estuary and month, and for the coast as a whole in each month when all seven estuaries were sampled. The number of hauls/tows, and so the number of individuals caught, differed among months. Therefore, every value was estimated at a common sample coverage of 0.90 (Chao and Jost 2012; Chao et al. 2014) with the iNEXT package (Hsieh et al. 2016).

The second class was the abundance of eight common taxa: bay anchovy (*Anchoa mitchilli*), spot (*Leiostomus xanthurus*), Atlantic croaker (*Micropogonias undulatus*), pinfish (*Lagodon rhomboides*), menhaden (*Brevoortia* spp.), brown shrimp (*Farfantepenaeus aztecus*), white shrimp (*Litopenaeus setiferus*), and blue crab (*Callinectes sapidus*). Abundance was measured as the mean catch per haul/tow (catch per unit effort, CPUE) and analyzed as log(CPUE + 1).

Each target was analyzed for each sampling method, in each estuary and coastwide. This gave 48 biodiversity series (7 estuaries and the coast, 2 sampling methods, and 3 orders) and 128 abundance series (8 taxa in place of the 3 orders). April and May 2020 were filled only for fitting the models and were never used to evaluate the forecasts (Online Resource 1).

### Forecasting design

The analysis is outlined in Fig. 1. Most series vary with the seasons. Therefore, the forecasts were made for the departures from a seasonal curve, a harmonic regression with periods of 12, 6, and 4 months (Shumway and Stoffer 2017; Fig. S1, Online Resource 3; Table S4). The curve was fitted at each forecast date to the months up to that date only, so no later observation entered a forecast.

**Fig. 1.**
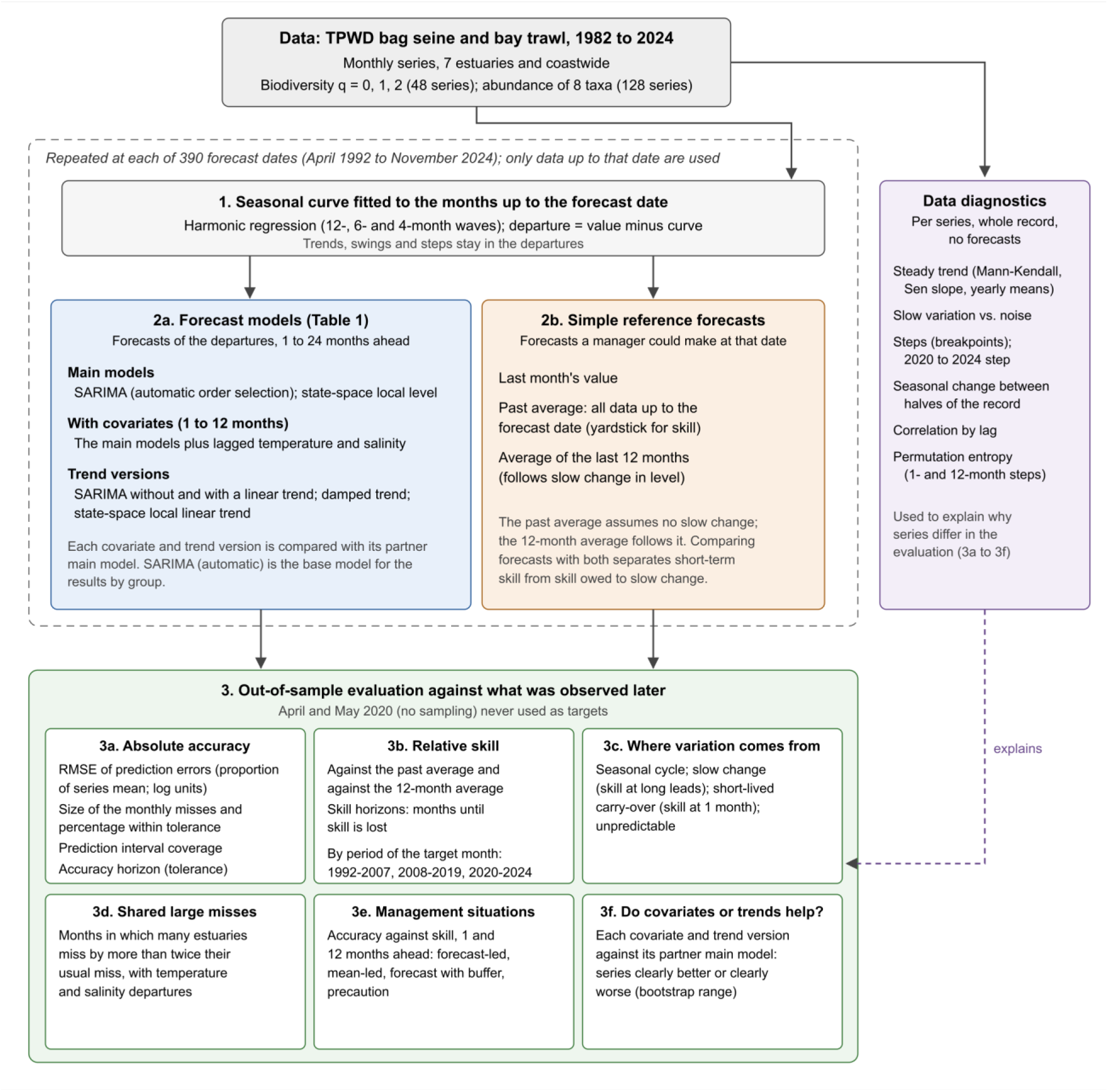
Overview of the analysis of the Texas Parks and Wildlife Department (TPWD) bag seine and bay trawl monitoring data. At each monthly forecast date, the seasonal curve was refitted to the months up to that date, the forecast models and reference forecasts were applied to the departures from it, and the forecasts were compared with the values observed later. The characteristics of the time series data, calculated from the whole record without forecasts, help explain why series differ. The models are listed in Table 1

**Table 1.**
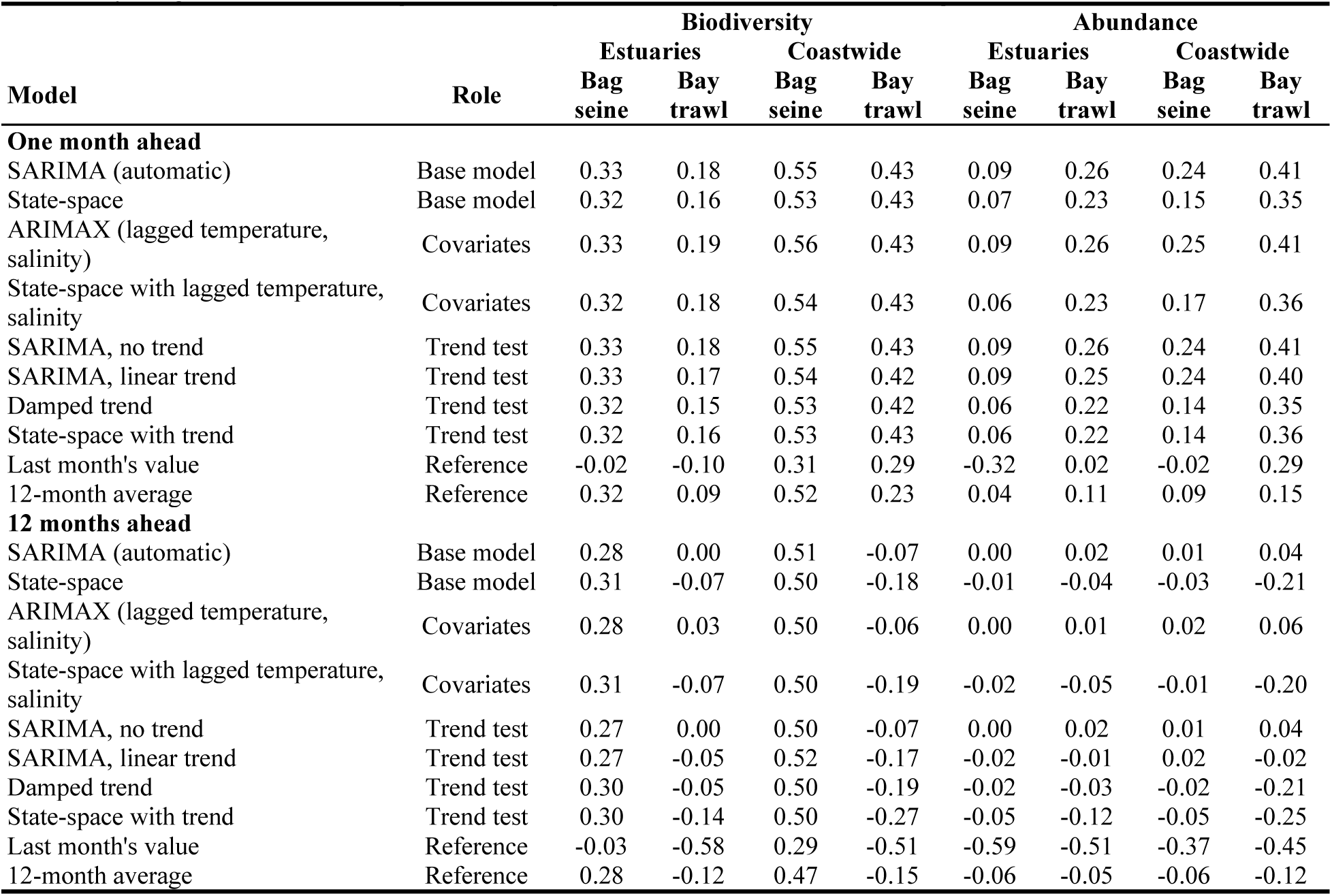
Skill of each forecasting model and reference against the cumulative average, one and 12 months ahead (median over the series of each group; 0 = no better than the cumulative average, negative = worse; Methods: Forecast accuracy and skill). Number of series per column: biodiversity 21 in estuaries and 3 coastwide; abundance 56 in estuaries and 8 coastwide. The models are described in Online Resource 1 (Sect. 1.9); full results with uncertainty ranges are in Tables S7 and S8 (Online Resource 2).

| Model | Role | **Biodiversity** |  |  |  | **Abundance** |  |  |  |
|---|---|---|---|---|---|---|---|---|---|
|  |  | **Estuaries** |  | **Coastwide** |  | **Estuaries** |  | **Coastwide** |  |
| **Model** | **Role** | **Bag seine** | **Bay trawl** | **Bag seine** | **Bay trawl** | **Bag seine** | **Bay trawl** | **Bag seine** | **Bay trawl** |
| **One month ahead** |  |  |  |  |  |  |  |  |  |
| SARIMA (automatic) | Base model | 0.33 | 0.18 | 0.55 | 0.43 | 0.09 | 0.26 | 0.24 | 0.41 |
| State-space | Base model | 0.32 | 0.16 | 0.53 | 0.43 | 0.07 | 0.23 | 0.15 | 0.35 |
| ARIMAX (lagged temperature, salinity) | Covariates | 0.33 | 0.19 | 0.56 | 0.43 | 0.09 | 0.26 | 0.25 | 0.41 |
| State-space with lagged temperature, salinity | Covariates | 0.32 | 0.18 | 0.54 | 0.43 | 0.06 | 0.23 | 0.17 | 0.36 |
| SARIMA, no trend | Trend test | 0.33 | 0.18 | 0.55 | 0.43 | 0.09 | 0.26 | 0.24 | 0.41 |
| SARIMA, linear trend | Trend test | 0.33 | 0.17 | 0.54 | 0.42 | 0.09 | 0.25 | 0.24 | 0.40 |
| Damped trend | Trend test | 0.32 | 0.15 | 0.53 | 0.42 | 0.06 | 0.22 | 0.14 | 0.35 |
| State-space with trend | Trend test | 0.32 | 0.16 | 0.53 | 0.43 | 0.06 | 0.22 | 0.14 | 0.36 |
| Last month's value | Reference | -0.02 | -0.10 | 0.31 | 0.29 | -0.32 | 0.02 | -0.02 | 0.29 |
| 12-month average | Reference | 0.32 | 0.09 | 0.52 | 0.23 | 0.04 | 0.11 | 0.09 | 0.15 |
| **12 months ahead** |  |  |  |  |  |  |  |  |  |
| SARIMA (automatic) | Base model | 0.28 | 0.00 | 0.51 | -0.07 | 0.00 | 0.02 | 0.01 | 0.04 |
| State-space | Base model | 0.31 | -0.07 | 0.50 | -0.18 | -0.01 | -0.04 | -0.03 | -0.21 |
| ARIMAX (lagged temperature, salinity) | Covariates | 0.28 | 0.03 | 0.50 | -0.06 | 0.00 | 0.01 | 0.02 | 0.06 |
| State-space with lagged temperature, salinity | Covariates | 0.31 | -0.07 | 0.50 | -0.19 | -0.02 | -0.05 | -0.01 | -0.20 |
| SARIMA, no trend | Trend test | 0.27 | 0.00 | 0.50 | -0.07 | 0.00 | 0.02 | 0.01 | 0.04 |
| SARIMA, linear trend | Trend test | 0.27 | -0.05 | 0.52 | -0.17 | -0.02 | -0.01 | 0.02 | -0.02 |
| Damped trend | Trend test | 0.30 | -0.05 | 0.50 | -0.19 | -0.02 | -0.03 | -0.02 | -0.21 |
| State-space with trend | Trend test | 0.30 | -0.14 | 0.50 | -0.27 | -0.05 | -0.12 | -0.05 | -0.25 |
| Last month's value | Reference | -0.03 | -0.58 | 0.29 | -0.51 | -0.59 | -0.51 | -0.37 | -0.45 |
| 12-month average | Reference | 0.28 | -0.12 | 0.47 | -0.15 | -0.06 | -0.05 | -0.06 | -0.12 |

Forecasts were made at 390 monthly forecast dates, from April 1992 to November 2024, each using all earlier months (at least 10 years of data). At each date, every model forecast 1 to 24 months ahead (1 to 12 months for the models with water temperature and salinity). The forecasts were compared with values observed later, which the models had not seen (out-of-sample evaluation). Accuracy and skill were calculated on the 367 forecast dates up to December 2022, so that every lead time was evaluated on the same dates; the comparison among periods uses all 390.

### Forecast models and references

Forecasts were made with a seasonal autoregressive integrated moving average model (SARIMA) with automatic order selection (Hyndman and Khandakar 2008; Box et al. 2015). Forecasts were also made with a state-space model, which separates a slowly changing level from observation noise (Durbin and Koopman 2012; Helske 2017). Both were also fitted with water temperature and salinity as covariates (ARIMAX and a state-space model with covariates). The covariates were values observed before the forecast date, with lags of 1 and 12 months one month ahead and a lag of 12 months from 2 to 12 months ahead. Therefore, these are true forecasts that do not depend on future environmental conditions. Trend versions were also fitted (Table 1; Online Resource 1).

Each model was compared with three references: the cumulative average and the 12-month average (the seasonal curve plus the mean departure of all earlier months or of the last 12 months), and last month’s value, which carries the latest departure forward. Each model also gave 80% and 95% prediction intervals.

### Forecast accuracy and skill

Accuracy was measured as the root mean squared error (RMSE) of the out-of-sample prediction errors. For biodiversity, the RMSE was divided by the series mean so that series of different sizes could be compared. Abundance was already on a log scale, so its RMSE was left in units of log(CPUE + 1) (hereafter log units). To judge whether a forecast was accurate enough to use, I set a tolerance of 30% of the series mean for biodiversity and 0.5 log units for abundance and recorded the percentage of monthly forecasts within it. This tolerance is approximately twice the sampling uncertainty of a typical value for one estuary and month (Online Resource 1; other tolerances in Fig. S2). The RMSE divided by the mean also depends on how variable a series is, so the RMSE divided by the standard deviation of the departures is given in Table S5 (Online Resource 2).

Skill was calculated as 1 minus the ratio of the mean squared prediction error of the forecast to that of the reference: 0 means no better than the reference, positive values mean better, and negative values mean worse. Skill was measured against both the cumulative average and the 12-month average. The 95% uncertainty ranges of the RMSE and the skill were obtained with a moving-block bootstrap over forecast dates (blocks of 12 months, 2,000 resamples; Künsch 1989). These ranges were used in two summaries. The forecast horizon was the number of consecutive months ahead, starting from one month, for which the whole range stayed within the tolerance (accuracy horizon) or above zero skill (skill horizon). Each series was also placed in one of the four management situations at 1, 12, and 24 months ahead, according to whether the whole range of its RMSE was within the tolerance and whether that of its skill against the cumulative average was above zero.

The month-to-month variation of each series was divided into four parts with the SARIMA forecasts and their skill against the cumulative average (Fig. 2): the seasonal cycle (the variation explained by the seasonal curve), slow change (variation that lasts for years, measured as the skill that persists 12 to 24 months ahead), short-lived carry-over (the resemblance of next month to this month, measured as the additional skill one month ahead), and the remainder (the unpredictable part). The remainder was what the SARIMA forecasts missed even one month ahead. To find months in which many estuaries missed together, a month counted as a large miss for a series when its one-month forecast missed by more than twice the series’ RMSE one month ahead.

**Fig. 2.**
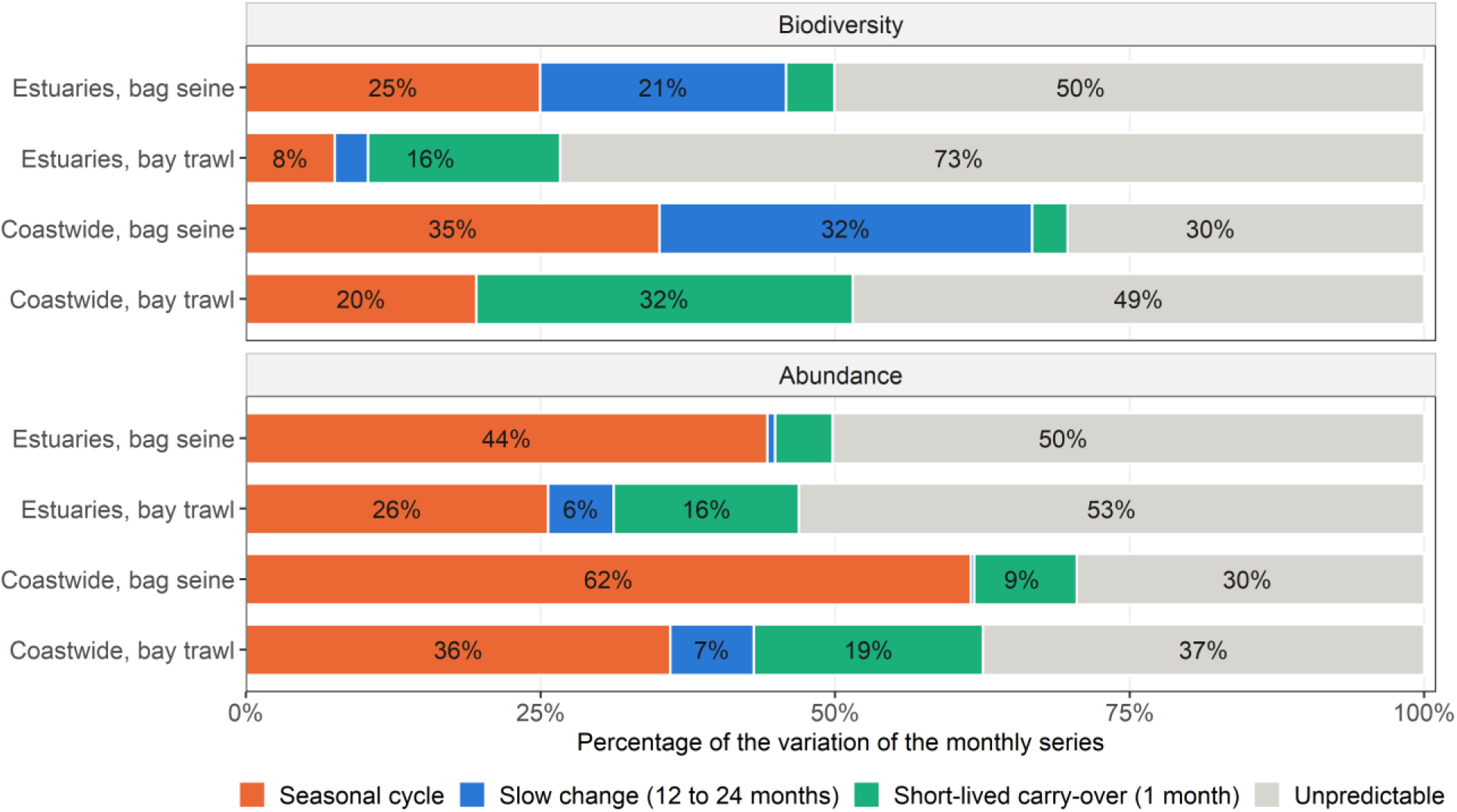
Division of the month-to-month variation of the monthly series into four parts, by target class, scale, and sampling method: the seasonal cycle, slow change (averaged over 12 to 24 months ahead and set to zero when negative), short-lived carry-over, and the remainder. Percentages are means over the series of each group; the four parts add to 100%

### Characteristics of the time series data

Three characteristics of the data were described without forecasting: slow change, order, and trends in water temperature and salinity. Each used the departures from a seasonal curve fitted once to the whole record of each series. Here, slow change was measured directly in the data, with the correlation between departures 1 to 36 months apart and with the proportion of their variance in a 25-month running mean (Fig. S3; Online Resource 1). This proportion was compared with that in simulated first-order autoregressive series with the same lag-1 correlation (the correlation between consecutive months). The order in each series was measured with weighted permutation entropy (Bandt and Pompe 2002; Fadlallah et al. 2013; Pennekamp et al. 2019). This describes how ordered a series is but does not by itself show whether its dynamics are random or nonlinear (Şen et al. 2024; Fig. S4).

Trends in water temperature and salinity were tested for each estuary and sampling method, and coastwide, with the Mann-Kendall test and the Sen slope (Mann 1945; Kendall 1970; Sen 1968). The tests used the yearly means of the departures from the seasonal curve (years with at least six sampled months), and p-values for the seven estuaries were adjusted for multiple tests (Benjamini and Hochberg 1995). The same test was applied to the yearly means of the departures of each forecast target.

### Software and use of generative AI

All analyses were done in R 4.6.1 (R Core Team 2026) with the packages forecast 9.0.2, KFAS 1.6.0, and iNEXT 3.0.2. Generative AI tools (ChatGPT, GPT-5, OpenAI; Claude Opus 5.5, Anthropic) assisted with R coding (interactive drafting, debugging, parallelization, annotation, and package suggestions) and with manuscript preparation (word and sentence choice, peer-review-style review, interactive revision, proofreading, and cross-references); the author reviewed all output and takes full responsibility for the content.

## Results

### Predictable variation and accuracy

A large part of the month-to-month variation was predictable (Fig. 2). In the estuaries, the seasonal cycle, slow change, and short-lived carry-over together accounted for approximately half of the variation in three of the four groups (47% to 50%) but only 27% in bay trawl biodiversity. Coastwide, they accounted for 51% to 70% (means across series of the SARIMA forecasts, with SDs of 7% to 23%; n = 21 or 56 series per group in the estuaries and 3 or 8 coastwide; Table S6). The other models gave nearly the same skill (Table 1). The seasonal cycle was the largest part for bag seine series in the estuaries (biodiversity 25% ± 9%; abundance 44% ± 25%). Slow change was large only for bag seine biodiversity, where it was present in every series (21% ± 6% in the estuaries; 32% ± 4% coastwide). In the other groups, it averaged 0% to 7%, and most series had none (Fig. 2). Slow change was also present in the data, beyond what the lag-1 correlation alone would give, in all biodiversity series and most abundance series (Fig. S3). Short-lived carry-over had group means of 16% to 32% for bay trawl series and 3% to 9% for bag seine series.

Forecast error grew little with lead time (Fig. 3; Table S6). In the estuaries, the median RMSE across series one month ahead was 32% to 33% of the series mean for biodiversity (n = 21 series per sampling method) and 0.65 to 0.66 log units for abundance (n = 56), and at 24 months it was 36% to 38% and 0.72 to 0.75 log units. Errors were smaller coastwide (19% to 23% of the series mean and 0.45 to 0.46 log units one month ahead; n = 3 and 8 series per sampling method). In the median estuary series, approximately two-thirds of the monthly forecasts were within the tolerance one month ahead (62% to 69%), and more than half at 24 months (53% to 62%).

**Fig. 3.**
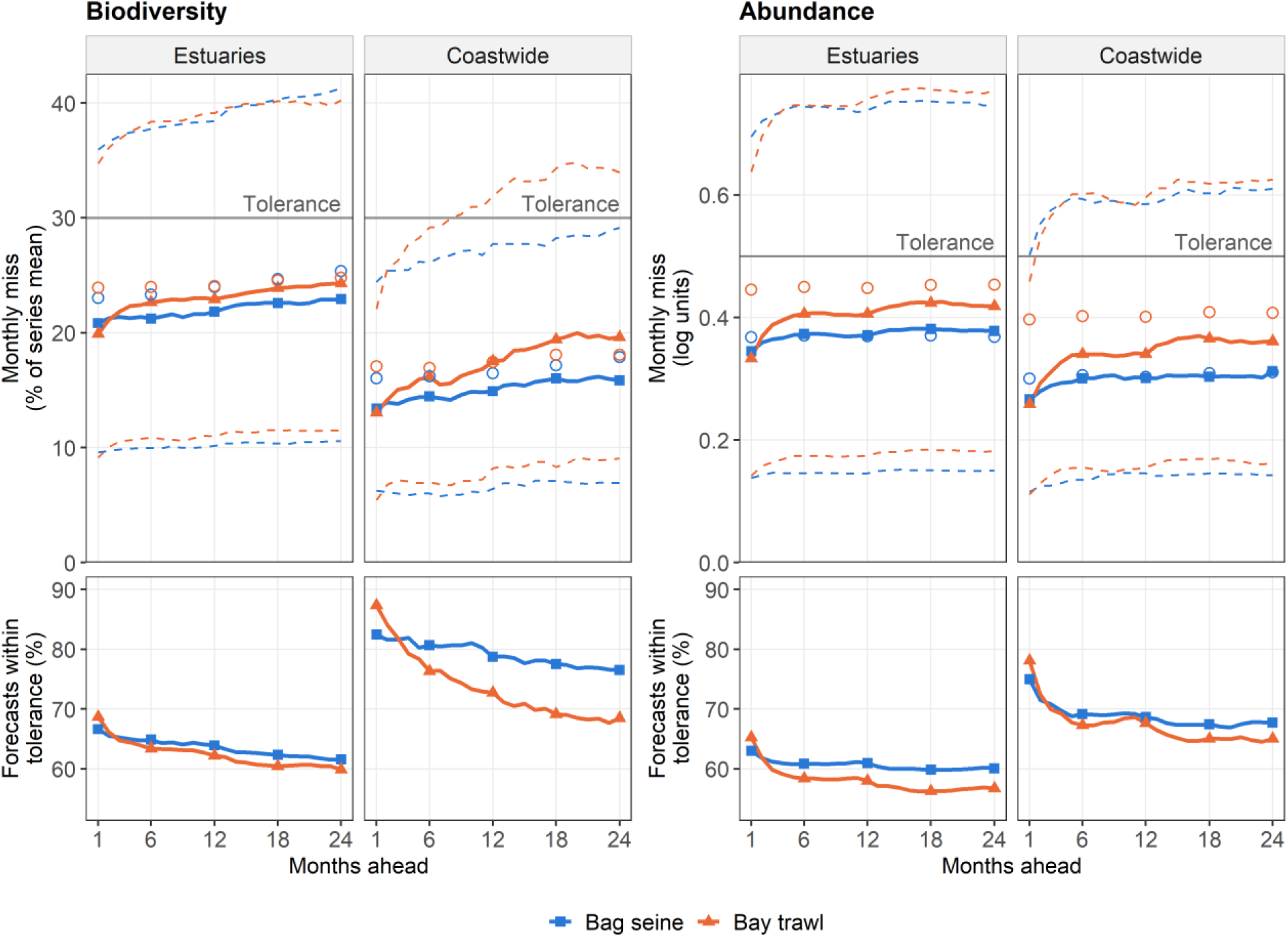
Size of the monthly forecast misses and percentage of forecasts within the tolerance, by months ahead. Top: absolute out-of-sample prediction errors of SARIMA forecasts, pooled over the series of each group and the forecast dates (per sampling method, 21 biodiversity and 56 abundance series in the estuaries; 3 and 8 coastwide); solid line: median (squares: bag seine; triangles: bay trawl); dashed lines: middle 50%. Biodiversity misses are a percentage of the series mean; abundance misses are in log units. Open circles: median miss of the cumulative average. Gray line: tolerance. Bottom: percentage of monthly forecasts within the tolerance. The cumulative average and the tolerance are defined in the Methods (Forecast models and references; Forecast accuracy and skill)

### Skill against the two references

Forecasts were better than the cumulative average one month ahead in every group (median skill across series 0.09 to 0.33 in the estuaries and 0.24 to 0.55 coastwide; Fig. 4; Table S6). This skill declined within a few months, and by 12 months ahead it was close to zero (−0.07 to 0.04) in every group except bag seine biodiversity (0.28 in the estuaries). Across series, the one-month skill against the cumulative average increased with the lag-1 correlation of the departures (Spearman’s ρ = 0.94 for the 112 estuary abundance series and 0.90 for the 42 estuary biodiversity series).

**Fig. 4.**
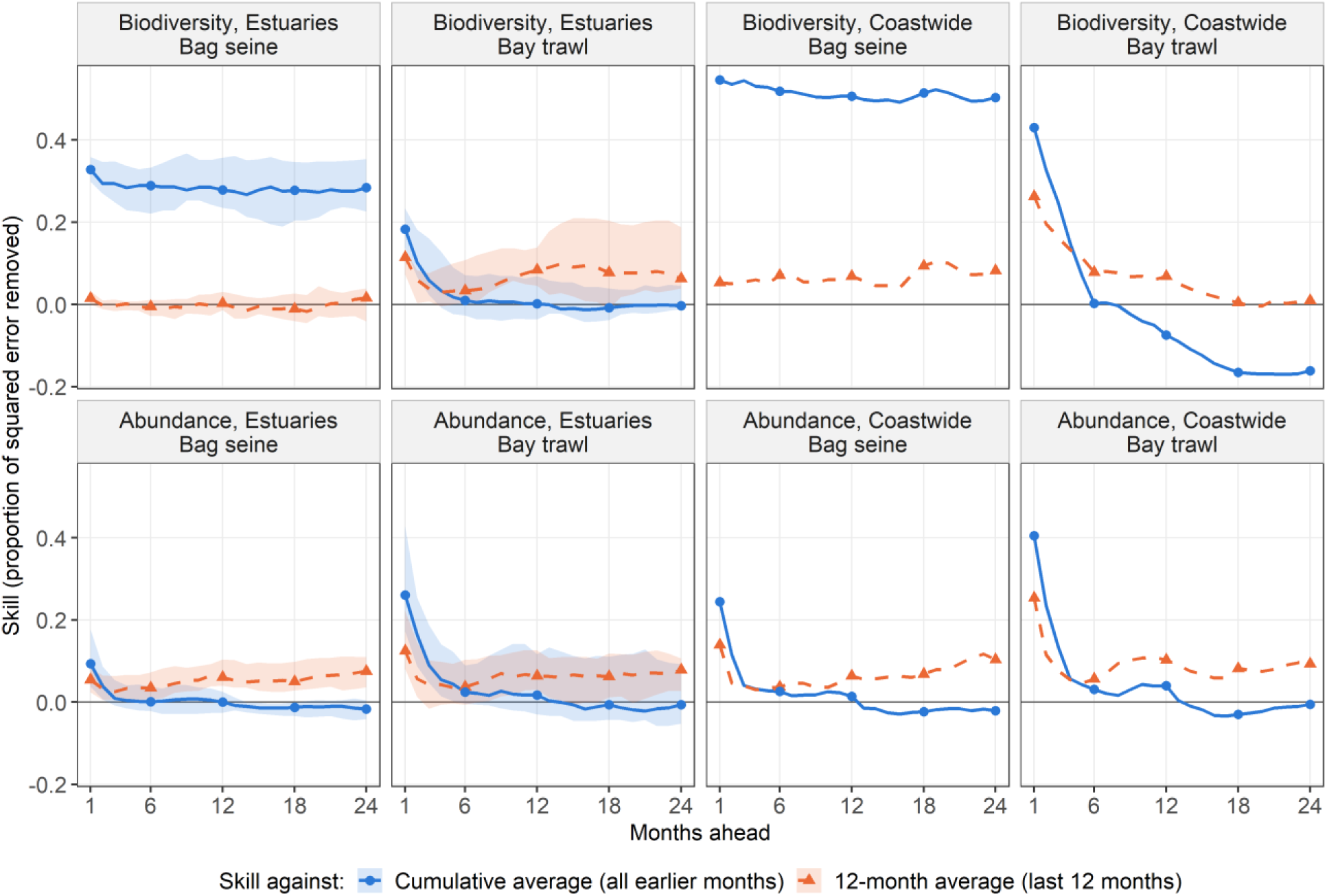
Skill of SARIMA forecasts against the cumulative average and the 12-month average (Methods: Forecast models and references; Forecast accuracy and skill), by months ahead. Solid lines and circles: skill against the cumulative average; dashed lines and triangles: skill against the 12-month average. Median over the series of each group (numbers as in Fig. 3); estuaries also with the band of the middle 50%. Where skill against the cumulative average is above skill against the 12-month average, the 12-month average is the better reference and recent levels persist; where it is below, departures from the seasonal curve fade and the cumulative average is the better reference

Comparing the two references showed how long recent levels persisted. One month ahead, the 12-month average was a better reference than the cumulative average in every group (median skill of the 12-month average against the cumulative average 0.04 to 0.52). Therefore, the forecasts had lower skill against it than against the cumulative average (median 0.01 to 0.26; Table S6). From 12 to 24 months ahead, it remained better only for bag seine biodiversity (0.28 to 0.29 in the estuaries and 0.43 to 0.47 coastwide). In the other groups, its mean squared error was 5% to 17% larger than that of the cumulative average, as the departures from the seasonal curve faded. In the estuaries, bag seine biodiversity forecasts had near-zero skill against the 12-month average (median −0.02 to 0.02 at 1 to 24 months). This suggests that their skill against the cumulative average came from knowing the recent level.

In the estuaries, the skill horizon against the cumulative average was longer for biodiversity than for abundance (Fig. S5). For biodiversity, it had a median of 3 months (n = 42 series), and 11 series stayed better than the cumulative average for all 24 months. For abundance, it had a median of 1 month (n = 112), and 85 series had a horizon of 2 months or less, including 30 whose forecasts were never better than the cumulative average (a horizon of 0).

The accuracy horizon depended strongly on the tolerance (Fig. S2). For estuary biodiversity, 7 of the 42 series (17%) were within the 30% tolerance at least one month ahead, and for at most 6 months; with a 40% tolerance, 81% were. All six coastwide biodiversity series were within the 30% tolerance for 4 to 20 months. For estuary abundance, the percentage of series within the tolerance at least one month ahead rose gradually, from 12.5% at 0.5 log units to 90% at 0.9 log units.

The 80% prediction intervals of the SARIMA forecasts contained close to the nominal 80% of the observed values for abundance and for estuary bay trawl biodiversity (medians of 78% to 84% at 1 to 24 months ahead; Fig. S6). They were too narrow for bag seine biodiversity in the estuaries (70% to 75%) and for coastwide bay trawl biodiversity at long lead times (from 83% one month ahead to 70% to 72% at 16 to 24 months).

### Estuaries and the coast

Coastwide forecasts were more accurate than estuary forecasts at every lead time, and their skill was higher one month ahead in all four groups (Fig. 5; Table S6). For example, the median skill against the cumulative average one month ahead was 0.55 coastwide against 0.33 in the estuaries for bag seine biodiversity, and 0.41 against 0.26 for bay trawl abundance. However, this advantage in skill lasted only a few months. By 12 months ahead, the skill at both scales was close to zero in three of the four groups. Therefore, the coastwide series had smaller errors, but they did not extend how far ahead the forecasts were better than the cumulative average.

**Fig. 5.**
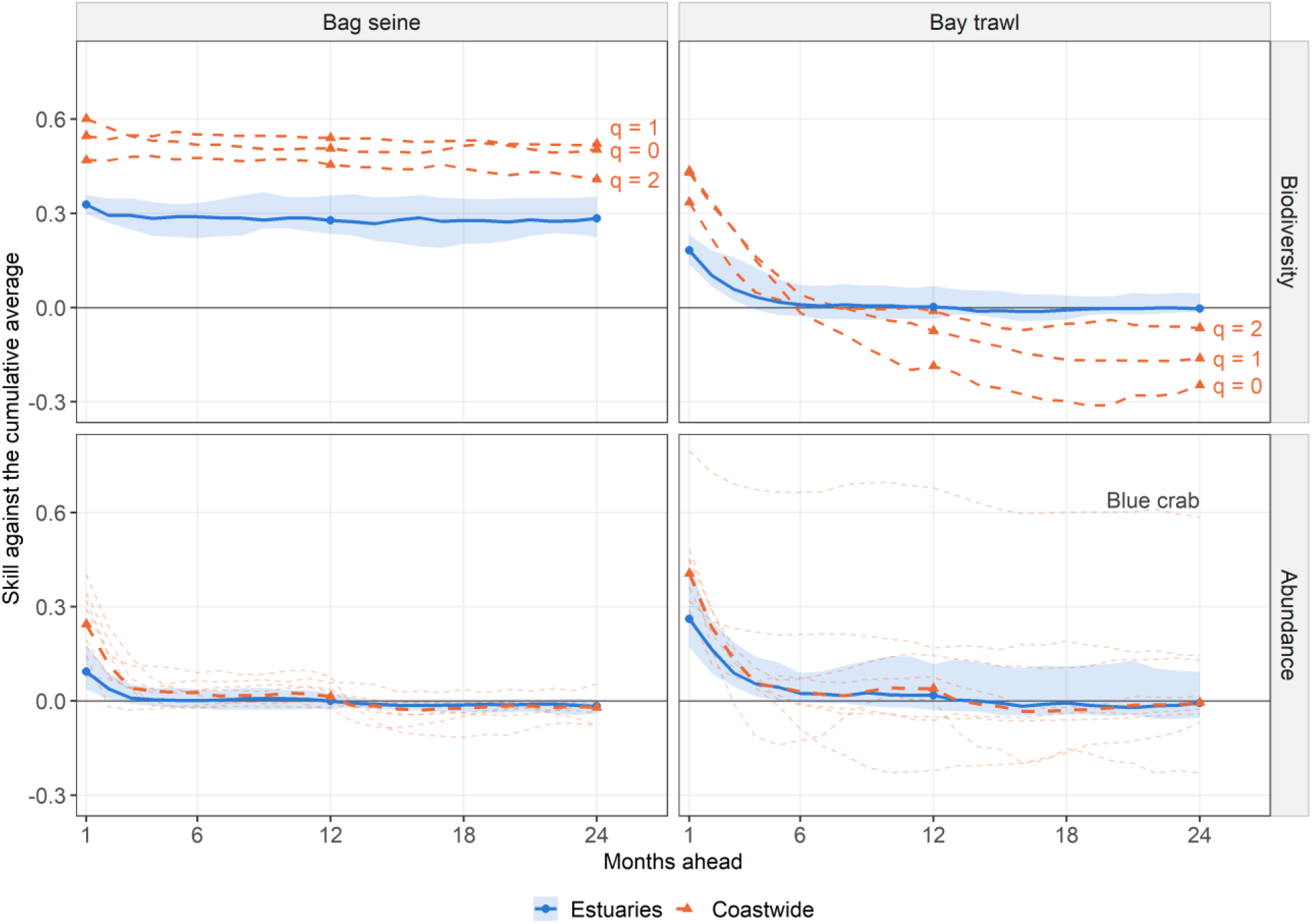
Single estuaries against the coast as a whole. Skill of SARIMA forecasts against the cumulative average, by months ahead. Estuaries (solid lines and circles): median over the estuary series (numbers as in Fig. 3) and band of the middle 50%. Coastwide (dashed lines and triangles): each biodiversity order (labeled); for abundance, the median over the eight taxa (thick line) and each taxon (thin lines)

There were two exceptions. Forecasts of coastwide bag seine biodiversity kept higher skill than those in the estuaries at every lead time (median skill 0.51 and 0.50 at 12 and 24 months ahead, against 0.28 in the estuaries; n = 3 and 21 series). Most of this skill came from 2020 to 2024 (median skill 0.93, 0.78, and 0.64 at 1, 12, and 24 months ahead, against 0.02 to 0.08 before 2020). Forecasts of coastwide bay trawl blue crab kept their skill for 24 months (0.68 at 12 months and 0.58 at 24 months), and this series declined steadily over the record (Mann-Kendall test on the yearly means, tau = −0.67, n = 43 years, p < 0.001). In contrast, coastwide bay trawl biodiversity forecasts became worse than the cumulative average from approximately 8 months ahead (median skill −0.07 at 12 months and −0.16 at 24 months; n = 3), most for q = 0 (−0.25 at 24 months). At 24 months, this loss occurred from 1992 to 2007 and from 2020 to 2024 (median skill −0.28 in both periods), but not from 2008 to 2019 (0.08). The yearly level of this series rose and fell back within a few years, most clearly from 1994 to 1997 (Fig. S3).

In a few months, many estuaries had large forecast misses together (Fig. 6). In an average month, 5% of the estuary series had a large miss. In July 2020, 21 of the 42 estuary biodiversity series (50%) had one, all higher than forecast (17 of them bag seine series), and in August 2020, 16 (38%) did. For abundance, 25 of the 112 estuary series (22%) had a large miss in July 2015, 23 of them higher than forecast. Large abundance misses were less frequent after mid-2020 (2% of the estuary series in an average month, against 6% before).

**Fig. 6.**
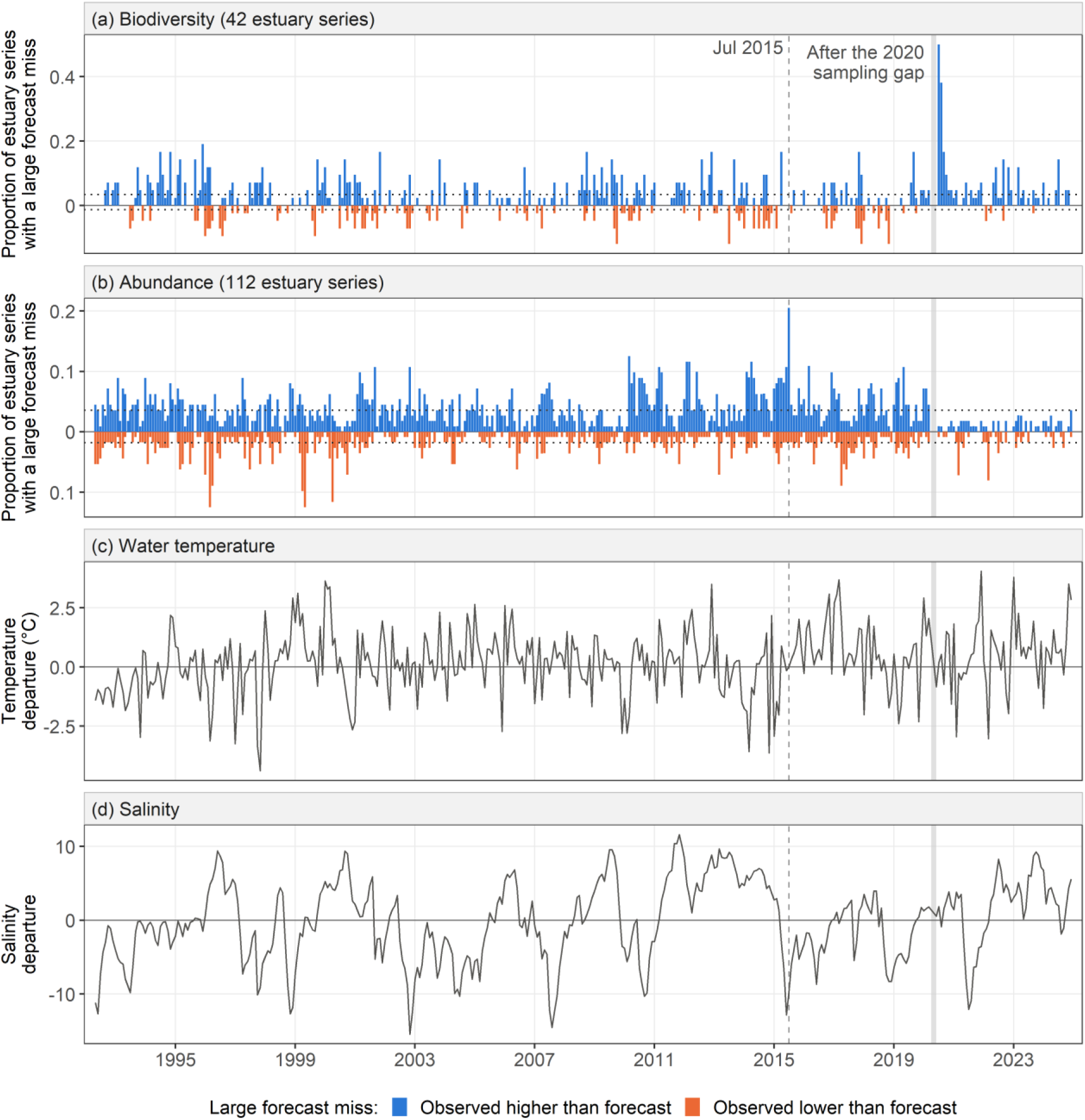
Months in which many estuaries missed together, with water temperature and salinity. (a, b) Proportion of estuary series with a large forecast miss (Methods: Forecast accuracy and skill) in each month, pooled over both sampling methods (42 biodiversity and 112 abundance series), drawn above the line when the observed value was higher than forecast and below when it was lower, both as positive proportions. Dotted lines: the average proportion over all months, in each direction. (c) Water temperature and (d) salinity: departure from the usual value for the calendar month, averaged over the estuaries and both sampling methods. Gray band: April and May 2020, not sampled; no one-month forecasts exist for June 2020, so the rise in bag seine biodiversity between March and June 2020 first appears in July

### Biodiversity, abundance, and sampling methods

Biodiversity forecasts were more accurate than abundance forecasts, but their skill was not higher. One month ahead in the estuaries, the median RMSE of biodiversity, 32% to 33% of the series mean, was approximately 0.28 log units, against 0.65 to 0.66 log units for abundance (Table S6). Before 2020, the median skill against the cumulative average one month ahead in the estuaries was 0.03 to 0.05 for bag seine biodiversity and 0.15 to 0.16 for bay trawl biodiversity (Fig. S7), against 0.05 to 0.14 and 0.22 to 0.27 for bag seine and bay trawl abundance (ranges over the periods 1992 to 2007 and 2008 to 2019).

Bag seine biodiversity increased in 2020 and remained high through 2024 (Fig. S7). In the estuaries, the mean departure from 2020 to 2024 was 1.77 standard deviations higher than from 2010 to 2019 (median across series; IQR 1.66 to 1.86), and it was higher in all 21 series. In the same period, bag seine hauls in the seven estuaries caught a median of 35% as many individuals as from 2010 to 2019 (range 21% to 46%), with approximately the same number of species per month (median ratio 1.01). The median skill of the forecasts against the cumulative average was 0.02 to 0.05 from 1992 to 2007 and from 2008 to 2019, but 0.80 one month ahead and 0.61 at 12 months from 2020 to 2024. Against the 12-month average, it was −0.04 and −0.10 from 2020 to 2024. One month ahead, forecast error changed little after the increase (median RMSE 36% of the series mean, against 32% to 33% before 2020), but the error of the cumulative average more than doubled (79%, against 32% to 34%), because it still averaged in the lower values before 2020. The forecasts followed a level that had already shifted; they did not anticipate the shift. The one-month skill of bay trawl biodiversity also increased over the three periods (0.15, 0.16, and 0.34), and in each period it declined to 0.05 or less by 6 months ahead.

Accuracy decreased slightly from q = 0 to q = 2, whereas skill differed little among the three Hill orders. In the estuaries one month ahead, the median RMSE was 28%, 31%, and 35% of the series mean for q = 0, 1, and 2, and the median skill against the cumulative average was 0.28, 0.30, and 0.25, respectively (n = 14 series each).

Among the eight taxa, blue crab forecasts had the highest skill (median skill against the cumulative average 0.34 one month ahead and 0.15 at 12 months; n = 14 series per taxon), and menhaden forecasts the lowest (0.05 one month ahead). Menhaden, white shrimp, and Atlantic croaker accounted for 105 of the 200 largest abundance misses (46, 30, and 29), measured in standard deviations of each series. One month ahead, the skill of bay trawl forecasts was higher than that of bag seine forecasts for the same taxon and estuary in 49 of 56 comparisons. By 12 months ahead, few series had skill clearly above zero, more of them bay trawl than bag seine series and nearly half of them blue crab (Fig. S8).

### Models, environmental covariates, and environmental trends

SARIMA, the state-space model, ARIMAX, and the state-space model with covariates gave nearly the same accuracy and skill for a given series (Table 1; Tables S7 and S8, Online Resource 2). One month ahead, the difference in skill between the best and the worst of these four models for the same series had a median of 0.02 for biodiversity and 0.05 for abundance (n = 48 and 128 series). Adding water temperature and salinity changed the skill little at 1 to 12 months ahead (median skill against the same model without covariates −0.01 to 0.01; maximum 0.10). The trend versions gave no gain: the state-space model with a trend was worse than the one without it, with the whole 95% uncertainty range of its skill below zero, in 40% to 52% of the biodiversity series and 65% to 93% of the abundance series at 6 to 24 months ahead, and the automatic SARIMA chose a trend term in only 6% (biodiversity) and 2% (abundance) of its fits. Last month’s value, used as a forecast, was less accurate than every model (Table 1). Failed fits removed at most 1.0% of the evaluated forecasts of any model and group.

Although the covariates added little to the forecasts, the environment changed over the record (Mann-Kendall test and Sen slope on the yearly means, n = 43 years; Table S9, Online Resource 2). Water temperature increased in 12 of the 14 estuary series (Kendall’s tau 0.30 to 0.58, adjusted p ≤ 0.007), by 0.26 to 0.64 °C per decade, and coastwide by 0.30 to 0.33 °C per decade (p < 0.001). The two exceptions were bag seine samples from Galveston Bay and Aransas Bay (adjusted p = 0.21 and 0.17). Salinity had positive Sen slopes in 11 of the 14 estuary series, but no trend was significant after adjustment for multiple tests (adjusted p ≥ 0.17), and the coastwide trends were not significant (p = 0.32 and 0.58).

### Management situations

One month ahead, most estuary series were in the forecast-with-buffer situation (29 of the 42 biodiversity series and 70 of the 112 abundance series; Fig. 7; Table S6), and five of the six coastwide biodiversity series were forecast-led. By 12 months ahead, 27 of the 42 estuary biodiversity series and 93 of the 112 estuary abundance series were in the precaution situation. The series formed a continuous cloud rather than separate groups (Fig. 7), and the mean-led situation held at most 4 of the 176 series at 1, 12, or 24 months.

**Fig. 7.**
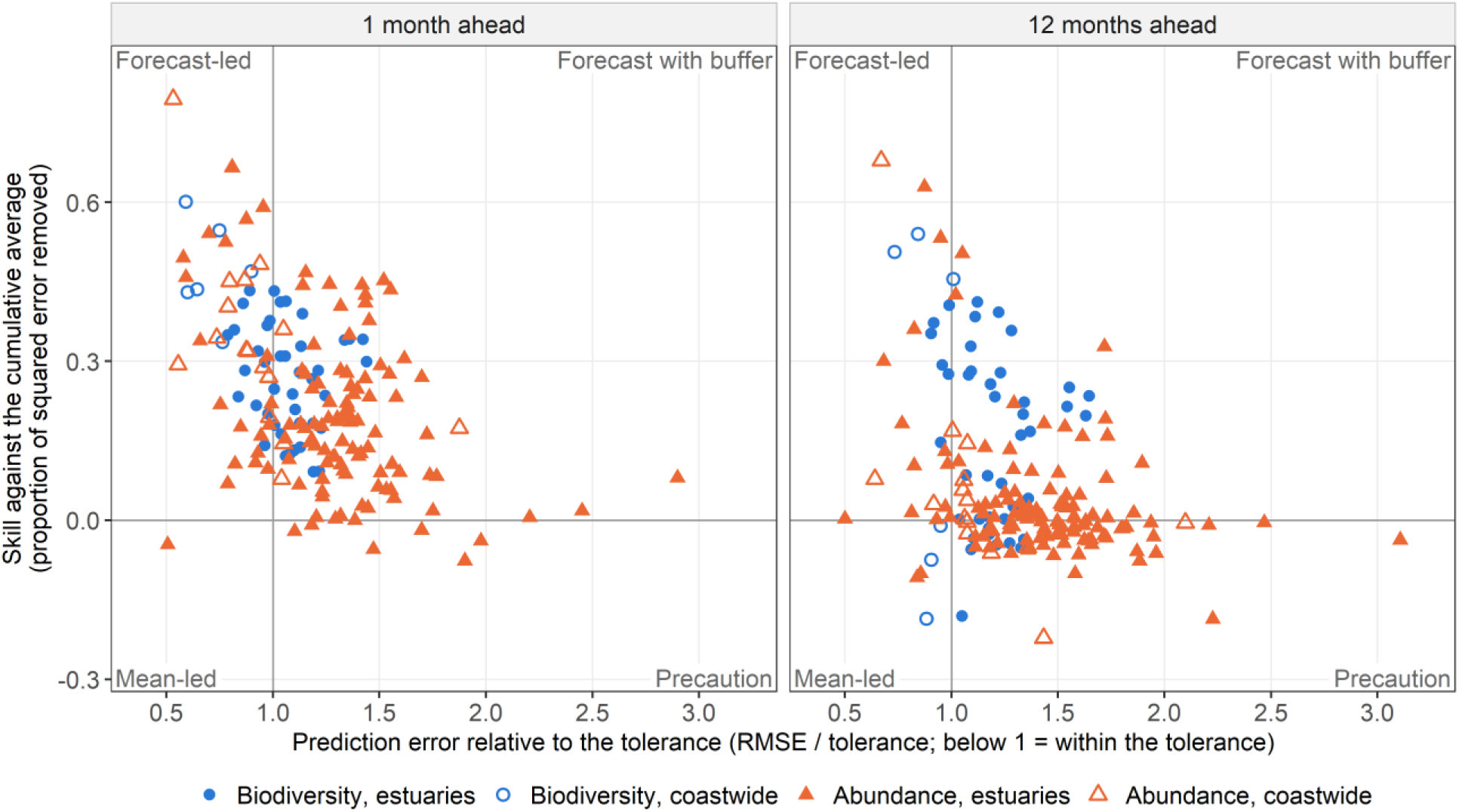
Accuracy against skill of SARIMA forecasts for every series (48 biodiversity and 128 abundance series), 1 and 12 months ahead. Accuracy: RMSE divided by the tolerance (below 1: within the tolerance). Skill: against the cumulative average (above 0: better than the cumulative average). The lines and corner labels mark the four management situations (Introduction; placement of the series: Methods: Forecast accuracy and skill): forecast-led, forecast with buffer, mean-led, and precaution. Biodiversity: blue circles; abundance: orange triangles; estuaries: filled; coastwide: open. Points are drawn at their central values; the counts in Table S6 place a series on the favorable side of a line only when its whole 95% uncertainty range clears the line

## Discussion

### Predictability beyond the seasonal cycle

The seasonal curve and the SARIMA forecasts together predicted a large part of the month-to-month variation in the biodiversity and abundance of estuarine fishes and invertebrates on the Texas coast: 27% to 50% in the estuaries and 51% to 70% coastwide (Fig. 2). In bag seine series, the largest part of this came from the seasonal cycle, which can be forecast at any lead time. In the median estuary series, approximately two-thirds of the monthly forecasts were within the tolerance one month ahead, and more than half were still within it 24 months ahead (Table S6). Beyond the seasonal cycle, the recent level added skill for one or two months, and five of the six coastwide biodiversity series were forecast-led one month ahead. Because SARIMA and the state-space model did as well without water temperature, salinity, or trend terms as with them (Table 1), such forecasts can be made from the biological data that monitoring programs already collect.

However, beyond the seasonal cycle the forecasts improved on the cumulative average mainly one or two months ahead. This was not a weakness of one model because the four models gave nearly the same accuracy and skill for a given series, and adding a trend or lagged water temperature and salinity did not help. The variation that the SARIMA forecasts did not predict (Fig. 2) is unpredictable only relative to these models and data. It includes sampling error, and other models or drivers may predict some of it. Weighted permutation entropy, a model-free measure of order in a series, was close to that of shuffled copies (group medians 0.95 to 0.985 against approximately 0.99; Fig. S4). This suggests that intrinsic predictability, how predictable a series could be at best, is also low, although observation error also raises permutation entropy (Pennekamp et al. 2019). Permutation entropy does not separate random variation from nonlinear dynamics (Şen et al. 2024), so this does not necessarily show that the dynamics are random.

### Lead time and recent levels

Forecast accuracy changed little with lead time (Fig. 3; Table S6). This differs from the common expectation that forecasts become less accurate as lead time increases (Petchey et al. 2015). Much of the accuracy came from the seasonal cycle, which does not depend on lead time, whereas the short-lived carry-over added accuracy only at short lead times. Therefore, for these series the seasonal expectation stays informative far ahead, and the question is how much the forecasts add to it.

Skill depends on both the forecast and the reference: it increases when the forecasts become more accurate and decreases when the reference becomes more accurate. High skill can therefore come from a poor reference as well as from good forecasts, as the increase in bag seine biodiversity after 2020 shows. From 2020 on, bag seine hauls caught fewer individuals with approximately the same number of species, so biodiversity standardized by sample coverage increased. The cause is not known; the sampling protocol has not changed. From 2020 to 2024, the forecasts had high skill against the cumulative average using all earlier months (0.80 one month ahead), mainly because this reference stayed close to the lower values before 2020 and missed by approximately the size of the increase every month. Against the 12-month average, which followed the new level, the forecasts had no skill (−0.04; Fig. S7). Therefore, skill should be measured against more than one reference, including one that follows recent levels (Wesselkamp et al. 2025), and checked for concentration in one period.

The data carried slow change: all 48 biodiversity series had more slow change than their lag-1 correlation alone would produce (Fig. S3). Forecasts that follow the recent level gain from a lasting shift, such as bag seine biodiversity after 2020. However, a short rise and return penalizes them. For example, coastwide bay trawl biodiversity was well above its usual level in 1994 and 1995 and below it by 1997. Which of the two is under way is often not clear when it happens. Such shifts probably also made the prediction intervals too narrow for bag seine biodiversity (Fig. S6).

### Spatial scale

Pooling the seven estuaries made the forecasts more accurate at every lead time and increased their skill one month ahead (Fig. 5; Table S6). Pooling approximately 125 hauls/tows of each sampling method each month probably averages out sampling error and variation specific to each estuary, as expected from spatial averaging of partly synchronous populations (Moran 1953; Liebhold et al. 2004; Wang and Loreau 2014). The gain in skill faded by 12 months in three of the four groups. Large misses were mostly local (Fig. 6), and pooling removes such variation but not variation shared by many estuaries, such as the misses in July 2015, which probably limited the coastwide gain to short lead times. The two coastwide series that kept their skill for 24 months, bag seine biodiversity and bay trawl blue crab, had a lasting shift and a steady decline, which the cumulative average did not follow.

### Biodiversity, abundance, and sampling methods

Biodiversity forecasts were more accurate than abundance forecasts. This is probably because combining many species into one measure reduces variability. However, before 2020 the skill of the two was similar, so combining species did not make the departures from the seasonal cycle more predictable. The three Hill orders gave similar skill. Species that are rare only in some seasons (Hewitt et al. 2016) probably change biodiversity in a similar way every year, and this part is captured by the seasonal curve.

One month ahead, bay trawl forecasts had higher skill than bag seine forecasts for the same taxon and estuary (Fig. S8), and bay trawl series carried more short-lived carry-over (16% to 32% of the variation, against 3% to 9%; Fig. 2; Table S6). It is plausible that this reflects what each method samples. Bay trawls sample deeper open-bay water, where conditions are steadier and animals may remain longer than along the shoreline, whereas bag seines sample shallow shoreline habitats, where conditions vary more under the influence of the adjacent land and many animals stay only briefly. This is consistent with the larger seasonal part of the bag seine series (25% against 8% of the variation for biodiversity and 44% against 26% for abundance in the estuaries; Fig. 2; Table S6). Movement and habitat use were not measured, so this remains a hypothesis to test.

Menhaden, white shrimp, and Atlantic croaker accounted for the most of the largest abundance misses: large, sudden changes in their catches were not anticipated even one month ahead. For conservation, taxa whose abundance changes in such pulses are the ones for which local monitoring gives the least warning, particularly when recruitment depends on conditions outside the area being monitored.

### Environmental drivers and climate change

The environment probably affected the series, as the misses shared by many estuaries in July 2015 suggest. They followed the floods of May 2015, when salinity averaged over the estuaries and both sampling methods was approximately 13 below its usual value for the month in June and approximately 10 below in July (Fig. 6). However, other large floods and the record drought of 2011 (Online Resource 1) were not followed by unusually many large misses. In addition, water temperature and salinity known at the forecast date added almost nothing at 1 to 12 months ahead. This is probably because values known at the forecast date cannot anticipate such events. Climate indices such as the El Niño-Southern Oscillation act on these estuaries largely through rainfall, freshwater inflow, temperature, and salinity. However, their signal reaches Texas estuarine salinity approximately 4 to 6 months later and in nearly all estuaries at once (Tolan 2007), so they may add lead time for misses shared by many estuaries, which values known at the forecast date did not anticipate. This is a direction for further work.

The monitoring data also show change consistent with a changing climate. Water temperature increased in 12 of the 14 estuary series and by approximately 0.3 °C per decade coastwide, and the slow change found in all biodiversity series is consistent with the long-term changes in biodiversity and species occupancy reported for this monitoring program (Fujiwara et al. 2019). Forecasts that follow the recent level adjust to such change once it appears in the data. However, three results suggest caution: the forecasts followed shifts only after they happened, the largest abundance misses shared by many estuaries followed an extreme flood, and the prediction intervals were too narrow where the level shifted. Therefore, forecasts used while conditions change need wider buffers, models refitted as new observations arrive, and regular evaluation against recent observations to detect when their skill changes.

### Comparison with other studies and generality

Pennekamp et al. (2019) found that a simple property of a time series, its permutation entropy, indicated how accurately it could be forecast. Similarly, a simple property, the lag-1 correlation, indicated where the forecasts here had skill one month ahead (rank correlation 0.94 among estuary abundance series). Beyond the seasonal cycle, the skill one month ahead came almost entirely from short-lived carry-over, the resemblance of next month to this month. The series showed little month-to-month order beyond that. Wesselkamp et al. (2025) showed that the forecast horizon depends on the reference forecast. Consistent with this, measuring skill against two references showed how much of the skill came from knowing the recent level. Studies of biodiversity change show that observed trends vary among places and can differ, even in direction, between spatial scales (Blowes et al. 2019; Leroy et al. 2023). The present results add how far ahead monthly biodiversity can be forecast at two spatial scales.

Two of the main results probably apply beyond estuaries because they rest on averaging rather than on marine processes: pooling places makes forecasts more accurate, and combining species reduces variability. How far ahead forecasts have skill probably depends on how much slow change a system carries, as the long-lead skill of bag seine biodiversity, the group with the most slow change, suggests. Environmental variation tends to be more persistent in marine than in terrestrial environments, with coastal sites in between (Steele 1985; Vasseur and Yodzis 2004). Therefore, in most terrestrial systems the seasonal cycle and the recent level are expected to be the main sources of skill, and long-lead skill is expected to be lower than here. Long, standardized terrestrial records such as breeding bird surveys could test this.

### Limitations

The long record from seven estuaries and two sampling methods allowed predictability to be compared across scales, targets, and lead times. However, the study rests on one monitoring program, on fishes and invertebrates caught by two sampling methods, and on eight common taxa for abundance. Only the taxonomic facet of biodiversity was forecast here. Functional, phylogenetic, and beta diversity can change differently from species-based measures (Monnet et al. 2014), so their predictability may also differ. The coastwide results rest on 6 biodiversity and 16 abundance series. The models were linear, with two local covariates. Nonlinear dynamics can limit or change forecasts of marine populations (Glaser et al. 2014), and nonlinear methods have been used to characterize biodiversity indicators (Otomo et al. 2023), but they were not tested here. However, nonlinear stochastic dynamics can be approximated by linear autoregressive models (Ives et al. 2003), and the approximation is probably closer for short-range forecasts when samples are frequent relative to the dynamics, as with the monthly samples here. The unpredictable part includes both sampling error and real change in the community or in how efficiently fish were caught, and the conclusions on accuracy depend on the tolerance (Fig. S2).

Long, consistent ecological records are rare (Pennekamp et al. 2019); the 461 time series they analyzed had a median length of 50 values, against approximately 510 monthly values here. How precisely the extinction risk of a population can be predicted also depends on how far ahead the prediction looks relative to the length of its record (Ellner and Holmes 2008). Where records are short, a few years of data already give a seasonal expectation and a 12-month average, and the lag-1 correlation shows whether short-range forecasting is worth attempting.

### Implications for conservation

Ecosystem-based management of the Gulf shrimp fishery shows how the results apply. Texas closes its Gulf waters to shrimp trawling from mid-May to mid-July so that brown shrimp can grow before harvest. The reopening date is set each year from TPWD bag seine and trawl samples of brown shrimp (TPWD 2025). The seasonal cycle made up approximately 60% of the month-to-month variation of brown shrimp abundance in the estuaries and 78% coastwide, and slow change and short-lived carry-over together approximately 5% (means across 14 estuary and 2 coastwide series, from the variance partition of Fig. 2). This suggests that setting the closure from the seasonal expectation adjusted to the recent level, with a buffer, uses the part that is predictable, whereas forecasts of the year’s departure would add little. Nonlinear methods have forecast Gulf penaeid shrimp abundance well (Tsai et al. 2023), and a comparison with the seasonal expectation would show whether they add skill.

I propose the following points as principles that need to be evaluated in other systems. First, for most biodiversity series, the seasonal expectation adjusted to the recent level (the seasonal curve plus a running estimate of the recent departure from it, as in the state-space model) gives forecasts nearly as good as those of any model tested one or two months ahead, and the cumulative average serves nearly as well beyond that, except in bag seine biodiversity after its increase in 2020 (Fig. S7). For abundance, SARIMA had somewhat higher skill one month ahead (Online Resource 1). Used with a buffer, it is a simple default forecast. Second, most estuary series called for forecasts with a buffer one month ahead and for precaution by 12 months, and mean-led situations almost never occurred (Fig. 7). Third, forecasts should be evaluated against two references and by period before they are trusted. Fourth, coastwide forecasts were more accurate and had higher skill one month ahead, which suits them to short-range regional planning, whereas forecasts for single estuaries need wider buffers. Long-term monitoring programs such as this one show when forecasts can be trusted and when they cannot.

## Supporting information

Online Resource 1: Supplementary methods and results

Online Resource 2: Supplementary tables

Online Resource 3: Supplementary figures

## Acknowledgments

The author thanks the Coastal Fisheries Division of the Texas Parks and Wildlife Department for collecting and providing the long-term monitoring data that made this study possible. The work was carried out during the author’s sabbatical leave. The author also thanks the editor and three anonymous reviewers for comments that improved the manuscript.

## Statements and Declarations

## Author contributions

Masami Fujiwara conceived and designed the study, performed the analyses, and wrote the manuscript.

## Data availability

The original monitoring data are available from the Coastal Fisheries Division of the Texas Parks and Wildlife Department. The prepared data files and all R code are available on GitHub and archived on Zenodo (https://doi.org/10.5281/zenodo.24348198).

## Ethics approval

This article does not contain any studies with human participants or animals performed by the author; all analyses used existing fisheries-independent monitoring data.

## Consent to participate and consent for publication

Not applicable.

## References

Bandt C, Pompe B (2002) Permutation entropy: a natural complexity measure for time series. Phys Rev Lett 88:174102. 10.1103/PhysRevLett.88.174102

Beckage B, Gross LJ, Kauffman S (2011) The limits to prediction in ecological systems. Ecosphere 2:125. 10.1890/ES11-00211.1

Benjamini Y, Hochberg Y (1995) Controlling the false discovery rate: a practical and powerful approach to multiple testing. J R Stat Soc B 57:289–300. 10.1111/j.2517-6161.1995.tb02031.x

Blowes SA, Supp SR, Antão LH, Bates A, Bruelheide H, Chase JM, Moyes F, Magurran A, McGill B, Myers-Smith IH, Winter M, Bjorkman AD, Bowler DE, Byrnes JEK, Gonzalez A, Hines J, Isbell F, Jones HP, Navarro LM, Thompson PL, Vellend M, Waldock C, Dornelas M (2019) The geography of biodiversity change in marine and terrestrial assemblages. Science 366:339–345. 10.1126/science.aaw1620

Box GEP, Jenkins GM, Reinsel GC, Ljung GM (2015) Time series analysis: Forecasting and control, 5th edn. Wiley, Hoboken

Chao A, Gotelli NJ, Hsieh TC, Sander EL, Ma KH, Colwell RK, Ellison AM (2014) Rarefaction and extrapolation with Hill numbers: a framework for sampling and estimation in species diversity studies. Ecol Monogr 84:45–67. 10.1890/13-0133.1

Chao A, Jost L (2012) Coverage-based rarefaction and extrapolation: standardizing samples by completeness rather than size. Ecology 93:2533–2547. 10.1890/11-1952.1

Clark JS, Carpenter SR, Barber M, Collins S, Dobson A, Foley JA, Lodge DM, Pascual M, Pielke R Jr, Pizer W, Pringle C, Reid WV, Rose KA, Sala O, Schlesinger WH, Wall DH, Wear D (2001) Ecological forecasts: an emerging imperative. Science 293:657–660. 10.1126/science.293.5530.657

Cooney R (2004) The precautionary principle in biodiversity conservation and natural resource management: an issues paper for policy-makers, researchers and practitioners. IUCN Policy and Global Change Series No. 2. IUCN, Gland, Switzerland and Cambridge, UK. https://portals.iucn.org/library/node/8528. Accessed 16 November 2026

Dietze MC (2017) Prediction in ecology: a first-principles framework. Ecol Appl 27:2048–2060. 10.1002/eap.1589

Dietze MC, Fox A, Beck-Johnson LM, Betancourt JL, Hooten MB, Jarnevich CS, Keitt TH, Kenney MA, Laney CM, Larsen LG, Loescher HW, Lunch CK, Pijanowski BC, Randerson JT, Read EK, Tredennick AT, Vargas R, Weathers KC, White EP (2018) Iterative near-term ecological forecasting: Needs, opportunities, and challenges. Proc Natl Acad Sci USA 115:1424–1432. 10.1073/pnas.1710231115

Doak DF, Bigger D, Harding EK, Marvier MA, O’Malley RE, Thomson D (1998) The statistical inevitability of stability-diversity relationships in community ecology. Am Nat 151:264–276. 10.1086/286117

Durbin J, Koopman SJ (2012) Time series analysis by state space methods, 2nd edn. Oxford University Press, Oxford. 10.1093/acprof:oso/9780199641178.001.0001

Ellner SP, Holmes EE (2008) Commentary on Holmes et al. (2007): resolving the debate on when extinction risk is predictable. Ecol Lett 11:E1–E5. 10.1111/j.1461-0248.2008.01211.x

Fadlallah B, Chen B, Keil A, Príncipe J (2013) Weighted-permutation entropy: a complexity measure for time series incorporating amplitude information. Phys Rev E 87:022911. 10.1103/PhysRevE.87.022911

Fujiwara M (2007) Extinction-effective population index: incorporating life-history variations in population viability analysis. Ecology 88:2345–2353. 10.1890/06-1405.1

Fujiwara M, Martinez-Andrade F, Wells RJD, Fisher M, Pawluk M, Livernois MC (2019) Climate-related factors cause changes in the diversity of fish and invertebrates in subtropical coast of the Gulf of Mexico. Commun Biol 2:403. 10.1038/s42003-019-0650-9

Glaser SM, Fogarty MJ, Liu H, Altman I, Hsieh CH, Kaufman L, MacCall AD, Rosenberg AA, Ye H, Sugihara G (2014) Complex dynamics may limit prediction in marine fisheries. Fish Fish 15:616–633. 10.1111/faf.12037

Helske J (2017) KFAS: exponential family state space models in R. J Stat Softw 78(10):1–39. 10.18637/jss.v078.i10

Hewitt JE, Thrush SF, Ellingsen KE (2016) The role of time and species identities in spatial patterns of species richness and conservation. Conserv Biol 30:1080–1088. 10.1111/cobi.12716

Hill MO (1973) Diversity and evenness: a unifying notation and its consequences. Ecology 54:427–432. 10.2307/1934352

Hsieh TC, Ma KH, Chao A (2016) iNEXT: an R package for rarefaction and extrapolation of species diversity (Hill numbers). Methods Ecol Evol 7:1451–1456. 10.1111/2041-210X.12613

Hyndman RJ, Athanasopoulos G (2021) Forecasting: Principles and practice, 3rd edn. OTexts, Melbourne. https://otexts.com/fpp3/. Accessed 16 November 2026

Hyndman RJ, Khandakar Y (2008) Automatic time series forecasting: the forecast package for R. J Stat Softw 27(3):1–22. 10.18637/jss.v027.i03

Hyndman RJ, Koehler AB (2006) Another look at measures of forecast accuracy. Int J Forecast 22:679–688. 10.1016/j.ijforecast.2006.03.001

Ives AR, Dennis B, Cottingham KL, Carpenter SR (2003) Estimating community stability and ecological interactions from time-series data. Ecol Monogr 73:301–330. 10.1890/0012-9615(2003)073[0301:ECSAEI]2.0.CO;2

Jost L (2006) Entropy and diversity. Oikos 113:363–375. 10.1111/j.2006.0030-1299.14714.x

Kendall MG (1970) Rank correlation methods, 4th edn. Griffin, London

Künsch HR (1989) The jackknife and the bootstrap for general stationary observations. Ann Stat 17:1217–1241. 10.1214/aos/1176347265

Lande R (1993) Risks of population extinction from demographic and environmental stochasticity and random catastrophes. Am Nat 142:911–927. 10.1086/285580

Leroy F, Reif J, Storch D, Keil P (2023) How has bird biodiversity changed over time? A review across spatio-temporal scales. Basic Appl Ecol 69:26–38. 10.1016/j.baae.2023.03.004

Leslie HM, McLeod KL (2007) Confronting the challenges of implementing marine ecosystem-based management. Front Ecol Environ 5:540–548. 10.1890/060093

Levin SA (1992) The problem of pattern and scale in ecology. Ecology 73:1943–1967. 10.2307/1941447

Liebhold A, Koenig WD, Bjørnstad ON (2004) Spatial synchrony in population dynamics. Annu Rev Ecol Evol Syst 35:467–490. 10.1146/annurev.ecolsys.34.011802.132516

Mann HB (1945) Nonparametric tests against trend. Econometrica 13:245–259. 10.2307/1907187

Margules CR, Pressey RL (2000) Systematic conservation planning. Nature 405:243–253. 10.1038/35012251

Martinez-Andrade F (2018) Marine resource monitoring operations manual. Texas Parks and Wildlife Department, Coastal Fisheries Division, Austin

Monnet A-C, Jiguet F, Meynard CN, Mouillot D, Mouquet N, Thuiller W, Devictor V (2014) Asynchrony of taxonomic, functional and phylogenetic diversity in birds. Glob Ecol Biogeogr 23:780–788. 10.1111/geb.12179

Moran PAP (1953) The statistical analysis of the Canadian lynx cycle. II. Synchronization and meteorology. Aust J Zool 1:291–298. 10.1071/ZO9530291

Mouquet N, Lagadeuc Y, Devictor V, Doyen L, Duputié A, Eveillard D, Faure D, Garnier E, Gimenez O, Huneman P, Jabot F, Jarne P, Joly D, Julliard R, Kéfi S, Kergoat GJ, Lavorel S, Le Gall L, Meslin L, Morand S, Morin X, Morlon H, Pinay G, Pradel R, Schurr FM, Thuiller W, Loreau M (2015) REVIEW: Predictive ecology in a changing world. J Appl Ecol 52:1293–1310. 10.1111/1365-2664.12482

Murphy AH (1988) Skill scores based on the mean square error and their relationships to the correlation coefficient. Mon Weather Rev 116:2417–2424. 10.1175/1520-0493(1988)116<2417:SSBOTM>2.0.CO;2

Otomo Y, Masuda R, Osada Y, Kawatsu K, Kondoh M (2023) Dynamics-based characterization and classification of biodiversity indicators. Ecol Evol 13:e10271. 10.1002/ece3.10271

Payne MR, Hobday AJ, MacKenzie BR, Tommasi D, Dempsey DP, Fässler SMM, Haynie AC, Ji R, Liu G, Lynch PD, Matei D, Miesner AK, Mills KE, Strand KO, Villarino E (2017) Lessons from the first generation of marine ecological forecast products. Front Mar Sci 4:289. 10.3389/fmars.2017.00289

Pennekamp F, Iles AC, Garland J, Brennan G, Brose U, Gaedke U, Jacob U, Kratina P, Matthews B, Munch S, Novak M, Palamara GM, Rall BC, Rosenbaum B, Tabi A, Ward C, Williams R, Ye H, Petchey OL (2019) The intrinsic predictability of ecological time series and its potential to guide forecasting. Ecol Monogr 89:e01359. 10.1002/ecm.1359

Petchey OL, Pontarp M, Massie TM, Kéfi S, Ozgul A, Weilenmann M, Palamara GM, Altermatt F, Matthews B, Levine JM, Childs DZ, McGill BJ, Schaepman ME, Schmid B, Spaak P, Beckerman AP, Pennekamp F, Pearse IS (2015) The ecological forecast horizon, and examples of its uses and determinants. Ecol Lett 18:597–611. 10.1111/ele.12443

R Core Team (2026) R: a language and environment for statistical computing. R Foundation for Statistical Computing, Vienna, Austria. https://www.R-project.org/

Şen B, Che-Castaldo C, Lynch HJ, Ventura F, LaRue MA, Jenouvrier S (2024) Detecting stochasticity in population time series using a non-parametric test of intrinsic predictability. Methods Ecol Evol 15:1834–1846. 10.1111/2041-210X.14423

Sen PK (1968) Estimates of the regression coefficient based on Kendall’s tau. J Am Stat Assoc 63:1379–1389. 10.1080/01621459.1968.10480934

Shumway RH, Stoffer DS (2017) Time series analysis and its applications: with R examples, 4th edn. Springer, Cham. 10.1007/978-3-319-52452-8

Steele JH (1985) A comparison of terrestrial and marine ecological systems. Nature 313:355–358. 10.1038/313355a0

Texas Parks and Wildlife Department (TPWD) (2025) Commercial Gulf shrimp season opens July 15. News release, 8 July 2025. https://tpwd.texas.gov/newsmedia/releases/?req=20250708a. Accessed 4 November 2026

Tilman D (1999) The ecological consequences of changes in biodiversity: a search for general principles. Ecology 80:1455–1474. 10.1890/0012-9658(1999)080[1455:TECOCI]2.0.CO;2

Tolan JM (2007) El Niño-Southern Oscillation impacts translated to the watershed scale: estuarine salinity patterns along the Texas Gulf Coast, 1982 to 2004. Estuar Coast Shelf Sci 72:247–260. 10.1016/j.ecss.2006.10.018

Tsai CH, Munch SB, Masi MD, Pollack AG (2023) Predicting nonlinear dynamics of short-lived penaeid shrimp species in the Gulf of Mexico. Can J Fish Aquat Sci 80:57–68. 10.1139/cjfas-2022-0029

Vasseur DA, Yodzis P (2004) The color of environmental noise. Ecology 85:1146–1152. 10.1890/02-3122

Wang S, Loreau M (2014) Ecosystem stability in space: α, β and γ variability. Ecol Lett 17:891–901. 10.1111/ele.12292

Ward EJ, Holmes EE, Thorson JT, Collen B (2014) Complexity is costly: a meta-analysis of parametric and non-parametric methods for short-term population forecasting. Oikos 123:652–661. 10.1111/j.1600-0706.2014.00916.x

Wesselkamp M, Albrecht J, Pinnington E, Castillo WJ, Pappenberger F, Dormann CF (2025) The ecological forecast limit revisited: potential, absolute and relative system predictability. Methods Ecol Evol 16:1521–1541. 10.1111/2041-210X.70049

Williams BK (2011) Adaptive management of natural resources: framework and issues. J Environ Manage 92:1346–1353. 10.1016/j.jenvman.2010.10.041

