## Supplementary material for "Forecasting estuarine biodiversity and population abundance for conservation: spatial scale, forecast horizon, and choice of measure": Online Resource 1: Supplementary methods and results

*Biodiversity and Conservation*

Masami Fujiwara

Department of Ecology and Conservation Biology, Texas A&M University, College Station, TX 77843-2258, USA

#### Part 1. Supplementary methods

##### 1.1 Sampling

The data come from the Texas Parks and Wildlife Department (TPWD) Coastal Fisheries Marine Resource Monitoring Program (Martinez-Andrade 2018). The program uses stratified cluster sampling, in which each estuary is a stratum with a fixed number of samples per month. Each estuary is divided into grids of 1 minute of latitude by 1 minute of longitude, and grids are drawn at random and without replacement for each sampling method, estuary, and month. For bag seines, one gridlet with shoreline (of the 144 gridlets of 5 by 5 seconds in each grid) is then drawn at random, and a 15.2-m section of shoreline within it is chosen at random. For bay trawls, the tow is made at the center of the drawn grid, or at the center of its area that can be sampled. All organisms caught in a haul/tow form one sample, which is a cluster of individuals. Half of each month's samples are taken from the 1st to the 15th of the month and half from the 16th to the end. All organisms longer than 5 mm are identified to the lowest possible taxon (species where possible) and counted.

The two sampling methods cover different habitats. Bag seines are 18.3 m long and 1.8 m deep, with 19-mm stretched mesh in the wings and 13-mm mesh in the bag. They are pulled parallel to the shoreline over a distance of 15.2 m, with a 12.2-m line between the two poles to keep the width constant, so each haul samples a similar area of shallow shoreline habitat. Bay trawls are 6.1-m-wide otter trawls with 38-mm stretched mesh throughout. They are towed at 4.8 km/h (3 miles per hour) for 10 min in open-bay grids in which at least one-third of the area is 1 m deep or more at mean low tide, so each tow samples a similar volume of water. The five larger estuaries (Galveston, Matagorda, San Antonio, Aransas, and Corpus Christi bays) are divided into upper and lower zones of equal size, and each zone receives half of the monthly bay trawl tows. Upper and Lower Laguna Madre are not divided (Martinez-Andrade 2018).

The seven estuaries are TPWD major areas 2 to 8. Matagorda Bay refers to the West Matagorda Bay system, including Lavaca Bay. East Matagorda Bay and Cedar Lakes are separate TPWD areas and were not used. Sabine Lake (major area 1) was excluded from the analysis. Its sampling started in January 1986, four years after the start of the record used here, and its water is much fresher. Its median monthly salinity was 6.8 in bag seine samples and 6.6 in bay trawl samples, against coastwide medians of 25.6 and 24.7 in the seven estuaries analyzed.

The bag seine and bay trawl records used here run from January 1982 to December 2024 (516 months). No samples were taken by either method in April or May 2020 because of COVID-19 restrictions. Each estuary was sampled by bag seine in 514 months, and Galveston, San Antonio, and Aransas bays were sampled by bay trawl in 514 months. Matagorda Bay, Corpus Christi Bay, and Upper and Lower Laguna Madre were sampled by bay trawl in 510 months because their bay trawl sampling started in May 1982 (Table S2; Martinez-Andrade 2018). In total, 64,814 bag seine hauls and 65,296 bay trawl tows were analyzed (Table S1).

Sampling effort changed during the record. Bag seine effort was 10 hauls per estuary and month from 1982 to 1987. It increased to 12 during 1988, to 16 in 1990, and to 20 in January 1992, and it remained at 20 until December 2024 with few exceptions (Sect. 1.4). Bay trawl effort was 20 tows per month in each of the five larger estuaries and 10 in each Laguna Madre estuary for most of the record. It was higher before 1986 in four of the larger estuaries (approximately 25 to 37 tows per month) and in Lower Laguna Madre (15 to 19), and it was higher in all seven estuaries from 1992 to 1995 (26 tows per month in the larger estuaries and 16 in each Laguna Madre estuary). Over the whole record, the mean number of bag seine hauls was 18.0 per estuary and month (Table S1). The mean number of bay trawl tows was 20.6 to 21.4 in the five larger estuaries, 10.6 in Upper Laguna Madre, and 11.2 in Lower Laguna Madre. To account for these changes, abundance was calculated per haul/tow (Sect. 1.5), and biodiversity was standardized to a common sample coverage (Sect. 1.4).

##### 1.2 Taxa

Only fishes and invertebrates were included. Each species code in the TPWD species list was treated as one taxon, and records identified only to genus or a higher rank (for example, the genus *Palaemonetes* or the family Mugilidae) were kept as separate taxa. The only exception is the silversides: the records of inland silverside (*Menidia beryllina*), tidewater silverside (*Menidia peninsulae*), and the genus *Menidia* were combined into one taxon (*Menidia* spp.) in every sample, because most records identified only to the genus are likely to be one of these two species rather than a separate group of species.

Some invertebrates were removed because their records do not give counts of individuals that can be compared among samples and years. They fall into four groups (the number of codes in each group is given in the note to Table S3). First, taxa recorded mostly as present without a count were removed. These are colonial, gelatinous, or sessile animals, such as comb jellies (Ctenophora), some jellyfish, hydroids and hydromedusae (Hydrozoa), bryozoans (Bryozoa), sponges (Porifera), corals and sea pens (Anthozoa), and sea squirts recorded only to class (Ascidiacea). No count was given for 83% of their bag seine records and 84% of their bay trawl records, and coverage-based Hill numbers need counts of individuals. Second, colonial cnidarians that are counted as colonies rather than individuals were removed. Third, three sea squirts that were counted (*Molgula manhattensis*, *Styela plicata*, and *Ciona intestinalis*) were removed because they may not have been recorded consistently in the early years of the survey. Fourth, codes that are not in the TPWD species list and could not be identified were removed (seven codes, one to eight individuals each, recorded from 2021 to 2024). A removed taxon was removed from every sample, including the samples in which it was counted.

Other invertebrates of similar kinds were kept because they are counted as individuals. These include the jellyfish that are counted and sessile animals that are not colonial, such as oysters, mussels, and sea anemones. The removed taxa made up 10.0% of the bay trawl catch by number, mostly sea squirts (9.3%), and 0.2% of the bag seine catch. Records with a catch of zero (24 records, all from 2020 to 2024) were treated as entry errors and removed.

##### 1.3 Water temperature and salinity

Water temperature (°C) and salinity were measured at nearly every haul and tow (values were missing for less than 0.1% of hauls/tows). They were measured at the surface (0 to 15 cm) approximately 3 m from the shore for bag seines and 0.3 m above the bottom for bay trawls (Martinez-Andrade 2018). For each sampling method, the values were averaged over all hauls/tows in each estuary and month. The coastwide value was the mean over all hauls/tows of the seven estuaries in the month, so it is weighted by sampling effort. As with coastwide biodiversity (Sect. 1.4), the coastwide value was calculated only for months in which every estuary was sampled. Therefore, the coastwide bay trawl series starts in May 1982.

In bag seine samples, the median monthly temperature was 24.2 to 26.0 °C among estuaries. Salinity was generally higher toward the south: the median monthly salinity was 17.3 in Galveston Bay, 20.2 to 20.6 in Matagorda, San Antonio, and Aransas bays, 30.4 in Corpus Christi Bay, 32.0 in Lower Laguna Madre, and 38.6 in Upper Laguna Madre, where the 90th percentile reached 49.2. Bay trawl samples, taken in deeper water, were 0.7 to 1.8 °C cooler and showed the same gradient in salinity. Coastwide, the median monthly temperature was 24.9 °C in bag seine samples (10th to 90th percentile, 16.1 to 30.9 °C) and 23.8 °C in bay trawl samples (14.3 to 29.9 °C). The median monthly salinity was 25.6 (18.0 to 31.9) and 24.7 (17.7 to 31.0), respectively.

Most of the largest departures of the coastwide monthly means from the median of the same calendar month coincided with natural events. Salinity was 10 to 16 below the usual value after floods in early 1992 (NWS 1995), November 1998 (Slade and Persky 1999), November 2002 (FEMA 2002), August 2007 (OSC 2007; Knabb 2008), and June 2015 (NWS Austin/San Antonio 2015). It was 10 to 12 above the usual value during the record drought of 2011 (Nielsen-Gammon 2012). Water temperature was approximately 7 °C below the usual value in December 1989, during a deep freeze that killed large numbers of fish in Texas bays (McEachron et al. 1994). These departures were seen in all seven estuaries.

Dissolved oxygen was also measured at nearly every haul and tow, but it was not used as a covariate. Its monthly means for each estuary stayed above the hypoxic level of 2 mg/L (minimum 2.8 mg/L), and the 10th to 90th percentile was 6.0 to 9.5 mg/L in bag seine samples and 5.7 to 9.3 mg/L in bay trawl samples. Water temperature and salinity explained 42% (bag seine) and 64% (bay trawl) of the variation in these monthly means (linear regression), so dissolved oxygen would have added little beyond them.

##### 1.4 Biodiversity estimation

For each estuary, sampling method, and month, the catches of all hauls/tows were pooled into one list of the number of individuals of each taxon. For the coast as a whole, the catches of all hauls/tows of the seven estuaries were pooled in the same way, so each estuary contributed in proportion to its sampling effort. Bag seine effort was the same in every estuary, so the estuaries contributed equally.

Hill numbers of order q = 0, 1, and 2 were estimated at a common sample coverage of 0.90 with the function estimateD() of the iNEXT package (Hsieh et al. 2016). Sample coverage is the estimated proportion of the individuals in the community that belong to taxa already found in the sample (Chao and Jost 2012). It was estimated with the correction of Chao and Jost (2012) when no taxon was caught exactly twice. When the coverage of a pooled catch was above 0.90, the Hill numbers were obtained by rarefaction, and when it was below 0.90, by extrapolation (Chao et al. 2014). The target was the smaller of 0.90 and the 10th percentile of the observed coverage (0.97 to 0.99), so it was 0.90. The observed coverage of estuary-month catches had a median of 0.996 for bag seines and 0.991 for bay trawls. Extrapolation was needed for 1 of 3,598 bag seine catches and 18 of 3,582 bay trawl catches. In two of them, the extrapolation went beyond twice the observed number of individuals, where the estimate of q = 0 may be biased (Chao et al. 2014). These were bay trawl catches in Upper Laguna Madre in February 1989 (32 individuals) and Lower Laguna Madre in January 2001 (53 individuals). Their estimates were used in the analysis like all other values. Every coastwide catch had a coverage of at least 0.989, so no coastwide value needed extrapolation. The smallest pooled catch was 21 individuals, and no estuary or month had a pooled catch of zero.

Standardizing to a common coverage, rather than to a common number of individuals or samples, compares months at the same completeness of sampling (Chao and Jost 2012). Therefore, a month with a larger catch does not receive a higher estimate of biodiversity only because more individuals were caught. This also reduces the effect of the change in bag seine effort from 10 to 20 hauls per month (Sect. 1.1). In addition, the forecasts were evaluated for target months from May 1992 to December 2024 (Sect. 1.8), when bag seine effort was 20 hauls per estuary and month in all but 8 of the 2,758 estuary-months sampled. Taxa that are rare or absent in some seasons affect q = 0 more than q = 1 and q = 2, which weight taxa by their abundance. When such taxa appear in the same months every year, their effect is part of the seasonal cycle and is removed with the seasonal curve (Sect. 1.7; Hewitt et al. 2016).

##### 1.5 Abundance

Abundance was analyzed for eight taxa that are among the most abundant and most frequently caught taxa in both sampling methods. For each estuary, sampling method, and month, abundance was measured as the catch per unit effort (CPUE), the mean number of individuals per haul/tow. Hauls/tows in which a taxon was not caught were included with a catch of zero, so CPUE reflects all sampling effort rather than only the hauls/tows that caught the taxon. The coastwide CPUE was the mean over all hauls/tows of the seven estuaries in the month. As with biodiversity and water temperature and salinity, it was weighted by sampling effort and calculated only for months in which every estuary was sampled (Sects. 1.3 and 1.4).

Among the eight taxa, CPUE was zero in 1.2% (blue crab) to 39.7% (menhaden) of estuary-months for bag seines and in 3.0% (spot) to 22.8% (menhaden) for bay trawls. Another 27% (bag seine) and 33% (bay trawl) of the estuary-month values of these taxa were between 0 and 1. Therefore, CPUE was analyzed as log(CPUE + 1) because log(CPUE) is undefined for a CPUE of zero.

Menhaden was analyzed at the genus level (*Brevoortia* spp.). Since 2013, TPWD has recorded part of the menhaden catch only to genus, mostly small fish in the southern estuaries, and no such records were made earlier. Therefore, the catches of Gulf menhaden (*Brevoortia patronus*), finescale menhaden (*Brevoortia gunteri*), and the genus *Brevoortia* were added together in each haul/tow. From 2013 to 2024, records identified only to genus made up 6.7% of the menhaden caught by bag seine and 0.6% of those caught by bay trawl. Finescale menhaden made up 0.1% to 0.2% of the identified menhaden by both sampling methods, before and after 2013. Records identified only to family (Clupeidae) were not included.

##### 1.6 Missing months

No samples were taken in April and May 2020 (Sect. 1.1), the only months missing inside any series. The models need complete monthly series, so the two months were filled in every series and used only for fitting the models: the filled values were used when the two months fell inside the training data of a later forecast or served as a lagged value. April and May 2020 were not used as forecast dates (Sect. 1.8), and forecasts whose target month was April or May 2020 were removed from every calculation of accuracy and skill.

Each biodiversity and abundance series was filled by interpolating its departures from the mean of each calendar month. First, the series was transformed: Hill numbers to log(Hill number) and CPUE to log(CPUE + 1). Then, for each calendar month, the mean of the transformed values was calculated from the observed months up to June 2020, the first month sampled after the gap. The departure from this monthly mean was interpolated linearly between March and June 2020, the monthly mean was added back, and the result was transformed back to the original scale. A filled CPUE below zero was set to zero. The filled Hill numbers were checked to keep the order $q = 0 \geq q = 1 \geq q = 2$. Water temperature and salinity were filled by linear interpolation between March and June 2020. No fill used data from later than June 2020.

##### 1.7 Seasonal curve

Each series had its own seasonal curve, a harmonic regression on the calendar month (Shumway and Stoffer 2017):

$$y(m) = b_0 + \sum_{k=1}^{3} \left[ s_k \, sin\left(\frac{2\pi km}{12}\right) + c_k \, cos\left(\frac{2\pi km}{12}\right) \right],$$

where $m$ is the calendar month (1 to 12), $b_0$ is the intercept, and $s_k$ and $c_k$ are the coefficients of the harmonics with periods of 12, 6, and 4 months ($k$ = 1, 2, and 3). The curve was fitted by least squares to the sampled months only, so the filled months (Sect. 1.6) did not affect it. Hill numbers were used on their original scale, and abundance as log(CPUE + 1). The seasonal swing of the Hill numbers increased less than in proportion to their level, so a log scale would have over-corrected it. The curve has no trend term, and its shape does not change over time. Therefore, differences between years and long-term trends remain in the departures from the curve, which are what the models forecast. Trends were examined with the trend versions of the models (Sect. 1.9). The departure of a month is its value minus the fitted value of the curve for that calendar month, so the departures average zero over the months used for fitting.

For the forecasts, the seasonal curve of each series was fitted again at every forecast date (Sect. 1.8), to all sampled months up to and including that month. Therefore, no forecast used a seasonal curve that had seen a later observation. The curves fitted to the whole record were used only to describe the series (Fig. S1; Table S4) and in the analysis of the characteristics of the time series data (Sect. 1.14).

The whole-record curves removed the regular seasonal cycle. After fitting, the calendar month explained 0.6% to 1.3% of the variance of the coastwide biodiversity departures, close to the 1.0% expected by chance (Table S4). For estuary biodiversity, the medians across estuaries were 0.4% to 0.9%. For abundance, the medians across estuaries were 0.2% to 2.9%, and the coastwide values were 0.2% to 5.1%. Each estuary needed its own seasonal curve because the timing and size of the seasonal cycle differed among estuaries (Table S4).

##### 1.8 Forecast design

A forecast date is the last month whose data a forecast could use. From each forecast date, forecasts were made for the target months 1 to 24 months later. The number of months between the forecast date and the target month is the lead time. Each forecast used all months from the start of the series up to and including the forecast date, so the training data grew by one month at each new forecast date (an expanding window). The models were fitted from the 13th month of each series onward, the first month for which the value 12 months earlier is available as a covariate (Sect. 1.9). Therefore, the models with and without covariates were fitted to the same months. The seasonal curve was fitted to all sampled months from the start of the series (Sect. 1.7).

The forecast dates were every month from April 1992 to November 2024, except April and May 2020 (Sect. 1.6), which gave 390 forecast dates. April 1992 is the first month at which every series had at least 120 sampled months (10 years). This is set by the series that start in May 1982 (bay trawl in four estuaries and coastwide; Sect. 1.1), and the same forecast dates were used for all 176 series. Therefore, every series was evaluated over the same target months, and every forecast used at least 10 years of data.

The models without covariates and the references forecast lead times of 1 to 24 months, and the models with covariates lead times of 1 to 12 months (Sect. 1.9). From the later forecast dates, the longer lead times fall after December 2024 and cannot be evaluated. Therefore, accuracy and skill were calculated on the 367 forecast dates from April 1992 to December 2022, the last forecast date for which all 24 lead times could be evaluated. In this way, every lead time was evaluated on the same forecast dates. The comparison among periods of the target month used all 390 forecast dates (Sect. 1.11).

##### 1.9 Forecast models and references

All models and references forecast the departures from the seasonal curve fitted at the forecast date (Sect. 1.7). A forecast value was the forecast departure plus the seasonal curve in the target month. Each model was fitted once at each forecast date, and that one fit gave the forecasts for all lead times.

The first model was a seasonal autoregressive integrated moving average model (SARIMA) selected automatically with auto.arima() of the forecast package (Hyndman and Khandakar 2008). The orders were chosen by the corrected Akaike information criterion with a stepwise search, without the approximation that auto.arima() uses by default for long series. The search could include seasonal terms, which would capture any seasonal pattern left in the departures, and a linear trend term. The second model was a state-space model in which the level of the series follows a random walk and each observation is the level plus independent noise (a local level model; Durbin and Koopman 2012). It had no seasonal part because the seasonal curve had already been removed. It was fitted by maximum likelihood with the KFAS package (Helske 2017).

The models with covariates added the monthly mean water temperature and salinity of the same estuary and sampling method (for the coastwide series, the coastwide means of the sampling method, Sect. 1.3). Only values observed at or before the forecast date were used. A forecast one month ahead used the values 1 and 12 months before the target month. A forecast 2 to 12 months ahead used the values 12 months before the target month, which are known at the forecast date. The models for one month ahead and for 2 to 12 months ahead were fitted separately, and lead times beyond 12 months were not forecast. ARIMAX was the automatic SARIMA with the covariates as regression terms (a regression with SARIMA errors). The state-space model with covariates was the state-space model with the covariates added as regression terms with fixed coefficients. For this model, each covariate was standardized with the mean and standard deviation of the training months, which changes only the scale of the coefficients.

Four trend versions tested whether a trend in the departures improved the forecasts (Sect. 2.1). The SARIMA without trend and the SARIMA with a linear trend kept the orders chosen by the automatic SARIMA at each forecast date and were refitted with the trend term removed or always included. A linear trend cannot be included in a model with two differences, so in those cases the SARIMA with a linear trend gave no forecast. The third trend version used exponential smoothing with a damped trend (Hyndman and Athanasopoulos 2021), hereafter the damped trend model. Exponential smoothing estimates the current level and slope of a series from past months, giving less weight to older months. With a damped trend, the slope of the forecast shrinks with each month ahead, so the forecast levels off instead of continuing in a straight line. This tested whether a trend helps when it is allowed to fade with lead time. The model had additive errors and was fitted with ets() of the forecast package. The fourth trend version, the state-space model with a trend, added to the state-space model a slope that also follows a random walk (a local linear trend model). Its forecast continues the current slope in a straight line at every lead time.

The three references needed no fitting. The cumulative average forecast every target month as the mean departure of all sampled months up to the forecast date. Because the seasonal curve was fitted to the same months, this mean is zero, and the forecast equals the seasonal curve. The 12-month average forecast the mean departure of the sampled months among the last 12 months. Last month's value forecast the departure of the forecast date for every lead time. The filled months were not used by the references.

The 80% and 95% prediction intervals were obtained from each fitted model on the departure scale, assuming normal errors, and the seasonal curve was added to their limits. For the SARIMA models, ARIMAX, and the damped trend model, they came from forecast() of the forecast package, and for the state-space models, from the Kalman filter in KFAS. The references had no prediction intervals.

A fit that failed was not replaced. Its forecasts were recorded as missing with the reason, and they were left out of the calculation of accuracy and skill. A SARIMA fit failed when the optimizer did not converge or when a coefficient or the error variance was not finite. A state-space fit failed when the optimizer did not converge. It also failed when all variances fell below $10^{-6}$ times the variance of the training departures, or when the standard deviation of a forecast fell below 1% of that of the training departures. These two checks catch fits that report almost no uncertainty. A state-space fit that failed was started again from up to four other sets of starting values. Failed forecasts made up less than 1% of the evaluated forecasts of every model in every group of series (Tables S7 and S8). The state-space model, the damped trend model, and the state-space model with covariates had almost none.

##### 1.10 Forecast accuracy and tolerance

The prediction error of a forecast was the forecast value minus the observed value in the target month. The forecast and the observation were both expressed as departures from the same seasonal curve, the one fitted at the forecast date (Sect. 1.7), so the error is the same on the departure scale and on the original scale. For each series, model, and lead time, the root mean squared error (RMSE) was calculated over the 367 forecast dates up to December 2022 (Sect. 1.8).

For biodiversity, the RMSE was divided by the series mean, the mean of all sampled monthly values of the series over the whole record (filled months excluded). The result is a relative error, analogous to a coefficient of variation. The series mean includes months after the forecast date, but it was used only to express errors as a percentage. It was never used in a forecast and does not change the ranking of the models or their skill. For abundance, the RMSE is in units of log(CPUE + 1) and was not divided by anything.

The second measure of accuracy was the RMSE divided by the standard deviation of the departures over the same target months (Table S5). A value below 1 means that the forecasts were more accurate than the mean of the target months, a value that is not known at the forecast date. This measure removes the differences in variability among series, but it depends on the evaluation period.

Two further summaries described single months. The size of a monthly miss was the absolute prediction error, divided by the series mean for biodiversity. The tolerance states how large a miss can be while the forecast is still accurate enough to use. It was set at 30% of the series mean for biodiversity and 0.5 log units for abundance, and it was the same for every lead time. A fixed tolerance describes the precision needed to act, which does not change with the lead time, and skill already allows for forecasts becoming harder at longer lead times. The percentage of forecasts within the tolerance was the percentage of evaluated forecasts whose monthly miss was not larger than the tolerance.

The value of the tolerance was tied to the precision of the survey itself. For every estuary and month, the hauls/tows of that month were drawn with replacement, as many as were taken, and the monthly value was calculated again from the drawn samples. For the coastwide values, the hauls/tows were drawn within each estuary and then pooled. The standard deviation over 100 resamples for biodiversity and 1,000 for abundance is the sampling uncertainty of that month's value. This was done for the evaluated target months from May 1992 to December 2024. For one estuary and month, the median sampling uncertainty was 16% (bag seine) and 13% (bay trawl) of the series mean for biodiversity and 0.26 and 0.22 log units for abundance. For the coastwide values, which pool approximately 120 to 140 hauls/tows, it was 10% (bag seine) and 7% (bay trawl) for biodiversity and 0.17 and 0.16 log units for abundance. The tolerance is therefore approximately twice the typical sampling uncertainty of one estuary and month (1.9 to 2.3 times). Tolerances from 20% to 60% for biodiversity and from 0.3 to 0.9 log units for abundance are shown in Fig. S2 (Online Resource 3).

##### 1.11 Skill, uncertainty ranges, horizons, and management situations

Skill was calculated as 1 minus the ratio of the mean squared prediction error of a forecast to that of a reference over the same target months. A skill of 0 means no better than the reference, 1 means a perfect forecast, and negative values mean worse than the reference. Skill was calculated against the cumulative average, the 12-month average, and last month's value. The trend versions and the models with covariates were also compared with a base model without the trend or the covariates: the automatic SARIMA for the two SARIMA trend versions, the damped trend model, and ARIMAX, and the state-space model for the state-space model with a trend and the state-space model with covariates. When a fit failed, its forecasts were removed from both members of each comparison, so the model and the reference were always evaluated on the same target months.

The 95% uncertainty ranges of the RMSE and of skill were obtained with a moving-block bootstrap over forecast dates (Künsch 1989). The forecast dates of a series were resampled in blocks of 12 consecutive dates, which keeps most of the dependence between neighboring dates, and the RMSE and skill were calculated again from each of 2,000 resamples. The same resamples were used for every model and lead time of a series. The 2.5th and 97.5th percentiles gave the range.

The forecast horizon was the number of consecutive lead times, starting from one month, at which the whole 95% uncertainty range met a criterion. For the accuracy horizon, the range of the RMSE had to be within the tolerance. For the skill horizon, the range of skill had to be above zero, against the cumulative average and, separately, against the 12-month average. Counting stopped at the first lead time that failed, so the horizon was 0 when the forecasts failed one month ahead. In some series, skill was lost and then returned at a longer lead time. These later lead times were not added to the horizon, but the lead time at which skill returned was recorded (Fig. S5). The horizons were also found from the values of the RMSE and skill alone, without their ranges, as a check.

Each series was placed in one of the four management situations at 1, 12, and 24 months ahead. A series was within the tolerance when the whole 95% range of its RMSE was within the tolerance, and better than the cumulative average when the whole 95% range of its skill against the cumulative average was above zero. The four situations are forecast-led (both), mean-led (within the tolerance only), forecast with buffer (better than the cumulative average only), and precaution (neither). The situations are one summary of continuous measures, and the cut-offs could be set differently for a particular decision (other tolerances in Fig. S2).

The prediction intervals were checked by their coverage, the percentage of observed values inside the 80% and 95% intervals, for each series, model, and lead time (Fig. S6). Coverage close to the nominal level means that the intervals describe the forecast uncertainty well. Coverage below it means the intervals are too narrow.

To see whether predictability changed over time, accuracy and skill were also calculated separately for three periods of the target month: 1992 to 2007, 2008 to 2019, and 2020 to 2024 (Fig. S7). The record was divided at 2008 into two halves of similar length, and the second half was divided again at 2020, when bag seine biodiversity increased, so that the effect of that increase is seen on its own. This comparison used all 390 forecast dates, because the 367 common dates end in December 2022 and would leave most of 2023 and 2024 out of the last period. Within each period, the bootstrap was applied in the same way.

##### 1.12 Variance partition

The month-to-month variation of each series was divided into four parts with the forecasts of the automatic SARIMA. Fig. 2 and Table S6 (Online Resource 2) show the means of the four parts across series, which add up to 1, with their standard deviations. All parts come from the 367 common forecast dates (Sect. 1.8).

The first part, the seasonal cycle, is the proportion of the variance of the observed values that the seasonal curve explains: 1 minus the variance of the departures divided by the variance of the observed values. It was calculated over the target months one month ahead, leaving out months whose forecast had failed. The departures were those from the seasonal curve fitted at each forecast date (Sect. 1.7). The rest, 1 minus the seasonal proportion, is the variation of the departures, and it was divided into three parts with the skill of the SARIMA against the cumulative average. Because the cumulative average forecasts the seasonal curve, its mean squared error is approximately the variance of the departures, and this skill is approximately the proportion of the departure variance that the SARIMA forecasts explained.

Slow change is the skill that remains far ahead, the mean skill from 12 to 24 months ahead (set to 0 if negative). Short-lived carry-over is the additional skill one month ahead, the skill at one month minus the slow part (set to 0 if negative). The unpredictable part is the rest, 1 minus the slow and short-lived parts, which the SARIMA forecasts missed even one month ahead. Each of these three proportions was multiplied by 1 minus the seasonal proportion, so that the four parts add up to 1.

Another model, or one with more information, might predict some of the unpredictable part, so this part is an upper limit of the variation that cannot be predicted, and the other three parts are lower limits of the variation that can. The other models gave nearly the same skill (Table 1), so the partition would change little with another model. The slow part also includes any long-term trend, which keeps the departures above or below zero for many months.

##### 1.13 Large misses shared by many estuaries

To find months in which many estuaries missed together, the one-month forecasts of the automatic SARIMA were examined (Fig. 6). Each one-month prediction error of an estuary series was compared with the RMSE of that series one month ahead over all 390 forecast dates. A miss counted as large when it was more than twice this RMSE, in either direction. If the errors were normally distributed, approximately 5% of the misses of a series would be large.

For each target month, the proportion of estuary series with a large miss was calculated separately for biodiversity (42 series) and abundance (112 series). Misses in which the observed value was higher than the forecast were counted apart from those in which it was lower. A month in which a much larger proportion than average had a large miss in the same direction points to an event that affected many estuaries at once. For biodiversity, the three Hill orders of an estuary and sampling method were counted as separate series. They often missed together, so a proportion of series is not a proportion of estuaries. The coastwide series were not included because each is a single series pooled over the estuaries.

To relate these months to the environment, the monthly departures of water temperature and salinity were plotted on the same time axis (Fig. 6). For each estuary and sampling method, the departure is the observed monthly mean minus the mean of the same calendar month over all sampled years. The departures were then averaged over the estuaries and both sampling methods. These departures use the mean of each calendar month rather than the seasonal curve of Sect. 1.7 because they serve only to show the environmental conditions at the time; Sect. 1.3 uses the median, which is less affected by the extreme months it describes.

##### 1.14 Characteristics of the time series data

The analyses in this section asked three questions about the data themselves, independent of any forecast model. First, does the variation beyond the seasonal cycle include slow change that lasts for years, which a forecast can follow? Second, how ordered is each series? A more ordered series is, in principle, easier to forecast (Pennekamp et al. 2019). Third, did water temperature and salinity change over the record, which could cause slow change in the biological series? These analyses used the departures from the seasonal curve fitted to the whole record of each series (Sect. 1.7) because they describe the record as a whole rather than forecasts made at each forecast date.

When a series changes slowly, its departures stay above or below zero for many months. Slow change was therefore measured in two ways (Fig. S3). First, the autocorrelation of the departures was calculated at lags of 1 to 36 months. Slow change keeps the autocorrelation above zero even at long lags. Second, the departures were smoothed with a centered 25-month running mean, which replaces each month by the mean of the 25 months around it. Smoothing removes ups and downs shorter than approximately two years and keeps the slow change. The variance of the smoothed series divided by the variance of the departures is the proportion of slow change.

A positive lag-1 correlation (the autocorrelation at a lag of 1 month) also makes neighboring months similar, so it produces some slow change by itself. To tell the two apart, 1,000 first-order autoregressive (AR(1)) series were simulated for each series. In an AR(1) series, each month depends only on the month before, so its only source of slow change is its lag-1 correlation. The simulated series had the same length and the same lag-1 correlation as the observed series. When the proportion of slow change of the observed series was above the 95th percentile of its simulated series, the series had more slow change than its lag-1 correlation alone would produce.

If the forecasts gain mainly from the resemblance between consecutive months, their skill should increase with the lag-1 correlation. For each series, the lag-1 correlation was therefore related to the skill of the automatic SARIMA one month ahead, measured against the cumulative average. The Spearman rank correlation was calculated separately for each target class (biodiversity and abundance) and scale (estuaries and coast).

The order in each series was measured with weighted permutation entropy (Bandt and Pompe 2002; Fadlallah et al. 2013), following Pennekamp et al. (2019), where the method is described in detail. The measure classifies short runs of consecutive values by the order of their values and asks how evenly the possible orders occur. It ranges from 0, when one order always occurs, to 1, when all orders occur equally often, as in a random series. The weighted version gives less weight to runs with small fluctuations. Runs had 3, 4, or 5 values, with 4 as the default, and the values in a run were 1 month apart (default) or 12 months apart. The 12-month spacing compares the same calendar month in consecutive years, so it detects slow change.

Each entropy was compared with 1,000 shuffled copies of the series, which keep the values but remove all order, and with 1,000 simulated AR(1) series, as for slow change. A series was more ordered than either when its entropy was below the 2.5th percentile of the shuffled copies or of the simulated series, respectively. Two features of the data could distort the entropy. In abundance series, consecutive months with zero catch differ only by the seasonal curve, so the curve rather than the catch decides their order. Series in which more than 25% of the one-month runs contained two or more zero months were flagged, and their entropy was also calculated without those runs. In the 12-month version, ties between equal values were broken at random, and the spread over 20 random tie-breaks was recorded. The entropy was related to skill and to the measures of slow change with Spearman rank correlations (Fig. S4). It does not show by itself whether the dynamics are random or nonlinear (Şen et al. 2024).

Trends in water temperature and salinity were tested on the yearly means of the departures, using years with at least six sampled months. Yearly means remove most of the dependence between consecutive months, which the test assumes is absent. The test was the Mann-Kendall test with the Sen slope (Mann 1945; Kendall 1970; Sen 1968), for each estuary and sampling method and for the coastwide means. The p-values of the seven estuaries were adjusted for multiple tests (Benjamini and Hochberg 1995) within each variable and sampling method. The same test was applied to the yearly means of the departures of each biodiversity and abundance series.

##### 1.15 Software and code

All analyses were done in R 4.6.1 (R Core Team 2026). Hill numbers were estimated with iNEXT 3.0.2 (Hsieh et al. 2016). The SARIMA models, ARIMAX, and the damped trend model were fitted with forecast 9.0.2 (Hyndman and Khandakar 2008), and the state-space models with KFAS 1.6.0 (Helske 2017). The moving-block bootstrap, the permutation entropy, the Mann-Kendall test, and the Sen slope were written for this study in base R. The prepared data files and all R code are available on GitHub and archived on Zenodo (https://doi.org/10.5281/zenodo.24348198). The original monitoring data are available from the Coastal Fisheries Division of the Texas Parks and Wildlife Department.

### Part 2. Supplementary results

#### 2.1 Models and references

Tables S7 and S8 (Online Resource 2) give the accuracy and skill of every model and reference for each group of series at 1, 12, and 24 months ahead. One month ahead, SARIMA, the state-space model, and both with water temperature and salinity gave nearly the same skill (main text). For abundance, the state-space model had somewhat lower skill than SARIMA one month ahead (median difference for the same series 0.02 to 0.04 in the estuaries and 0.03 to 0.09 coastwide). Farther ahead, SARIMA and the state-space model sometimes gave different skill for the same series. Twelve months ahead in bay trawl biodiversity in the estuaries, the SARIMA forecasts were nearly as accurate as the cumulative average (median skill 0.00), and the state-space forecasts were less accurate (−0.07). The difference was largest in coastwide bay trawl abundance (0.04 and −0.21). In bag seine biodiversity in the estuaries, the order was reversed. There, the state-space model had slightly higher skill than SARIMA at 1, 12, and 24 months ahead (median difference for the same series 0.01 to 0.03).

At 1 and 12 months ahead, adding water temperature and salinity changed the skill of SARIMA and the state-space model very little (median change −0.01 to 0.01 in every group). Compared with the same model without covariates, the whole 95% uncertainty range of the skill was above zero in at most 5 series of a group. It was below zero in at most 13 (the state-space model with covariates, bay trawl abundance in the estuaries, 12 months ahead).

The SARIMA without trend gave nearly the same forecasts as the automatic SARIMA because the automatic SARIMA seldom chose a trend term (main text). SARIMA with a linear trend always included was less accurate far ahead. At 24 months, the whole 95% range of its skill against the automatic SARIMA was below zero in 13 of the 21 bay trawl biodiversity series in the estuaries. For abundance in the estuaries, it was below zero in 18 of the 56 bag seine series and 20 of the 56 bay trawl series. It was better in coastwide bag seine biodiversity (2 of the 3 series at 24 months), where the increase after 2020 acted like a trend. One month ahead, the damped trend model was less accurate than the automatic SARIMA, with the whole 95% range of its skill below zero, in 31 of the 56 bay trawl abundance series in the estuaries. The state-space model with a trend is described in the Results of the main text.

The three references (the cumulative average, the 12-month average, and last month's value) were also compared with one another. Last month's value was less accurate than every model in every group. One month ahead, it was still more accurate than the cumulative average in three of the four coastwide groups (skill 0.29 to 0.31). At 12 and 24 months, it was much less accurate than the cumulative average, except in coastwide bag seine biodiversity, where it followed the increase after 2020. One month ahead, the 12-month average was more accurate than the cumulative average in every group (median skill 0.04 to 0.52). It gained most in bag seine biodiversity (0.32 in the estuaries and 0.52 coastwide), where it followed the higher level after 2020 and the cumulative average did not (Fig. S7). At 12 and 24 months, it was less accurate than the cumulative average, except in bag seine biodiversity.

##### 2.2 Trends in water temperature and salinity

Table S9 (Online Resource 2) gives Kendall's tau, the Sen slope, and the p-values for water temperature and salinity in every estuary, sampling method, and coastwide (methods in Sect. 1.14). Water temperature had a positive Sen slope in every series (0.11 to 0.64 °C per decade), and the trend was significant in all but the bag seine series of Galveston Bay and Aransas Bay. Salinity showed no trend after adjustment for multiple tests, and its Sen slopes ranged from −0.17 to 1.49 per decade.

##### 2.3 Permutation entropy

Fig. S4 plots the weighted permutation entropy of every series against the skill of its SARIMA forecasts one month ahead, measured against the cumulative average. With runs of four values 1 month apart, 90 of the 176 series were more ordered than their shuffled copies, but only 7 were more ordered than AR(1) series with the same lag-1 correlation. The order in most series was therefore no more than their lag-1 correlation alone would produce. More abundance series than biodiversity series were more ordered than their shuffled copies (64 of 112 in the estuaries and 13 of 16 coastwide, against 11 of 42 and 2 of 6). Runs of three and five values gave the same picture: 57 and 122 series were more ordered than their shuffled copies, and 3 and 23 more ordered than the AR(1) series. With values 12 months apart, 59 series were more ordered than their shuffled copies and 64 more ordered than the AR(1) series.

One month ahead, the skill increased with the order (1 minus the entropy) for abundance (Spearman rank correlation 0.41 in the estuaries and 0.72 coastwide) but decreased with it for biodiversity in the estuaries (−0.41). The lag-1 correlation was more closely related to the skill in every group (0.94 and 0.98 for abundance and 0.90 and 0.77 for biodiversity in the estuaries and coastwide). With values 12 months apart, the order was related to the skill 12 months ahead for biodiversity in the estuaries (0.63) and weakly for abundance (0.23).

In 24 of the 112 estuary abundance series, more than 25% of the one-month runs contained two or more months with zero catch (8 menhaden, 8 white shrimp, 5 Atlantic croaker, 2 bay anchovy, and 1 spot series). Leaving out those runs changed the entropy of 23 of these series by −0.05 to 0.01 (median −0.005); in the remaining series (bay trawl menhaden in Lower Laguna Madre), too few runs were left to calculate it. Without these 24 series, the rank correlation with the skill in the estuaries was 0.54 instead of 0.41. In the 12-month version, the standard deviation of the entropy over 20 random tie-breaks was at most 0.02. As further checks, the entropy was also calculated from the departures from the twelve monthly means and from the departures divided by the spread of each calendar month. Neither changed the entropy by more than 0.05 (typically 0.004).

**2.4 Large misses after floods and drought**

Fig. 6 shows, for each month, the proportion of estuary series with a large miss (Sect. 1.13). In an average month, 5.3% of the 112 abundance series and 4.6% of the 42 biodiversity series had a large miss, and the 95th percentile across the 389 target months from May 1992 to December 2024 was 11.6% and 16.7%. Of the events in Sect. 1.3, only the floods of May 2015 were followed by an unusually high proportion: 12.5% of the abundance series in June 2015 and 22.3% in July 2015, most of them higher than forecast, whereas no biodiversity series had a large miss in either month. In the months after the other floods and storms (May to July 1992, the first evaluated target months, while salinity was still low; November and December 1998; November and December 2002; and August and September 2007), the proportions were 2.7% to 10.7% for abundance and 0% to 16.7% for biodiversity; the highest, 16.7% of the biodiversity series in November 2002, equaled the 95th percentile. During the drought of 2011 (September 2011 to January 2012), they were 1.8% to 8.0% and 2.4% to 9.5%. The deep freeze of December 1989 came before the first evaluated target month.

#### References


Bandt C, Pompe B (2002) Permutation entropy: a natural complexity measure for time series. Phys Rev Lett 88:174102. https://doi.org/10.1103/PhysRevLett.88.174102

Benjamini Y, Hochberg Y (1995) Controlling the false discovery rate: a practical and powerful approach to multiple testing. J R Stat Soc B 57:289-300. https://doi.org/10.1111/j.2517-6161.1995.tb02031.x

Chao A, Gotelli NJ, Hsieh TC, Sander EL, Ma KH, Colwell RK, Ellison AM (2014) Rarefaction and extrapolation with Hill numbers: a framework for sampling and estimation in species diversity studies. Ecol Monogr 84:45-67. https://doi.org/10.1890/13-0133.1

Chao A, Jost L (2012) Coverage-based rarefaction and extrapolation: standardizing samples by completeness rather than size. Ecology 93:2533-2547. https://doi.org/10.1890/11-1952.1

Durbin J, Koopman SJ (2012) Time series analysis by state space methods, 2nd edn. Oxford University Press, Oxford. https://doi.org/10.1093/acprof:oso/9780199641178.001.0001

Fadlallah B, Chen B, Keil A, Príncipe J (2013) Weighted-permutation entropy: a complexity measure for time series incorporating amplitude information. Phys Rev E 87:022911. https://doi.org/10.1103/PhysRevE.87.022911

FEMA (2002) Texas severe storms, tornadoes, and flooding (FEMA-1439-DR), declared 5 November 2002. Federal Emergency Management Agency, Washington, DC. https://www.fema.gov/disaster/1439. Accessed 16 November 2026

Helske J (2017) KFAS: exponential family state space models in R. J Stat Softw 78(10):1-39. https://doi.org/10.18637/jss.v078.i10

Hewitt JE, Thrush SF, Ellingsen KE (2016) The role of time and species identities in spatial patterns of species richness and conservation. Conserv Biol 30:1080-1088. https://doi.org/10.1111/cobi.12716

Hsieh TC, Ma KH, Chao A (2016) iNEXT: an R package for rarefaction and extrapolation of species diversity (Hill numbers). Methods Ecol Evol 7:1451-1456. https://doi.org/10.1111/2041-210X.12613

Hyndman RJ, Athanasopoulos G (2021) Forecasting: Principles and practice, 3rd edn. OTexts, Melbourne. https://otexts.com/fpp3/. Accessed 16 November 2026

Hyndman RJ, Khandakar Y (2008) Automatic time series forecasting: the forecast package for R. J Stat Softw 27(3):1-22. https://doi.org/10.18637/jss.v027.i03

Kendall MG (1970) Rank correlation methods, 4th edn. Griffin, London

Knabb RD (2008) Tropical cyclone report: Tropical Storm Erin (AL052007), 15-17 August 2007. National Hurricane Center, Miami. https://www.nhc.noaa.gov/data/tcr/AL052007_Erin.pdf. Accessed 16 November 2026

Künsch HR (1989) The jackknife and the bootstrap for general stationary observations. Ann Stat 17:1217-1241. https://doi.org/10.1214/aos/1176347265

Mann HB (1945) Nonparametric tests against trend. Econometrica 13:245-259. https://doi.org/10.2307/1907187

Martinez-Andrade F (2018) Marine resource monitoring operations manual. Texas Parks and Wildlife Department, Coastal Fisheries Division, Austin

McEachron LW, Matlock GC, Bryan CE, Unger P, Cody TJ, Martin JH (1994) Winter mass mortality of animals in Texas bays. Northeast Gulf Sci 13:121-138. https://doi.org/10.18785/negs.1302.06

Nielsen-Gammon JW (2012) The 2011 Texas drought. Tex Water J 3:59-95. https://doi.org/10.21423/twj.v3i1.6463

NWS (1995) Disastrous floods on the Trinity, Brazos, Colorado, and Guadalupe rivers in Texas, December 1991-January 1992. Natural disaster survey report. National Weather Service, Silver Spring, MD. https://repository.library.noaa.gov/view/noaa/50935. Accessed 16 November 2026

NWS Austin/San Antonio (2015) 2015 Memorial Day weekend flooding. National Weather Service, Austin/San Antonio Weather Forecast Office, New Braunfels. https://www.weather.gov/media/ewx/wxevents/ewx-20150524.pdf. Accessed 16 November 2026

OSC (2007) Texas climate report: August 2007. Office of the State Climatologist, Texas A&M University, College Station. https://climatexas.tamu.edu/products/texas-climate-bulletins/monthly/2007/august-2007.html. Accessed 16 November 2026

Pennekamp F, Iles AC, Garland J, Brennan G, Brose U, Gaedke U, Jacob U, Kratina P, Matthews B, Munch S, Novak M, Palamara GM, Rall BC, Rosenbaum B, Tabi A, Ward C, Williams R, Ye H, Petchey OL (2019) The intrinsic predictability of ecological time series and its potential to guide forecasting. Ecol Monogr 89:e01359. https://doi.org/10.1002/ecm.1359

R Core Team (2026) R: a language and environment for statistical computing. R Foundation for Statistical Computing, Vienna, Austria. https://www.R-project.org/

Şen B, Che-Castaldo C, Lynch HJ, Ventura F, LaRue MA, Jenouvrier S (2024) Detecting stochasticity in population time series using a non-parametric test of intrinsic predictability. Methods Ecol Evol 15:1834-1846. https://doi.org/10.1111/2041-210X.14423

Sen PK (1968) Estimates of the regression coefficient based on Kendall's tau. J Am Stat Assoc 63:1379-1389. https://doi.org/10.1080/01621459.1968.10480934

Shumway RH, Stoffer DS (2017) Time series analysis and its applications: with R examples, 4th edn. Springer, Cham. https://doi.org/10.1007/978-3-319-52452-8

Slade RM, Persky K (1999) Floods in the Guadalupe and San Antonio River basins in Texas, October 1998. US Geological Survey Fact Sheet 147-99. https://doi.org/10.3133/fs14799
