## Supplementary material for "Forecasting estuarine biodiversity and population abundance for conservation: spatial scale, forecast horizon, and choice of measure": Online Resource 2: Supplementary tables

*Biodiversity and Conservation*

Masami Fujiwara
Department of Ecology and Conservation Biology, Texas A&M University, College Station, TX 77843-2258, USA

**Table S1** Sampling design and community biodiversity of the seven estuaries analyzed, Texas Parks and Wildlife Department Coastal Fisheries Marine Resource Monitoring Program. Sampling months: calendar months with at least one haul or tow. Station visits: all hauls or tows over the record. Stations per month: mean number of hauls or tows per sampled month. Median biodiversity (interquartile range): Hill numbers of each estuary, sampling method, and month at the common sample coverage of 0.90 (orders q = 0, q = 1 and q = 2). Cohen's *d* compares each estuary's monthly values with those of the other six estuaries for the same sampling method (positive: higher biodiversity); the 95% range is from 2,000 bootstrap resamples of months treated as independent, so it understates the uncertainty and is indicative only.

| | | | Sampling design | | | Median biodiversity (IQR) | | | Cohen's *d* against the other estuaries [95% range] | | |
|---|---|---|---|---|---|---|---|---|---|---|---|
| **Estuary** | **Sampling method** | **Years** | **Sampling months** | **Station visits** | **Stations per month** | **q = 0** | **q = 1** | **q = 2** | ***d*, q = 0** | ***d*, q = 1** | ***d*, q = 2** |
| Galveston Bay | Bag seine | 1982 to 2024 | 514 | 9,265 | 18.0 | 9.85 (6.88) | 5.60 (4.19) | 3.82 (3.15) | -0.05 [-0.15, 0.07] | -0.21 [-0.31, -0.11] | -0.26 [-0.37, -0.16] |
| | Bay trawl | 1982 to 2024 | 514 | 10,942 | 21.3 | 11.70 (5.79) | 6.79 (3.66) | 4.72 (2.56) | -0.12 [-0.20, -0.03] | -0.11 [-0.20, -0.02] | -0.11 [-0.20, -0.03] |
| Matagorda Bay | Bag seine | 1982 to 2024 | 514 | 9,261 | 18.0 | 10.41 (7.06) | 5.64 (4.26) | 4.00 (3.09) | 0.00 [-0.11, 0.09] | -0.22 [-0.32, -0.13] | -0.29 [-0.38, -0.20] |
| | Bay trawl | 1982 to 2024 | 510 | 10,930 | 21.4 | 13.74 (7.00) | 7.60 (4.51) | 5.21 (3.28) | 0.43 [0.33, 0.54] | 0.31 [0.21, 0.41] | 0.23 [0.12, 0.33] |
| San Antonio Bay | Bag seine | 1982 to 2024 | 514 | 9,266 | 18.0 | 10.81 (5.28) | 6.82 (3.63) | 4.90 (2.85) | 0.03 [-0.05, 0.11] | 0.09 [0.00, 0.18] | 0.11 [0.02, 0.19] |
| | Bay trawl | 1982 to 2024 | 514 | 10,567 | 20.6 | 10.40 (4.75) | 6.88 (3.57) | 5.24 (3.03) | -0.42 [-0.51, -0.34] | -0.13 [-0.22, -0.04] | 0.02 [-0.06, 0.11] |
| Aransas Bay | Bag seine | 1982 to 2024 | 514 | 9,246 | 18.0 | 11.05 (5.95) | 6.94 (4.15) | 5.00 (3.24) | 0.12 [0.02, 0.21] | 0.20 [0.11, 0.29] | 0.22 [0.13, 0.31] |
| | Bay trawl | 1982 to 2024 | 514 | 10,930 | 21.3 | 11.39 (4.85) | 7.08 (3.29) | 5.20 (2.58) | -0.13 [-0.21, -0.05] | 0.02 [-0.06, 0.10] | 0.10 [0.02, 0.19] |
| Corpus Christi Bay | Bag seine | 1982 to 2024 | 514 | 9,246 | 18.0 | 11.31 (5.48) | 7.13 (4.02) | 5.03 (3.17) | 0.25 [0.16, 0.34] | 0.31 [0.20, 0.41] | 0.31 [0.21, 0.41] |
| | Bay trawl | 1982 to 2024 | 510 | 10,814 | 21.2 | 12.15 (6.11) | 6.36 (4.01) | 4.37 (2.96) | 0.03 [-0.07, 0.13] | -0.12 [-0.22, -0.02] | -0.17 [-0.26, -0.07] |
| Upper Laguna Madre | Bag seine | 1982 to 2024 | 514 | 9,266 | 18.0 | 9.36 (5.72) | 6.00 (4.14) | 4.50 (3.12) | -0.31 [-0.39, -0.22] | -0.16 [-0.25, -0.08] | -0.10 [-0.18, -0.01] |
| | Bay trawl | 1982 to 2024 | 510 | 5,387 | 10.6 | 11.61 (5.04) | 7.03 (3.43) | 4.96 (2.76) | -0.20 [-0.28, -0.11] | -0.09 [-0.18, 0.00] | -0.04 [-0.13, 0.05] |
| Lower Laguna Madre | Bag seine | 1982 to 2024 | 514 | 9,264 | 18.0 | 10.38 (4.89) | 6.33 (3.68) | 4.61 (2.83) | -0.05 [-0.13, 0.04] | 0.00 [-0.08, 0.08] | 0.01 [-0.07, 0.10] |
| | Bay trawl | 1982 to 2024 | 510 | 5,726 | 11.2 | 13.35 (6.62) | 7.20 (3.64) | 4.76 (2.84) | 0.41 [0.31, 0.52] | 0.12 [0.02, 0.22] | -0.03 [-0.13, 0.06] |

Sabine Lake was excluded from all analyses, at the estuary and the coastwide scale, and is not shown. No samples were taken in April and May 2020. Bay trawl sampling in Matagorda Bay, Corpus Christi Bay and the two Laguna Madre estuaries began in May 1982.

**Table S2** Months without sampling in the monthly series used by the forecasts. Biodiversity: Hill numbers q = 0, 1, 2; abundance: log(CPUE + 1) of eight taxa, CPUE being catch per unit effort; water temperature and salinity: monthly means used as covariates. Estuaries: seven estuaries; coastwide: all estuaries pooled, a month counting only when every estuary was sampled. All series end in December 2024.

| Data set | Scale | Sampling method | Number of series | First month | Number of months | Number of months not sampled (April and May 2020) |
|---|---|---|---|---|---|---|
| Biodiversity | Estuaries | Bag seine | 21 | Jan 1982 | 516 | 2 |
| | | Bay trawl | 21 | Jan 1982 (3 estuaries); May 1982 (4 estuaries) | 516 or 512 | 2 |
| | Coastwide | Bag seine | 3 | Jan 1982 | 516 | 2 |
| | | Bay trawl | 3 | May 1982 | 512 | 2 |
| Abundance | Estuaries | Bag seine | 56 | Jan 1982 | 516 | 2 |
| | | Bay trawl | 56 | Jan 1982 (3 estuaries); May 1982 (4 estuaries) | 516 or 512 | 2 |
| | Coastwide | Bag seine | 8 | Jan 1982 | 516 | 2 |
| | | Bay trawl | 8 | May 1982 | 512 | 2 |
| Water temperature and salinity | Estuaries | Bag seine | 7 | Jan 1982 | 516 | 2 |
| | | Bay trawl | 7 | Jan 1982 (3 estuaries); May 1982 (4 estuaries) | 516 or 512 | 2 |
| | Coastwide | Bag seine | 1 | Jan 1982 | 516 | 2 |
| | | Bay trawl | 1 | May 1982 | 512 | 2 |

No samples were taken in April and May 2020, the only months without sampling inside any series. For model fitting, these two months were filled: biodiversity and abundance by interpolating the departure from each calendar month's mean (log scale for biodiversity, log(CPUE + 1) for abundance; monthly means from the months up to June 2020 only), water temperature and salinity by linear interpolation. The filled months were never used as forecast targets or forecast dates, so they enter no accuracy or skill measure. Bay trawl sampling in Matagorda Bay, Corpus Christi Bay and the two Laguna Madre estuaries began in May 1982; these series, and the coastwide bay trawl series, start then and are not extended backwards. Forecast dates run from April 1992 to November 2024.

**Table S3** Composition of the sampled community. Catch records from the Texas Parks and Wildlife Department Coastal Fisheries Marine Resource Monitoring Program, 1982 to 2024, in the seven estuaries analyzed, summarized separately for the two sampling methods. Percentages are of the total catch for that sampling method. Taxa are reported as recorded by the survey, except that silverside records were combined at genus level (*Menidia*).

**(a) Taxonomic composition. Counts are taxa with non-zero catch; percentages in parentheses are percentages of the total catch.**

| **Taxonomic group** | **Bag seine: taxa (% of catch)** | **Bay trawl: taxa (% of catch)** |
|---|---|---|
| Fishes | 236 (64.9) | 266 (67.0) |
| Invertebrates | 175 (25.2) | 198 (32.0) |
| Not identified to species | 83 (10.0) | 74 (1.0) |
| All taxa recorded | 494 (100.0) | 538 (100.0) |

**(b) The 10 most abundant taxa for each sampling method, with the percentage of total catch in parentheses.**

| **Rank** | **Bag seine: taxon (% of catch)** | **Bay trawl: taxon (% of catch)** |
|---|---|---|
| 1 | Gulf menhaden, *Brevoortia patronus* (18.0) | Atlantic croaker, *Micropogonias undulatus* (15.1) |
| 2 | Brown shrimp, *Farfantepenaeus aztecus* (11.4) | Spot, *Leiostomus xanthurus* (14.8) |
| 3 | Grass shrimp (unidentified), Genus *Palaemonetes* (9.2) | Pinfish, *Lagodon rhomboides* (14.7) |
| 4 | Pinfish, *Lagodon rhomboides* (8.8) | Brown shrimp, *Farfantepenaeus aztecus* (8.6) |
| 5 | White shrimp, *Litopenaeus setiferus* (8.8) | White shrimp, *Litopenaeus setiferus* (5.8) |
| 6 | Sheepshead minnow, *Cyprinodon variegatus* (8.5) | Bay anchovy, *Anchoa mitchilli* (4.9) |
| 7 | Spot, *Leiostomus xanthurus* (6.3) | Atlantic brief squid, *Lolliguncula brevis* (3.8) |
| 8 | Silverside, *Menidia* (4.6) | Gulf menhaden, *Brevoortia patronus* (3.0) |
| 9 | Atlantic croaker, *Micropogonias undulatus* (3.4) | Blue crab, *Callinectes sapidus* (2.8) |
| 10 | Bay anchovy, *Anchoa mitchilli* (2.7) | Silver perch, *Bairdiella chrysoura* (2.7) |

**(c) The 3 most abundant taxa in each estuary, with the percentage of that estuary's catch under that sampling method in parentheses.**

| **Estuary** | **Sampling method** | **First (% of catch)** | **Second (% of catch)** | **Third (% of catch)** |
|---|---|---|---|---|
| Galveston Bay | Bag seine | Gulf menhaden (47) | White shrimp (13) | Brown shrimp (8) |
| | Bay trawl | Atlantic croaker (24) | White shrimp (19) | Spot (8) |
| Matagorda Bay | Bag seine | Gulf menhaden (28) | White shrimp (18) | Brown shrimp (10) |
| | Bay trawl | Atlantic croaker (20) | Spot (15) | Atlantic brief squid (8) |

| Estuary | Sampling method | First (% of catch) | Second (% of catch) | Third (% of catch) |
|---|---|---|---|---|
| San Antonio Bay | Bag seine | Grass shrimp (unidentified) (17) | Gulf menhaden (13) | Pinfish (13) |
| | Bay trawl | Atlantic croaker (17) | Brown shrimp (17) | Spot (12) |
| Aransas Bay | Bag seine | Grass shrimp (unidentified) (15) | Pinfish (13) | Brown shrimp (13) |
| | Bay trawl | Spot (20) | Atlantic croaker (14) | Pinfish (12) |
| Corpus Christi Bay | Bag seine | Pinfish (15) | Brown shrimp (13) | Spot (10) |
| | Bay trawl | Pinfish (31) | Spot (19) | Atlantic croaker (10) |
| Upper Laguna Madre | Bag seine | Sheepshead minnow (26) | Grass shrimp (unidentified) (14) | Brown shrimp (10) |
| | Bay trawl | Pinfish (21) | Bay anchovy (9) | Grass shrimp (unidentified) (8) |
| Lower Laguna Madre | Bag seine | Brown shrimp (19) | Sheepshead minnow (12) | Spot (11) |
| | Bay trawl | Pinfish (32) | Atlantic croaker (8) | Silver perch (7) |

Notes. "Not identified to species" holds mostly records identified only to genus or a higher rank, of fishes and invertebrates alike, so it is reported separately rather than assigned to either. Removed before analysis and not in this table: 14 (bag seine) and 16 (bay trawl) codes of colonial, gelatinous or sessile invertebrates recorded mostly as present without a count; 2 (bag seine) and 2 (bay trawl) codes of colonial cnidarians, whose colonies are not counted as individuals; 3 (bag seine) and 3 (bay trawl) codes of sea squirts, which may not have been recorded in the early years of the survey; 2 (bag seine) and 6 (bay trawl) codes of taxa not in the TPWD species list and not identified; and 8 (bag seine) and 6 (bay trawl) codes of reptiles, amphibians and birds. All panels cover the seven estuaries analyzed (Sabine Lake is not included), which the coastwide series pool.

**Table S4** Seasonal curves fitted to the whole record (harmonic regression with 12-, 6-, and 4-month waves), as shown in Fig. S1: (a) coastwide biodiversity; (b) biodiversity in the seven estuaries; (c) abundance, log(CPUE + 1), of the eight taxa, estuaries and coastwide. In (b) and in the estuary columns of (c), each cell summarizes the seven estuary values as median [minimum, maximum]. Peak months (b): the months in which the estuaries' curves peak, with the number of estuaries in parentheses. Every series has its own curve. Variance explained: proportion of the series' variance accounted for by the curve. Leftover seasonality: proportion of the variance of the departures from the curve still explained by calendar month (expected by chance approximately 1.0%). Lag-1 correlation: the correlation between the departures of consecutive months. Annual amplitude and peak to trough (a) are in effective species; relative annual amplitude (b) is the annual amplitude divided by the series mean.

**(a) Coastwide biodiversity**

| Sampling method | Hill number | Months | Annual amplitude | Peak to trough | Peak, trough | Variance explained (%) | Leftover seasonality (%) | Lag-1 correlation |
|---|---|---|---|---|---|---|---|---|
| Bag seine | q = 0 | 514 | 4.25 | 8.50 | Sep, Jan | 31.0 | 0.8 | 0.75 |
| | q = 1 | 514 | 2.09 | 4.17 | Aug, Jan | 21.2 | 1.2 | 0.67 |
| | q = 2 | 514 | 1.31 | 2.62 | Aug, Jan | 19.1 | 1.3 | 0.59 |
| Bay trawl | q = 0 | 510 | 3.55 | 7.10 | Nov, May | 28.3 | 0.8 | 0.63 |
| | q = 1 | 510 | 1.52 | 3.03 | Nov, May | 18.6 | 0.6 | 0.62 |
| | q = 2 | 510 | 0.86 | 1.72 | Nov, May | 13.3 | 0.8 | 0.56 |

**(b) Biodiversity in the seven estuaries**

| Sampling method | Hill number | Variance explained (%) | Relative annual amplitude | Peak months | Leftover, coastwide curve (%) | Leftover, own curve (%) | Lag-1 correlation |
|---|---|---|---|---|---|---|---|
| Bag seine | q = 0 | 24.4 [17.2, 36.4] | 0.26 [0.20, 0.36] | Jul (1), Aug (2), Oct (4) | 11.1 [4.7, 21.7] | 0.9 [0.3, 1.5] | 0.55 [0.43, 0.59] |
| | q = 1 | 19.2 [10.5, 30.1] | 0.23 [0.19, 0.30] | Jul (1), Aug (2), Sep (2), Oct (2) | 9.7 [6.0, 18.3] | 0.7 [0.5, 2.1] | 0.53 [0.41, 0.60] |
| | q = 2 | 14.8 [7.0, 24.7] | 0.18 [0.15, 0.28] | Jul (1), Aug (4), Sep (1), Oct (1) | 13.0 [7.6, 20.2] | 0.7 [0.4, 2.0] | 0.51 [0.39, 0.58] |
| Bay trawl | q = 0 | 15.4 [2.2, 26.1] | 0.13 [0.03, 0.25] | Mar (1), Oct (3), Nov (3) | 16.0 [9.1, 28.7] | 0.7 [0.2, 1.2] | 0.41 [0.36, 0.61] |
| | q = 1 | 10.9 [1.1, 17.8] | 0.09 [0.04, 0.21] | Mar (1), Aug (1), Sep (1), Oct (2), Nov (2) | 15.7 [4.4, 23.0] | 0.8 [0.2, 1.5] | 0.42 [0.32, 0.67] |
| | q = 2 | 8.8 [1.9, 14.1] | 0.13 [0.07, 0.19] | Mar (1), Jul (2), Aug (1), Sep (1), Oct (1), Nov (1) | 9.9 [2.6, 19.9] | 0.4 [0.2, 1.5] | 0.38 [0.28, 0.66] |

**(c) Abundance**

| Sampling method | Taxon | Variance explained, estuaries (%) | Variance explained, coastwide (%) | Leftover, estuaries (%) | Leftover, coastwide (%) | Lag-1 correlation, estuaries | Lag-1 correlation, coastwide |
|---|---|---|---|---|---|---|---|
| Bag seine | Brown shrimp | 72.7 [48.3, 80.7] | 80.9 | 2.9 [1.2, 6.8] | 2.9 | 0.25 [0.10, 0.37] | 0.39 |
| | Spot | 65.8 [57.6, 68.0] | 79.1 | 0.8 [0.4, 2.0] | 0.4 | 0.42 [0.24, 0.50] | 0.62 |
| | Pinfish | 62.9 [50.4, 75.3] | 83.1 | 1.3 [0.4, 3.0] | 0.6 | 0.38 [0.08, 0.50] | 0.55 |
| | White shrimp | 59.2 [24.9, 78.4] | 81.8 | 0.6 [0.1, 1.1] | 0.4 | 0.29 [0.16, 0.36] | 0.44 |
| | Atlantic croaker | 40.5 [18.5, 58.1] | 62.0 | 0.7 [0.3, 1.6] | 2.1 | 0.44 [0.33, 0.56] | 0.55 |
| | Menhaden | 34.6 [24.5, 48.1] | 53.2 | 0.8 [0.4, 2.2] | 0.6 | 0.19 [0.10, 0.35] | 0.38 |
| | Bay anchovy | 27.4 [4.4, 38.4] | 42.6 | 1.2 [0.3, 1.9] | 1.2 | 0.21 [0.06, 0.25] | 0.32 |
| | Blue crab | 14.8 [10.0, 24.2] | 27.1 | 0.7 [0.3, 0.8] | 0.4 | 0.30 [0.15, 0.38] | 0.49 |
| Bay trawl | Atlantic croaker | 59.6 [38.0, 75.5] | 72.0 | 0.3 [0.1, 1.6] | 0.4 | 0.55 [0.44, 0.68] | 0.72 |
| | Brown shrimp | 58.3 [15.9, 69.4] | 77.5 | 2.1 [0.2, 5.3] | 5.1 | 0.36 [0.27, 0.48] | 0.49 |
| | White shrimp | 27.2 [7.1, 52.6] | 43.1 | 0.8 [0.0, 3.3] | 1.8 | 0.44 [0.31, 0.64] | 0.56 |
| | Pinfish | 24.6 [4.8, 49.5] | 16.4 | 0.6 [0.1, 3.2] | 0.4 | 0.47 [0.25, 0.68] | 0.69 |
| | Blue crab | 21.7 [15.0, 25.7] | 28.0 | 0.2 [0.1, 0.7] | 0.2 | 0.66 [0.58, 0.74] | 0.84 |
| | Spot | 15.6 [1.8, 40.3] | 23.0 | 0.5 [0.2, 1.5] | 0.2 | 0.56 [0.51, 0.67] | 0.70 |
| | Bay anchovy | 5.0 [2.7, 11.2] | 11.5 | 0.5 [0.3, 1.1] | 0.3 | 0.45 [0.37, 0.60] | 0.54 |
| | Menhaden | 3.6 [1.1, 11.7] | 4.6 | 0.7 [0.1, 1.7] | 0.3 | 0.36 [0.11, 0.63] | 0.47 |

The forecasts do not use these curves: at each forecast date the curve was refitted on the months up to that date. From one forecast date to the next the refitted curve moved by a median of 0.01 standard deviations of the departures (at most 0.27); at the last forecast date it differed from the whole-record curve by at most 0.05. In every coastwide biodiversity series the departures were autocorrelated (Ljung-Box test at lag 12, $p < 0.001$), the month-to-month memory that the forecast models use. Leftover with the coastwide curve (b): the same measure after subtracting the coastwide curve from each estuary's series instead of the estuary's own curve. April and May 2020 (no sampling) are left out of every value. Leftover seasonality was highest for brown shrimp (bay trawl: estuaries up to 5.3%, coastwide 5.1%; bag seine: up to 6.8%, coastwide 2.9%), whose short spring peak the three waves do not follow fully.

**Table S5** Accuracy of SARIMA forecasts on a second measure, 1 and 12 months ahead: the root mean squared error (RMSE) of the out-of-sample prediction errors divided by the standard deviation of the departures from the seasonal curve over the same target months. Values below 1 mean that the forecasts were more accurate than the mean of the target months, which is not available at the forecast date. Calculated on the 367 forecast dates up to December 2022, as in the main text. Estuaries: median over the seven estuaries; coastwide: the single series. Biodiversity: Hill numbers; abundance: log(CPUE + 1), CPUE being catch per unit effort.

| Sampling method | Target | Estuaries, 1 month | Estuaries, 12 months | Coastwide, 1 month | Coastwide, 12 months |
|---|---|---|---|---|---|
| Bag seine | q = 0 | 0.91 | 0.96 | 0.73 | 0.82 |
| | q = 1 | 0.86 | 0.92 | 0.74 | 0.76 |
| | q = 2 | 0.88 | 0.92 | 0.78 | 0.80 |
| | Atlantic croaker | 0.93 | 1.00 | 0.83 | 1.01 |
| | Bay anchovy | 0.99 | 1.01 | 0.96 | 0.98 |
| | Blue crab | 0.94 | 1.00 | 0.87 | 1.00 |
| | Brown shrimp | 0.96 | 1.00 | 0.90 | 0.98 |
| | Menhaden | 1.00 | 1.01 | 0.92 | 1.02 |
| | Pinfish | 0.95 | 1.02 | 0.82 | 0.99 |
| | Spot | 0.93 | 1.01 | 0.81 | 1.05 |
| | White shrimp | 0.98 | 1.01 | 0.93 | 1.02 |
| Bay trawl | q = 0 | 0.90 | 1.00 | 0.78 | 1.11 |
| | q = 1 | 0.92 | 1.00 | 0.77 | 1.06 |
| | q = 2 | 0.93 | 1.00 | 0.83 | 1.02 |
| | Atlantic croaker | 0.86 | 0.99 | 0.75 | 1.00 |
| | Bay anchovy | 0.90 | 1.00 | 0.93 | 1.04 |
| | Blue crab | 0.84 | 1.00 | 0.71 | 0.98 |
| | Brown shrimp | 0.93 | 1.01 | 0.88 | 1.02 |
| | Menhaden | 0.95 | 1.02 | 0.87 | 1.05 |
| | Pinfish | 0.95 | 1.01 | 0.75 | 0.95 |
| | Spot | 0.85 | 1.03 | 0.72 | 1.12 |
| | White shrimp | 0.89 | 1.02 | 0.80 | 1.03 |

Menhaden is the genus *Brevoortia* (Gulf menhaden, finescale menhaden, and records identified only to the genus, combined), as in all abundance results.

**Table S6** Forecast accuracy, skill, and management situations by group (SARIMA forecasts; medians over the series of each group, with interquartile ranges in parentheses for accuracy). RMSE: root mean squared out-of-sample prediction error, as a proportion of the series mean for biodiversity and in log units (log(CPUE + 1)) for abundance, 1, 12, and 24 months ahead. Within tolerance: percentage of monthly forecasts within 30% of the series mean (biodiversity) or 0.5 log units (abundance), 1 and 24 months ahead. Skill: 1 minus the ratio of the mean squared prediction error to that of the cumulative average (all data up to the forecast date) or of the 12-month average (the average of the last 12 months); 0 means no better than that reference. Situations: numbers of series that are forecast-led (FL), mean-led (ML), forecast with buffer (FB), or precaution (P) (Fig. 7); a series counts as within the tolerance or better than the cumulative average only when the whole 95% uncertainty range clears the line. Series counts: biodiversity 21 in the estuaries (7 estuaries × q = 0, 1, 2) and 3 coastwide per sampling method; abundance 56 in the estuaries (7 estuaries × 8 taxa) and 8 coastwide. Calculated on the 367 forecast dates up to December 2022. Lower table: the division of the month-to-month variation into the seasonal cycle, slow change, short-lived carry-over, and the remainder (Fig. 2), with the sum of the first three (predictable), as means ± SD (%) across the series of each group.

| Group | Series | RMSE, 1 month | RMSE, 12 months | RMSE, 24 months | Within tolerance (%), 1 month | Within tolerance (%), 24 months | Skill vs cumulative average 1 / 12 / 24 | Skill vs 12-month average 1 / 12 | Situations at 1 month (FL / ML / FB / P) | Situations at 12 months (FL / ML / FB / P) |
|---|---|---|---|---|---|---|---|---|---|---|
| **Biodiversity (RMSE as a proportion of the series mean)** | | | | | | | | | | |
| Estuaries, bag seine | 21 | 0.33 (0.29 to 0.36) | 0.36 (0.32 to 0.40) | 0.38 (0.34 to 0.43) | 66 (62 to 73) | 62 (58 to 67) | 0.33 / 0.28 / 0.28 | 0.01 / 0.00 | 5 / 0 / 11 / 5 | 0 / 0 / 14 / 7 |
| Estuaries, bay trawl | 21 | 0.32 (0.30 to 0.34) | 0.35 (0.33 to 0.39) | 0.36 (0.33 to 0.39) | 69 (64 to 73) | 61 (57 to 63) | 0.18 / 0.00 / 0.00 | 0.11 / 0.08 | 2 / 0 / 18 / 1 | 0 / 0 / 1 / 20 |
| Coastwide, bag seine | 3 | 0.23 (0.20 to 0.25) | 0.25 (0.24 to 0.28) | 0.28 (0.26 to 0.31) | 83 (78 to 87) | 78 (73 to 81) | 0.55 / 0.51 / 0.50 | 0.05 / 0.07 | 2 / 1 / 0 / 0 | 1 / 0 / 1 / 1 |
| Coastwide, bay trawl | 3 | 0.19 (0.19 to 0.21) | 0.27 (0.27 to 0.28) | 0.29 (0.29 to 0.30) | 88 (86 to 89) | 67 (67 to 70) | 0.43 / -0.07 / -0.16 | 0.26 / 0.07 | 3 / 0 / 0 / 0 | 0 / 0 / 0 / 3 |
| **Abundance (RMSE in log units)** | | | | | | | | | | |
| Estuaries, bag seine | 56 | 0.66 (0.60 to 0.77) | 0.70 (0.64 to 0.81) | 0.72 (0.64 to 0.81) | 63 (57 to 70) | 59 (53 to 66) | 0.09 / 0.00 / -0.02 | 0.05 / 0.06 | 4 / 0 / 26 / 26 | 2 / 0 / 3 / 51 |
| Estuaries, bay trawl | 56 | 0.65 (0.52 to 0.71) | 0.73 (0.58 to 0.81) | 0.75 (0.60 to 0.83) | 62 (56 to 74) | 53 (46 to 67) | 0.26 / 0.02 / -0.01 | 0.12 / 0.06 | 8 / 2 / 44 / 2 | 3 / 2 / 9 / 42 |
| Coastwide, bag seine | 8 | 0.46 (0.39 to 0.52) | 0.54 (0.51 to 0.55) | 0.56 (0.52 to 0.58) | 77 (70 to 85) | 69 (67 to 72) | 0.24 / 0.01 / -0.02 | 0.14 / 0.06 | 4 / 0 / 3 / 1 | 0 / 1 / 0 / 7 |
| Coastwide, bay trawl | 8 | 0.45 (0.42 to 0.48) | 0.53 (0.52 to 0.61) | 0.57 (0.54 to 0.61) | 76 (73 to 79) | 65 (61 to 69) | 0.41 / 0.04 / -0.01 | 0.25 / 0.10 | 4 / 0 / 4 / 0 | 1 / 0 / 1 / 6 |

| Group | Series | Seasonal cycle (%) | Slow change (%) | Short-lived carry-over (%) | Remainder (%) | Predictable, sum (%) |
|---|---|---|---|---|---|---|
| **Biodiversity** | | | | | | |
| Estuaries, bag seine | 21 | 25 ± 9 | 21 ± 6 | 4 ± 4 | 50 ± 7 | 50 ± 7 |
| Estuaries, bay trawl | 21 | 8 ± 9 | 3 ± 5 | 16 ± 7 | 73 ± 11 | 27 ± 11 |
| Coastwide, bag seine | 3 | 35 ± 9 | 32 ± 4 | 3 ± 2 | 30 ± 8 | 70 ± 8 |
| Coastwide, bay trawl | 3 | 20 ± 10 | 0 ± 0 | 32 ± 3 | 49 ± 10 | 51 ± 10 |
| **Abundance** | | | | | | |
| Estuaries, bag seine | 56 | 44 ± 25 | 1 ± 2 | 5 ± 4 | 50 ± 23 | 50 ± 23 |
| Estuaries, bay trawl | 56 | 26 ± 22 | 6 ± 10 | 16 ± 11 | 53 ± 22 | 47 ± 22 |
| Coastwide, bag seine | 8 | 62 ± 28 | 0 ± 1 | 9 ± 9 | 30 ± 22 | 70 ± 22 |
| Coastwide, bay trawl | 8 | 36 ± 28 | 7 ± 11 | 19 ± 11 | 37 ± 22 | 63 ± 22 |

**Table S7** Results of every forecasting model and simple reference for biodiversity (Hill numbers q = 0, 1, 2), by scale and sampling method: median over the series of each group (estuaries: 21 series per sampling method, 7 estuaries × q = 0, 1, 2; coastwide: 3), calculated on the 367 forecast dates up to December 2022. All models forecast the monthly departures from the seasonal curve, which was refitted at each forecast date using only the data up to that date. RMSE: root mean squared out-of-sample prediction error divided by the series mean (a relative error), 1, 12 and 24 months ahead. Cumulative average: the seasonal curve plus the mean departure of all earlier months; 12-month average: the same with the last 12 months only. Skill: 1 minus the ratio of the mean squared prediction error to that of the cumulative average (0 = no better than the cumulative average). Base model: the model without the feature being tested (covariates or trend): SARIMA (automatic) for ARIMAX and the SARIMA and damped trend versions; the state-space model for the state-space models with covariates or a trend. Skill against the base model measures what that feature adds (0 = no gain). Better / worse: numbers of series whose bootstrap range of skill against the base model lies above / below zero, 1 and 12 months ahead. Failed forecasts: cases (series × forecast date × months ahead) lost because a model could not be fitted at a forecast date, with their percentage of the model's evaluated cases; these cases were left out of the comparison with the base model for both models. Models with covariates were run to 12 months (n/a at 24).

| Model | Role | RMSE 1 / 12 / 24 | Skill vs cumulative average 1 / 12 / 24 | Skill vs base model 1 / 12 / 24 | Better / worse than base model at 1; at 12 | Failed forecasts (% of evaluated) |
|---|---|---|---|---|---|---|
| **Estuaries, bag seine** | | | | | | |
| SARIMA (automatic) | Base model | 0.33 / 0.36 / 0.38 | 0.33 / 0.28 / 0.28 | | | 96 (0.05%) |
| State-space | Base model | 0.33 / 0.35 / 0.37 | 0.32 / 0.31 / 0.30 | | | 0 |
| ARIMAX (temperature, salinity) | Covariates | 0.33 / 0.36 / n/a | 0.33 / 0.28 / n/a | 0.00 / 0.00 / n/a | 0 / 1; 3 / 0 | 0 |
| State-space with covariates | Covariates | 0.32 / 0.35 / n/a | 0.32 / 0.31 / n/a | 0.00 / 0.01 / n/a | 0 / 0; 2 / 0 | 0 |
| SARIMA, no trend | Trend test | 0.33 / 0.36 / 0.38 | 0.33 / 0.27 / 0.26 | 0.00 / 0.00 / 0.00 | 0 / 0; 0 / 0 | 168 (0.09%) |
| SARIMA, linear trend | Trend test | 0.33 / 0.35 / 0.37 | 0.33 / 0.27 / 0.27 | 0.00 / 0.00 / 0.02 | 0 / 1; 2 / 0 | 622 (0.34%) |
| Damped trend | Trend test | 0.33 / 0.35 / 0.37 | 0.32 / 0.30 / 0.29 | 0.00 / 0.01 / 0.02 | 0 / 0; 1 / 1 | 0 |
| State-space with trend | Trend test | 0.33 / 0.35 / 0.37 | 0.32 / 0.30 / 0.30 | 0.00 / -0.01 / -0.01 | 0 / 0; 0 / 1 | 0 |
| Last month's value (naive) | Reference | 0.41 / 0.44 / 0.46 | -0.02 / -0.03 / -0.05 | | | 0 |
| 12-month average | Reference | 0.33 / 0.36 / 0.38 | 0.32 / 0.28 / 0.29 | | | 0 |
| Cumulative average | Reference | 0.40 / 0.43 / 0.46 | (reference) | | | 0 |
| **Estuaries, bay trawl** | | | | | | |
| SARIMA (automatic) | Base model | 0.32 / 0.35 / 0.36 | 0.18 / 0.00 / 0.00 | | | 120 (0.07%) |
| State-space | Base model | 0.32 / 0.38 / 0.38 | 0.16 / -0.07 / -0.06 | | | 0 |
| ARIMAX (temperature, salinity) | Covariates | 0.32 / 0.36 / n/a | 0.19 / 0.03 / n/a | 0.00 / 0.00 / n/a | 0 / 1; 3 / 0 | 0 |
| State-space with covariates | Covariates | 0.32 / 0.38 / n/a | 0.18 / -0.07 / n/a | 0.00 / -0.01 / n/a | 0 / 0; 0 / 2 | 0 |
| SARIMA, no trend | Trend test | 0.32 / 0.35 / 0.36 | 0.18 / 0.00 / 0.00 | 0.00 / 0.00 / 0.00 | 0 / 0; 2 / 0 | 240 (0.13%) |
| SARIMA, linear trend | Trend test | 0.32 / 0.37 / 0.38 | 0.17 / -0.05 / -0.05 | -0.01 / -0.04 / -0.05 | 0 / 8; 0 / 8 | 239 (0.13%) |
| Damped trend | Trend test | 0.32 / 0.38 / 0.39 | 0.15 / -0.05 / -0.05 | -0.04 / -0.07 / -0.06 | 2 / 5; 0 / 6 | 0 |
| State-space with trend | Trend test | 0.32 / 0.39 / 0.41 | 0.16 / -0.14 / -0.17 | -0.01 / -0.05 / -0.10 | 0 / 6; 0 / 16 | 0 |
| Last month's value (naive) | Reference | 0.37 / 0.45 / 0.47 | -0.10 / -0.58 / -0.58 | | | 0 |
| 12-month average | Reference | 0.33 / 0.38 / 0.39 | 0.09 / -0.12 / -0.08 | | | 0 |
| Cumulative average | Reference | 0.35 / 0.36 / 0.37 | (reference) | | | 0 |
| **Coastwide, bag seine** | | | | | | |
| SARIMA (automatic) | Base model | 0.23 / 0.25 / 0.28 | 0.55 / 0.51 / 0.50 | | | 0 |
| State-space | Base model | 0.23 / 0.26 / 0.30 | 0.53 / 0.50 / 0.47 | | | 0 |
| ARIMAX (temperature, salinity) | Covariates | 0.22 / 0.25 / n/a | 0.56 / 0.50 / n/a | 0.00 / 0.00 / n/a | 0 / 0; 0 / 0 | 7 (0.05%) |
| State-space with covariates | Covariates | 0.23 / 0.27 / n/a | 0.54 / 0.50 / n/a | 0.00 / 0.00 / n/a | 1 / 0; 0 / 0 | 0 |

| Model | Role | RMSE 1 / 12 / 24 | Skill vs cumulative average 1 / 12 / 24 | Skill vs base model 1 / 12 / 24 | Better / worse than base model at 1; at 12 | Failed forecasts (% of evaluated) |
|---|---|---|---|---|---|---|
| SARIMA, no trend | Trend test | 0.23 / 0.25 / 0.28 | 0.55 / 0.50 / 0.48 | 0.00 / 0.00 / 0.00 | 0 / 0; 0 / 0 | 0 |
| SARIMA, linear trend | Trend test | 0.23 / 0.25 / 0.28 | 0.54 / 0.52 / 0.51 | -0.01 / 0.02 / 0.03 | 0 / 0; 1 / 0 | 24 (0.09%) |
| Damped trend | Trend test | 0.23 / 0.26 / 0.30 | 0.53 / 0.50 / 0.47 | -0.04 / -0.02 / -0.01 | 0 / 0; 0 / 0 | 0 |
| State-space with trend | Trend test | 0.23 / 0.26 / 0.29 | 0.53 / 0.50 / 0.48 | 0.00 / 0.01 / 0.02 | 0 / 0; 1 / 0 | 0 |
| Last month's value (naive) | Reference | 0.28 / 0.31 / 0.36 | 0.31 / 0.29 / 0.25 | | | 0 |
| 12-month average | Reference | 0.23 / 0.27 / 0.31 | 0.52 / 0.47 / 0.44 | | | 0 |
| Cumulative average | Reference | 0.33 / 0.37 / 0.41 | (reference) | | | 0 |
| **Coastwide, bay trawl** | | | | | | |
| SARIMA (automatic) | Base model | 0.19 / 0.27 / 0.29 | 0.43 / -0.07 / -0.16 | | | 236 (0.90%) |
| State-space | Base model | 0.19 / 0.28 / 0.30 | 0.43 / -0.18 / -0.21 | | | 0 |
| ARIMAX (temperature, salinity) | Covariates | 0.19 / 0.27 / n/a | 0.43 / -0.06 / n/a | 0.00 / 0.01 / n/a | 0 / 0; 0 / 0 | 0 |
| State-space with covariates | Covariates | 0.19 / 0.29 / n/a | 0.43 / -0.19 / n/a | -0.01 / -0.01 / n/a | 0 / 0; 0 / 0 | 0 |
| SARIMA, no trend | Trend test | 0.19 / 0.27 / 0.29 | 0.43 / -0.07 / -0.16 | 0.00 / 0.00 / 0.00 | 0 / 0; 0 / 0 | 236 (0.90%) |
| SARIMA, linear trend | Trend test | 0.20 / 0.28 / 0.32 | 0.42 / -0.17 / -0.34 | -0.02 / -0.08 / -0.15 | 0 / 0; 0 / 0 | 260 (0.99%) |
| Damped trend | Trend test | 0.20 / 0.29 / 0.30 | 0.42 / -0.19 / -0.21 | -0.02 / -0.11 / -0.05 | 0 / 0; 0 / 0 | 0 |
| State-space with trend | Trend test | 0.19 / 0.30 / 0.32 | 0.43 / -0.27 / -0.41 | 0.00 / -0.06 / -0.15 | 0 / 0; 0 / 3 | 0 |
| Last month's value (naive) | Reference | 0.22 / 0.32 / 0.33 | 0.29 / -0.51 / -0.52 | | | 0 |
| 12-month average | Reference | 0.23 / 0.28 / 0.29 | 0.23 / -0.15 / -0.16 | | | 0 |
| Cumulative average | Reference | 0.26 / 0.26 / 0.27 | (reference) | | | 0 |

**Table S8** Results of every forecasting model and simple reference for abundance (log(CPUE + 1) of eight taxa, CPUE being catch per unit effort), by scale and sampling method: median over the series of each group (estuaries: 56 series per sampling method, 7 estuaries × 8 taxa; coastwide: 8), calculated on the 367 forecast dates up to December 2022. All models forecast the monthly departures from the seasonal curve, which was refitted at each forecast date using only the data up to that date. RMSE: root mean squared out-of-sample prediction error in log units, 1, 12 and 24 months ahead. Cumulative average: the seasonal curve plus the mean departure of all earlier months; 12-month average: the same with the last 12 months only. Skill: 1 minus the ratio of the mean squared prediction error to that of the cumulative average (0 = no better than the cumulative average). Base model: the model without the feature being tested (covariates or trend): SARIMA (automatic) for ARIMAX and the SARIMA and damped trend versions; the state-space model for the state-space models with covariates or a trend. Skill against the base model measures what that feature adds (0 = no gain). Better / worse: numbers of series whose bootstrap range of skill against the base model lies above / below zero, 1 and 12 months ahead. Failed forecasts: cases (series × forecast date × months ahead) lost because a model could not be fitted at a forecast date, with their percentage of the model's evaluated cases; these cases were left out of the comparison with the base model for both models. Models with covariates were run to 12 months (n/a at 24).

| Model | Role | RMSE 1 / 12 / 24 | Skill vs cumulative average 1 / 12 / 24 | Skill vs base model 1 / 12 / 24 | Better / worse than base model at 1; at 12 | Failed forecasts (% of evaluated) |
|---|---|---|---|---|---|---|
| **Estuaries, bag seine** | | | | | | |
| SARIMA (automatic) | Base model | 0.66 / 0.70 / 0.72 | 0.09 / 0.00 / -0.02 | | | 1,210 (0.25%) |
| State-space | Base model | 0.67 / 0.72 / 0.74 | 0.07 / -0.01 / -0.05 | | | 0 |
| ARIMAX (temperature, salinity) | Covariates | 0.66 / 0.70 / n/a | 0.09 / 0.00 / n/a | -0.01 / 0.00 / n/a | 3 / 1; 5 / 4 | 590 (0.24%) |
| State-space with covariates | Covariates | 0.67 / 0.72 / n/a | 0.06 / -0.02 / n/a | -0.01 / 0.00 / n/a | 3 / 3; 2 / 8 | 1 (<0.01%) |
| SARIMA, no trend | Trend test | 0.66 / 0.70 / 0.72 | 0.09 / 0.00 / -0.02 | 0.00 / 0.00 / 0.00 | 0 / 0; 1 / 0 | 1,258 (0.26%) |
| SARIMA, linear trend | Trend test | 0.66 / 0.70 / 0.73 | 0.09 / -0.02 / -0.03 | 0.00 / -0.01 / -0.01 | 0 / 2; 0 / 9 | 2,566 (0.52%) |
| Damped trend | Trend test | 0.68 / 0.72 / 0.74 | 0.06 / -0.02 / -0.05 | -0.03 / -0.03 / -0.04 | 0 / 12; 1 / 8 | 0 |
| State-space with trend | Trend test | 0.68 / 0.74 / 0.77 | 0.06 / -0.05 / -0.10 | -0.01 / -0.04 / -0.06 | 0 / 14; 0 / 36 | 0 |
| Last month's value (naive) | Reference | 0.80 / 0.90 / 0.92 | -0.32 / -0.59 / -0.70 | | | 0 |
| 12-month average | Reference | 0.69 / 0.73 / 0.75 | 0.04 / -0.06 / -0.10 | | | 0 |
| Cumulative average | Reference | 0.71 / 0.71 / 0.70 | (reference) | | | 0 |
| **Estuaries, bay trawl** | | | | | | |
| SARIMA (automatic) | Base model | 0.65 / 0.73 / 0.75 | 0.26 / 0.02 / -0.01 | | | 552 (0.11%) |
| State-space | Base model | 0.67 / 0.76 / 0.78 | 0.23 / -0.04 / -0.10 | | | 0 |
| ARIMAX (temperature, salinity) | Covariates | 0.65 / 0.73 / n/a | 0.26 / 0.01 / n/a | -0.01 / 0.00 / n/a | 5 / 7; 0 / 1 | 85 (0.03%) |
| State-space with covariates | Covariates | 0.67 / 0.76 / n/a | 0.23 / -0.05 / n/a | -0.01 / -0.01 / n/a | 4 / 8; 0 / 13 | 0 |
| SARIMA, no trend | Trend test | 0.65 / 0.73 / 0.74 | 0.26 / 0.02 / -0.01 | 0.00 / 0.00 / 0.00 | 0 / 1; 1 / 0 | 624 (0.13%) |
| SARIMA, linear trend | Trend test | 0.66 / 0.74 / 0.76 | 0.25 / -0.01 / -0.05 | -0.01 / -0.03 / -0.04 | 1 / 4; 0 / 11 | 792 (0.16%) |
| Damped trend | Trend test | 0.67 / 0.76 / 0.78 | 0.22 / -0.03 / -0.09 | -0.06 / -0.06 / -0.08 | 0 / 31; 0 / 22 | 0 |
| State-space with trend | Trend test | 0.67 / 0.77 / 0.84 | 0.22 / -0.12 / -0.26 | 0.00 / -0.04 / -0.09 | 0 / 12; 0 / 46 | 0 |
| Last month's value (naive) | Reference | 0.74 / 0.93 / 0.95 | 0.02 / -0.51 / -0.63 | | | 0 |
| 12-month average | Reference | 0.70 / 0.74 / 0.78 | 0.11 / -0.05 / -0.07 | | | 0 |
| Cumulative average | Reference | 0.74 / 0.75 / 0.74 | (reference) | | | 0 |
| **Coastwide, bag seine** | | | | | | |
| SARIMA (automatic) | Base model | 0.46 / 0.54 / 0.56 | 0.24 / 0.01 / -0.02 | | | 72 (0.10%) |
| State-space | Base model | 0.49 / 0.56 / 0.58 | 0.15 / -0.03 / -0.11 | | | 0 |
| ARIMAX (temperature, salinity) | Covariates | 0.47 / 0.53 / n/a | 0.25 / 0.02 / n/a | 0.00 / 0.01 / n/a | 1 / 0; 1 / 0 | 25 (0.07%) |
| State-space with covariates | Covariates | 0.49 / 0.56 / n/a | 0.17 / -0.01 / n/a | 0.00 / 0.00 / n/a | 1 / 0; 1 / 1 | 0 |

| Model | Role | RMSE 1 / 12 / 24 | Skill vs cumulative average 1 / 12 / 24 | Skill vs base model 1 / 12 / 24 | Better / worse than base model at 1; at 12 | Failed forecasts (% of evaluated) |
|---|---|---|---|---|---|---|
| SARIMA, no trend | Trend test | 0.46 / 0.54 / 0.56 | 0.24 / 0.01 / -0.02 | 0.00 / 0.00 / 0.00 | 0 / 0; 0 / 0 | 72 (0.10%) |
| SARIMA, linear trend | Trend test | 0.46 / 0.53 / 0.55 | 0.24 / 0.02 / -0.03 | 0.00 / 0.00 / 0.00 | 1 / 0; 0 / 0 | 234 (0.33%) |
| Damped trend | Trend test | 0.49 / 0.56 / 0.58 | 0.14 / -0.02 / -0.12 | -0.13 / -0.04 / -0.07 | 0 / 6; 0 / 1 | 0 |
| State-space with trend | Trend test | 0.49 / 0.57 / 0.59 | 0.14 / -0.05 / -0.18 | 0.00 / -0.03 / -0.06 | 0 / 1; 0 / 6 | 24 (0.03%) |
| Last month's value (naive) | Reference | 0.52 / 0.64 / 0.70 | -0.02 / -0.37 / -0.57 | | | 0 |
| 12-month average | Reference | 0.51 / 0.56 / 0.58 | 0.09 / -0.06 / -0.17 | | | 0 |
| Cumulative average | Reference | 0.54 / 0.54 / 0.55 | (reference) | | | 0 |
| **Coastwide, bay trawl** | | | | | | |
| SARIMA (automatic) | Base model | 0.45 / 0.53 / 0.57 | 0.41 / 0.04 / -0.01 | | | 0 |
| State-space | Base model | 0.46 / 0.64 / 0.64 | 0.35 / -0.21 / -0.24 | | | 0 |
| ARIMAX (temperature, salinity) | Covariates | 0.46 / 0.53 / n/a | 0.41 / 0.06 / n/a | 0.01 / 0.00 / n/a | 0 / 0; 2 / 0 | 13 (0.04%) |
| State-space with covariates | Covariates | 0.46 / 0.63 / n/a | 0.36 / -0.20 / n/a | 0.00 / -0.01 / n/a | 1 / 1; 2 / 0 | 0 |
| SARIMA, no trend | Trend test | 0.45 / 0.53 / 0.57 | 0.41 / 0.04 / -0.01 | 0.00 / 0.00 / 0.00 | 0 / 0; 0 / 0 | 0 |
| SARIMA, linear trend | Trend test | 0.46 / 0.56 / 0.59 | 0.40 / -0.02 / -0.05 | -0.01 / -0.02 / -0.05 | 0 / 0; 0 / 1 | 46 (0.07%) |
| Damped trend | Trend test | 0.46 / 0.64 / 0.64 | 0.35 / -0.21 / -0.23 | -0.07 / -0.27 / -0.25 | 0 / 3; 1 / 4 | 0 |
| State-space with trend | Trend test | 0.47 / 0.65 / 0.67 | 0.36 / -0.25 / -0.35 | 0.00 / -0.04 / -0.09 | 0 / 1; 0 / 6 | 0 |
| Last month's value (naive) | Reference | 0.51 / 0.67 / 0.69 | 0.29 / -0.45 / -0.59 | | | 0 |
| 12-month average | Reference | 0.52 / 0.58 / 0.60 | 0.15 / -0.12 / -0.14 | | | 0 |
| Cumulative average | Reference | 0.58 / 0.58 / 0.58 | (reference) | | | 0 |

**Table S9** Trends in water temperature and salinity from 1982 to 2024 in each estuary and sampling method and coastwide. Mann-Kendall test and Sen slope on the yearly means of the departures from the seasonal curve (43 years in every series, each with at least six sampled months).

| | | Water temperature | | | | Salinity | | | |
|---|---|---|---|---|---|---|---|---|---|
| **Estuary** | **Sampling method** | **Tau** | **Sen slope (°C per decade)** | **p** | **Adjusted p** | **Tau** | **Sen slope (per decade)** | **p** | **Adjusted p** |
| Galveston Bay | Bag seine | 0.13 | 0.11 | 0.205 | 0.205 | 0.14 | 0.64 | 0.184 | 0.643 |
| | Bay trawl | 0.40 | 0.29 | < 0.001 | < 0.001 | 0.07 | 0.24 | 0.483 | 0.810 |
| Matagorda Bay | Bag seine | 0.45 | 0.39 | < 0.001 | < 0.001 | 0.09 | 0.45 | 0.402 | 0.733 |
| | Bay trawl | 0.37 | 0.26 | < 0.001 | < 0.001 | -0.04 | -0.17 | 0.714 | 0.810 |
| San Antonio Bay | Bag seine | 0.58 | 0.64 | < 0.001 | < 0.001 | 0.24 | 1.49 | 0.024 | 0.171 |
| | Bay trawl | 0.34 | 0.27 | 0.001 | 0.001 | 0.20 | 1.43 | 0.064 | 0.448 |
| Aransas Bay | Bag seine | 0.15 | 0.13 | 0.146 | 0.170 | 0.08 | 0.70 | 0.470 | 0.733 |
| | Bay trawl | 0.37 | 0.30 | < 0.001 | < 0.001 | 0.03 | 0.14 | 0.810 | 0.810 |
| Corpus Christi Bay | Bag seine | 0.30 | 0.26 | 0.005 | 0.007 | 0.05 | 0.15 | 0.653 | 0.761 |
| | Bay trawl | 0.42 | 0.38 | < 0.001 | < 0.001 | 0.04 | 0.13 | 0.683 | 0.810 |
| Upper Laguna Madre | Bag seine | 0.41 | 0.36 | < 0.001 | < 0.001 | 0.07 | 0.52 | 0.523 | 0.733 |
| | Bay trawl | 0.36 | 0.33 | < 0.001 | < 0.001 | 0.05 | 0.51 | 0.653 | 0.810 |
| Lower Laguna Madre | Bag seine | 0.42 | 0.41 | < 0.001 | < 0.001 | 0.00 | -0.01 | 0.992 | 0.992 |
| | Bay trawl | 0.38 | 0.38 | < 0.001 | < 0.001 | -0.03 | -0.07 | 0.794 | 0.810 |
| Coastwide | Bag seine | 0.40 | 0.33 | < 0.001 | | 0.11 | 0.55 | 0.320 | |
| | Bay trawl | 0.43 | 0.30 | < 0.001 | | 0.06 | 0.15 | 0.579 | |

Tau: Kendall's tau of the Mann-Kendall test. Adjusted p: p adjusted for multiple tests over the seven estuaries within each variable and sampling method (Benjamini and Hochberg 1995). The coastwide series are single tests and have no adjusted p. Salinity has no unit. Coastwide: effort-weighted monthly means over all estuaries (Online Resource 1, Sect. 1.3).
