## Supplementary material for "Forecasting estuarine biodiversity and population abundance for conservation: spatial scale, forecast horizon, and choice of measure": Online Resource 3: Supplementary figures

Biodiversity and Conservation

Masami Fujiwara

Department of Ecology and Conservation Biology, Texas A&M University, College Station, TX 77843-2258, USA

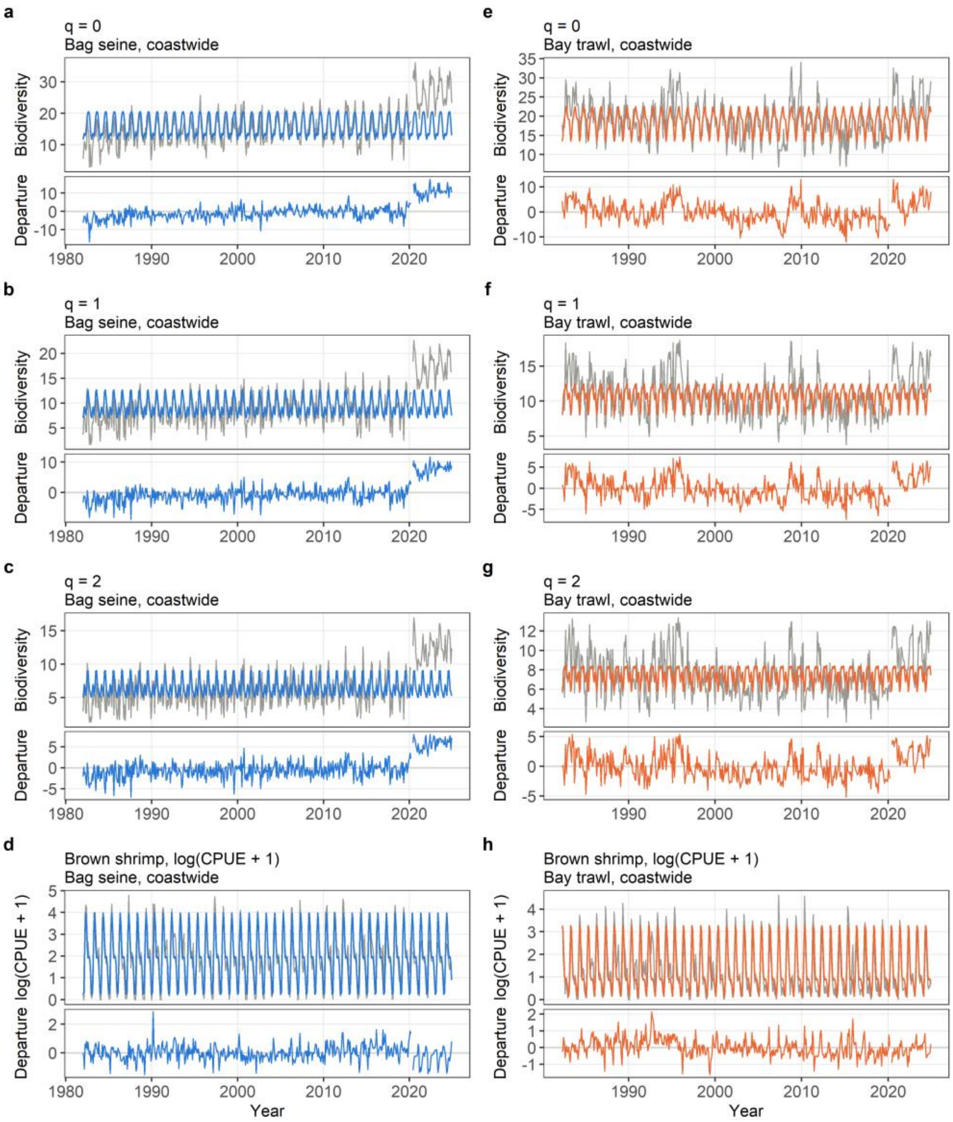


**Fig. S1** Seasonal curves and the departures from them for eight example series. Each panel: top, observed monthly values (gray) and the seasonal curve (a harmonic regression with periods of 12, 6, and 4 months) fitted to the whole record (color); bottom, the departures, which the models forecast. Left: bag seine; right: bay trawl; all coastwide (the fits for the single estuaries are summarized in Table S4). (a to c, e to g) Biodiversity, q = 0, 1, 2; (d, h) brown shrimp, log(CPUE + 1), an abundant target taxon caught by both sampling methods with a strong seasonal cycle, as an example of abundance. The forecasts used a curve refitted at each forecast date on earlier months only; the whole-record curve is shown for description. April and May 2020 (no sampling) appear as a gap

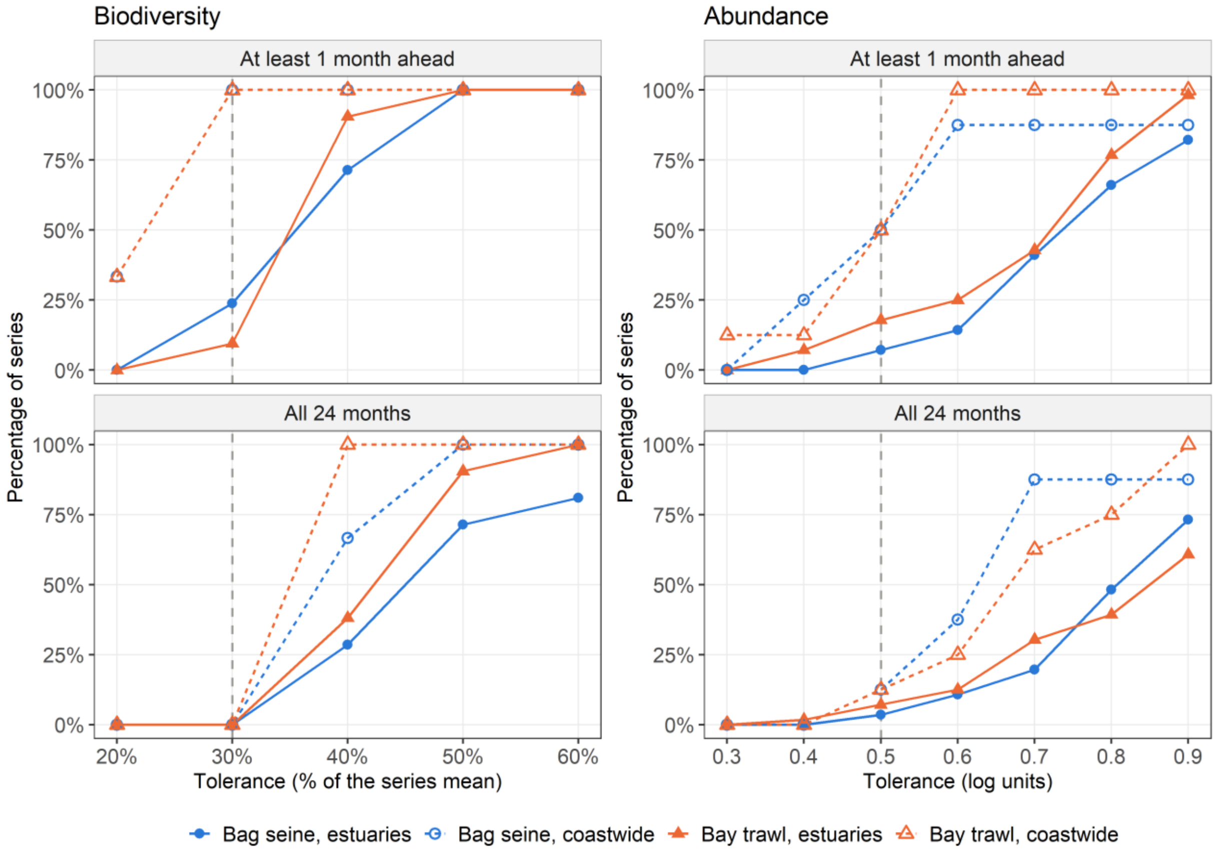


**Fig. S2** Percentage of series whose SARIMA forecasts stay within the tolerance, for tolerances from 20% to 60% of the series mean (biodiversity) and 0.3 to 0.9 log units (abundance): (top) at least one month ahead; (bottom) at every month up to 24 months ahead. Within the tolerance: the whole 95% uncertainty range of the RMSE of the out-of-sample prediction errors lies inside the tolerance (moving-block bootstrap over forecast dates, blocks of 12 months, 2,000 resamples). Dashed vertical line: the tolerance used in the other figures and tables (30%; 0.5 log units). Bag seine: blue with circles; bay trawl: orange with triangles; estuaries: solid lines and filled symbols; coastwide: dashed lines and open symbols (estuaries: 21 biodiversity and 56 abundance series per sampling method; coastwide: 3 and 8, so the coastwide biodiversity curves move in steps of 33%). Calculated on the 367 forecast dates up to December 2022. Because forecast error grew little with lead time (Fig. 3), the bottom curves follow the top ones

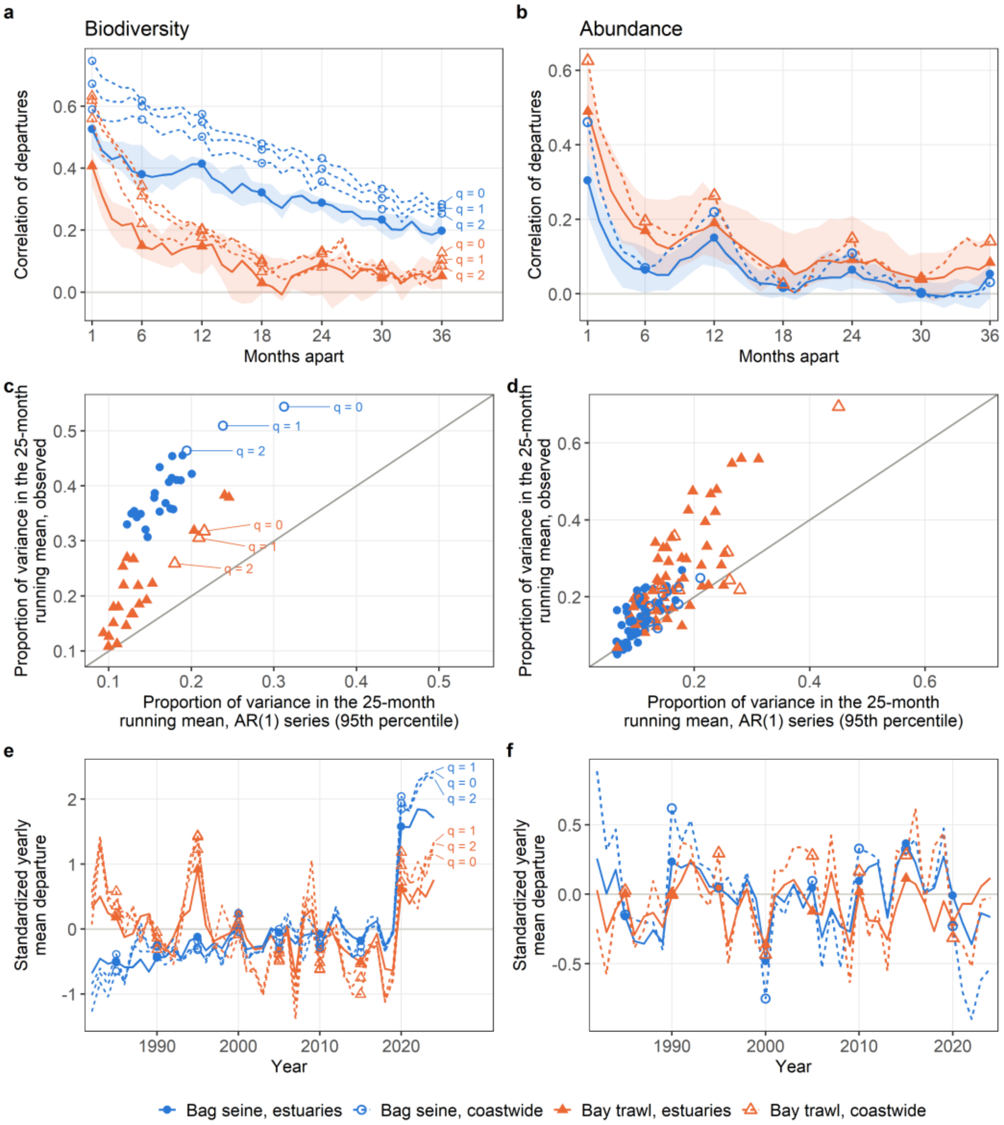


**Fig. S3** The departures from the seasonal curve over time, for biodiversity (left: Hill numbers q = 0, 1, 2) and abundance (right: log(CPUE + 1) of eight taxa, CPUE being catch per unit effort). Departures are each month's value minus the seasonal curve fitted to the whole record; April and May 2020 (not sampled) are left out. Bag seine: blue with circles; bay trawl: orange with triangles; estuaries: solid lines and filled symbols; coastwide: dashed lines and open symbols (estuaries: 21 biodiversity and 56 abundance series per sampling method; coastwide: 3 and 8). (a, b) Correlation of each series' departures with its own departures 1 to 36 months later; lines: median over the series of each group, except coastwide biodiversity, whose three orders are drawn separately and labeled; shaded: middle 50% of the estuary series. (c, d) One point per series: the proportion of the variance of the departures that lies in their 25-month running mean (slow change), against the 95th percentile of the same proportion in 1,000 simulated first-order autoregressive series with the series' own lag-1 correlation. Points above the line carry more slow change than their lag-1 correlation alone would give: all 48 biodiversity series and 108 of the 128 abundance series (95 of 112 in the estuaries, 13 of 16 coastwide). (e, f) Standardized yearly mean departure: for each series, the mean of its monthly departures in each year (at least six sampled months), divided by the standard deviation of its monthly departures so that series in different units can be combined; lines: median over the series of each group (coastwide biodiversity: each order)

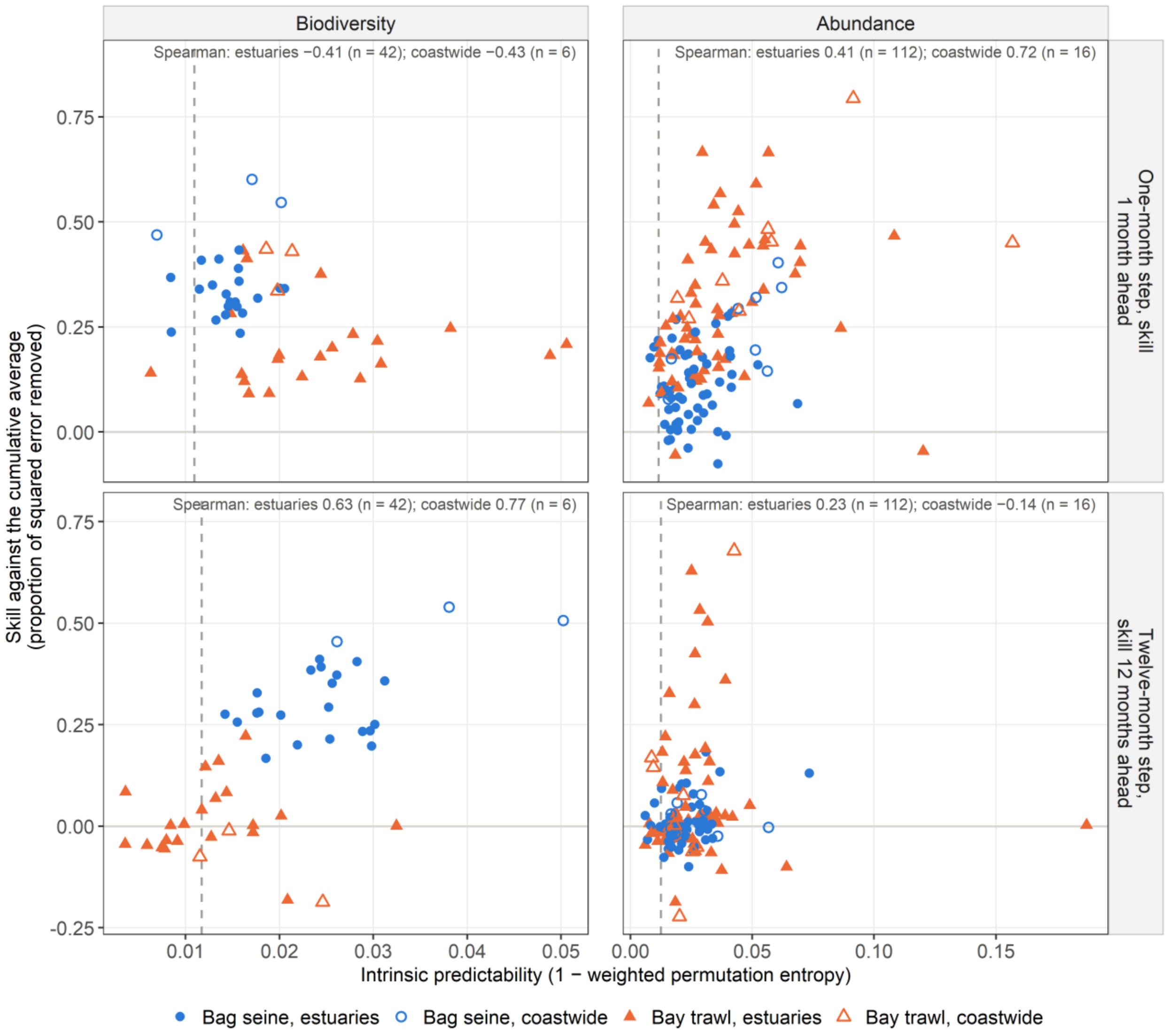


**Fig. S4** Weighted permutation entropy, expressed as 1 minus the entropy (intrinsic predictability), against the skill of SARIMA forecasts against the cumulative average, one point per series. The entropy is calculated from the order of 4 values of the departures from the seasonal curve: (top) 4 consecutive months, set against skill 1 month ahead; (bottom) the same calendar month in 4 consecutive years, set against skill 12 months ahead. Intrinsic predictability runs from 0 (no order) to 1 (fully ordered); dashed line: the median for shuffled copies of the series. Bag seine: blue circles; bay trawl: orange triangles; estuaries: filled; coastwide: open (estuaries: 21 biodiversity and 56 abundance series per sampling method; coastwide: 3 and 8). Rank correlations (Spearman) in each panel, with the number of series. Skill calculated on the 367 forecast dates up to December 2022; the axes differ between panels. Nine in ten series lie below 0.06 (shuffled copies about 0.01), so the departures carry little ordinal structure. Without the 24 estuary abundance series with many runs of zero catches, the estuary correlations for abundance are 0.54 (one-month step) and 0.31 (twelve-month step). Permutation entropy describes how ordered a series is; it does not separate nonlinear dynamics from random variation

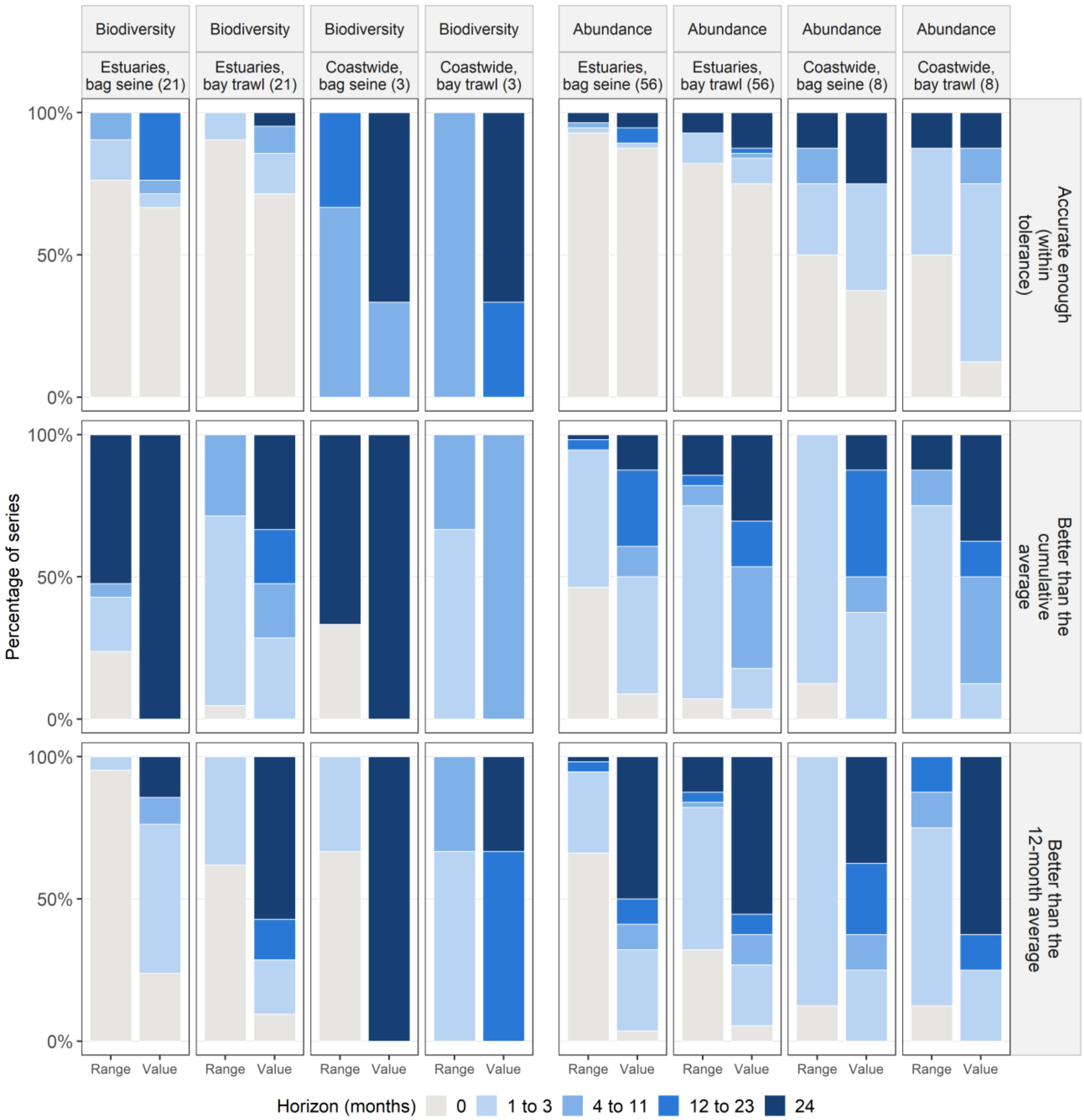


**Fig. S5** Forecast horizons of SARIMA forecasts: the number of consecutive months ahead, from one month ahead, for which the forecasts (top) stay within the tolerance, (middle) are better than the cumulative average, and (bottom) are better than the 12-month average. Tolerance: RMSE of the out-of-sample prediction errors within 30% of the series mean (biodiversity) or 0.5 log units (abundance). Both references are the seasonal curve plus the mean departure of all earlier months (cumulative average) or of the last 12 months (12-month average). Bars: percentage of the series of each group in each horizon class (0 = fails one month ahead; 24 = passes at every month); number of series in brackets. Range: the whole 95% uncertainty range must meet the condition (moving-block bootstrap over forecast dates, blocks of 12 months, 2,000 resamples); Value: the central estimate alone. A horizon ends at the first month that fails; skill that is lost and later returns is not counted (9 estuary biodiversity and 11 estuary abundance series; 1 and 2 coastwide). Calculated on the 367 forecast dates up to December 2022

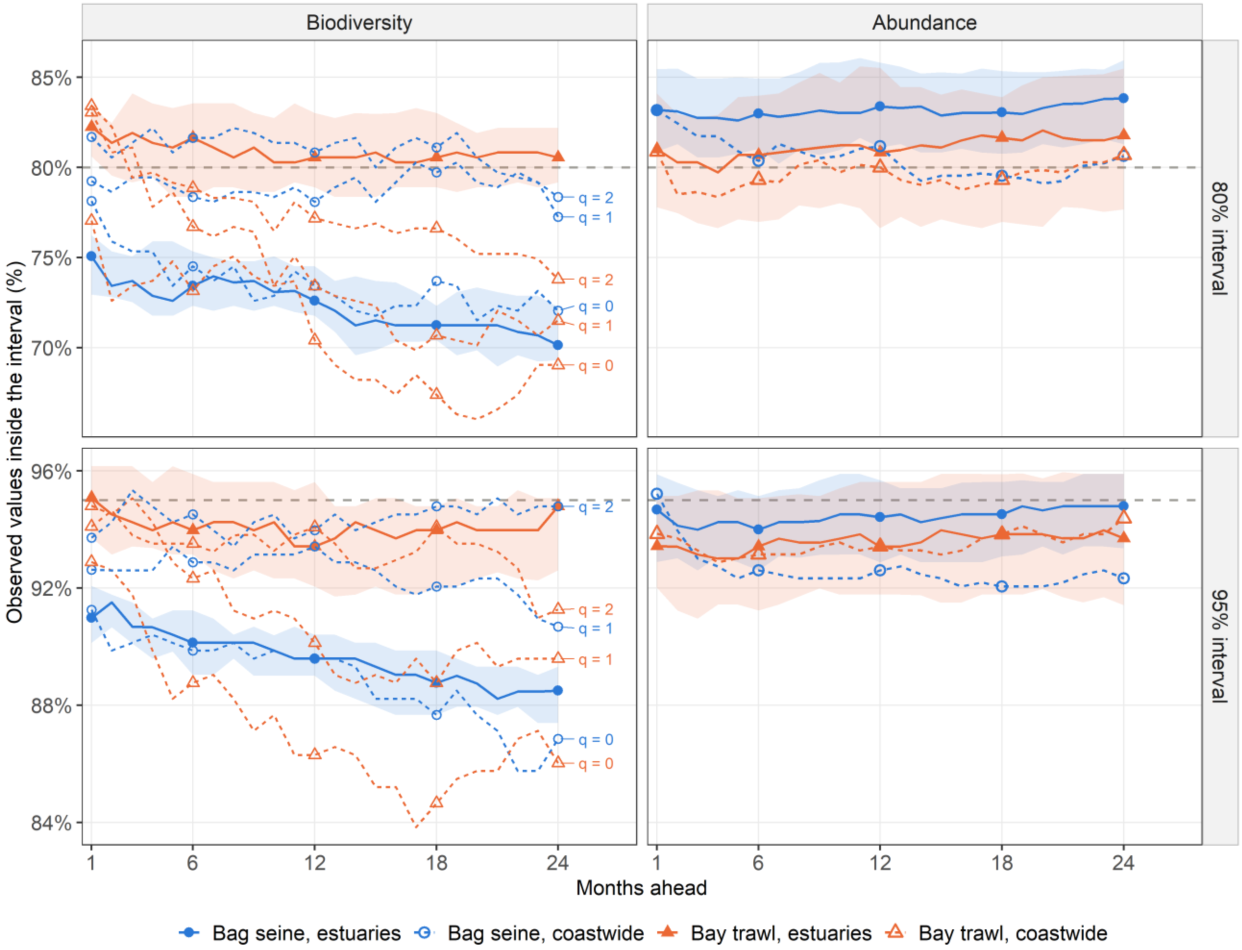


**Fig. S6** Percentage of observed values inside the 80% (top) and 95% (bottom) prediction intervals of SARIMA forecasts, by months ahead. Lines: median over the series of each group (estuaries: 21 biodiversity and 56 abundance series per sampling method; coastwide abundance: 8), except coastwide biodiversity, whose three orders are drawn separately and labeled; shaded: middle 50% of the estuary series; dashed gray line: the nominal level. Bag seine: blue with circles; bay trawl: orange with triangles; estuaries: solid lines and filled symbols; coastwide: dashed lines and open symbols. Calculated on the 367 forecast dates up to December 2022; the vertical scales differ between panels. The 80% intervals contained 70% to 75% of the observed values for bag seine biodiversity in the estuaries, and the coverage of coastwide bay trawl biodiversity declined from 83% one month ahead to 70% to 72% at 16 to 24 months; the other groups had 78% to 84%

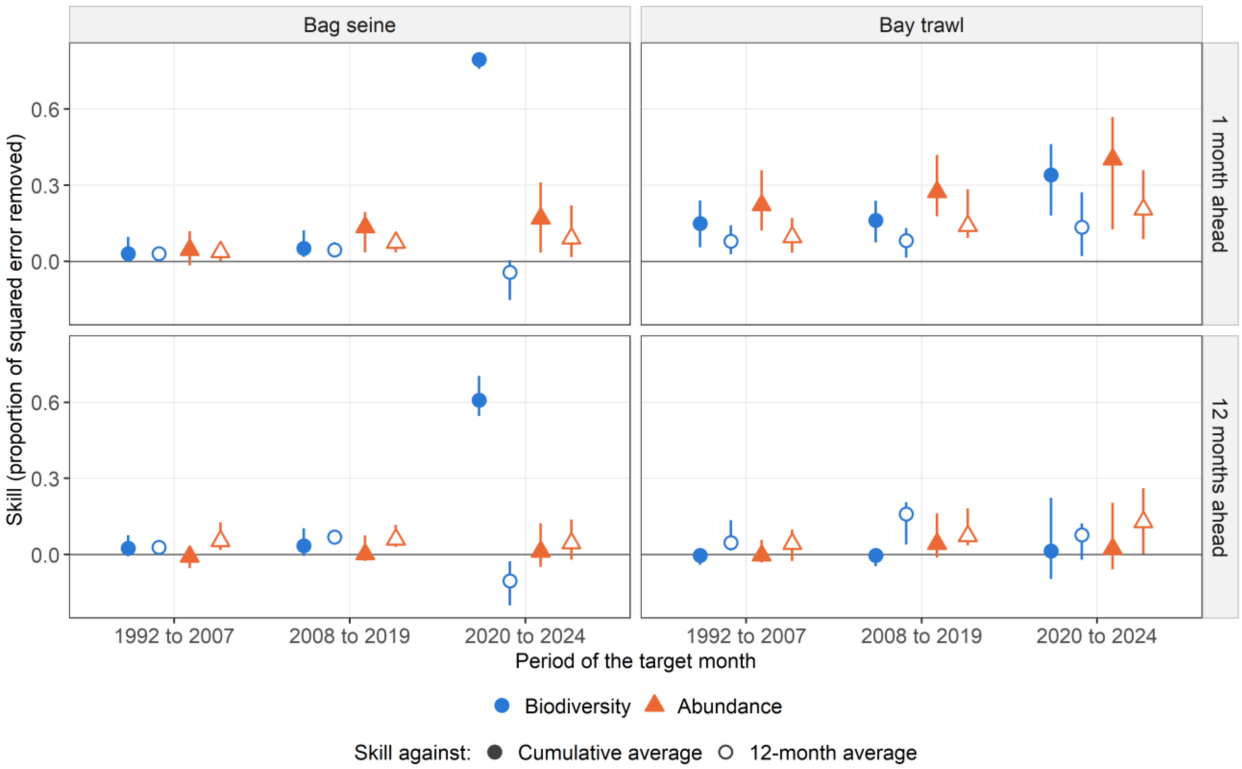


**Fig. S7** Skill of SARIMA forecasts for the estuary series by period of the target month, one and 12 months ahead, against the cumulative average (filled) and against the 12-month average (open). Biodiversity: blue circles; abundance: orange triangles. Median over the estuary series of each class and sampling method (21 biodiversity and 56 abundance series per sampling method), with the range of the middle 50%. Target months per period: 188, 144, and 57 one month ahead (177, 144, and 56 twelve months ahead); April and May 2020 were not sampled. Where skill is high against the cumulative average but near zero against the 12-month average, as for bag seine biodiversity in 2020 to 2024, the forecasts follow a level that had already shifted rather than anticipate the shift

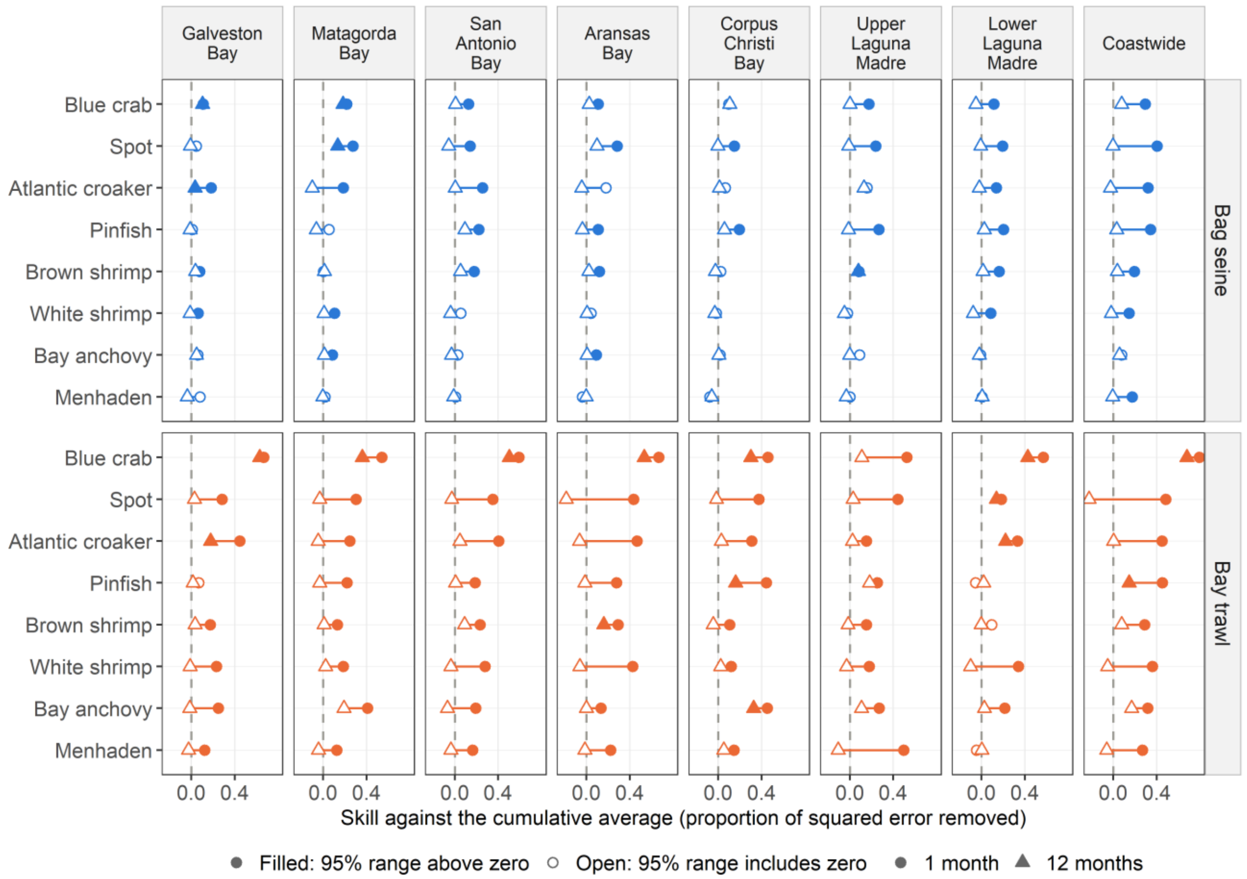


**Fig. S8** Skill of SARIMA abundance forecasts against the cumulative average for each of the eight taxa, by estuary (north to south) and coastwide, for each sampling method, 1 month (circles) and 12 months (triangles) ahead. Skill: proportion of the cumulative average's squared error that the forecast removes (0 = no better); cumulative average: the seasonal curve plus the mean departure of all earlier months. Filled: the 95% uncertainty range lies above zero (moving-block bootstrap over forecast dates); open: it includes zero. The line joins the two values of a series; taxa are ordered by their median one-month skill. Scientific names are in Table S3; menhaden is the genus *Brevoortia*. Calculated on the 367 forecast dates up to December 2022. One month ahead, bay trawl forecasts had higher skill than bag seine forecasts for the same taxon and estuary in 49 of 56 comparisons. By 12 months, the whole 95% uncertainty range of the skill was above zero in only 5 of the 56 bag seine series and 12 of the 56 bay trawl series; 8 of these 17 were blue crab
